# Multipartite coevolution shapes plant apoplastic immunity against rice blast fungus

**DOI:** 10.1101/2025.07.03.663104

**Authors:** Takumi Takeda, Motoki Shimizu, Atsushi Kodan, Hiroe Utsushi, Eiko Kanzaki, Satoshi Natsume, Tomoya Imai, Kaori Oikawa, Akira Abe, Ryohei Terauchi

## Abstract

β-1,3-glucan-binding proteins (GBPs) play crucial roles in cell wall integrity, host– pathogen interactions, and stress responses in fungi, making them essential for fungal survival and virulence. Here, we show that a GBP of the causal agent for blast fungus, *Magnaporthe oryzae*, is involved in multipartite coevolutionary arms race with rice (*Oryza sativa*). The rice thaumatin-like pathogenesis-related protein 5 (OsPR5) binds to *M. oryzae* GBP and sequesters it from the fungal cell wall to confer immunity, but this interaction is counteracted by thaumatin-binding proteins secreted by the fungus to suppress apoplastic immunity. Moreover, a rice receptor kinase located at the cell surface has evolved a thaumatin-like domain to detect the fungal GBP and activate immunity. Our findings reveal that GBPs and thaumatin-domain-containing proteins are engaged in intricate coevolutionary dynamics in the rice apoplast. Given the conservation of thaumatin across plants, we propose a general defense mechanism against fungal pathogens.

## INTRODUCTION

Plants are continually attacked by a plethora of pathogens, among which fungal pathogens are the major agents causing diseases of economic importance. The apoplast, the outer cellular space of fungal pathogens and plant hosts consisting of fluid-filled cell walls, is the major battleground of these interacting organisms. Fungal cell wall β-1,3-glucan contributes to a large proportion of polysaccharides in the fungal apoplast.^1^ Whereas chitin forms the inner layer of the fungal cell wall, β-1,3-glucan resides in the outer layer of this structure, where it is presumably involved in various interactions. Glucan-binding proteins (GBPs), which bind to β-1,3-glucan, are distributed in a wide variety of organisms, including animals, insects, fungi, and bacteria.^2–8^ However, the extent to which fungal GBPs and their plant interactors mediate apoplastic battles in fungal–plant pathosystems remains largely unknown.

Thaumatin-like proteins (TLPs) are found in insects, fungi, and plants and are thought to have various biological functions. TLPs in the green lineage usually comprise a secretion signal peptide and a thaumatin domain (ThD, Pfam PF00314) and are widely distributed from algae to angiosperms.^9–11^ Many studies have shown that TLPs participate in plant responses to biotic and abiotic stresses.^9,12^ The expression of genes encoding TLPs, also known as *PATHOGENESIS-RELATED 5* (*PR5*) genes is induced by pathogen infection and is commonly used as a marker of plant responses to pathogen infection. Overexpressing individual *TLP* genes confers resistance against fungal pathogens in both dicot and monocot species.^13–18^ The direct application of recombinant purified TLPs to various pathogens resulted in diminished fungal growth,^19–23^ suggesting that TLPs have antifungal activity, although the underlying mechanism remains elusive.

Plant pathogens employ countermeasures against plant-derived TLPs. Alta1, an apoplastic effector from the fungus *Alternaria alternata*, binds to and inhibits the β-1,3- glucanase activity of kiwi (*Actinidia deliciosa*) PR5.^24^ Similarly, PevD1 from *Verticillium dahlia* binds to cotton (*Gossypium hirsutum*) PR5 (GhPR5) and inhibits its antifungal activity.^25^ Unveiling the significance of the actions of TLPs, their inhibiting proteins, and their interactions is important for understanding the mechanisms of pathogen infection and plant defense.

In this study, we determined that a GBP from the blast fungus *Magnaporthe oryzae* is the target of rice (*Oryza sativa*) PR5 (OsPR5). We also show that a pair of *M. oryzae* TLP- binding proteins (ThBPs) inhibit the OsPR5–GBP interaction as a countermeasure against plant defenses. We identified the crucial amino acids responsible for the interactions of OsPR5 with GBP and ThBPs, suggesting that these proteins are involved in a multipartite arms race. We propose a trajectory of plant TLP evolution based on phylogenetic analysis and binding analysis. Finally, we demonstrate that GBP is recognized by a rice cell surface–localized receptor thaumatin kinase (OsThK1) that transduces the defense signal.

## RESULTS

### Rice TLPs interact with apoplastic fungal GBP and thaumatin-binding proteins

Our laboratory studies host–pathogen interactions in the plant extracellular (apoplastic) space using rice and blast fungus (*M. oryzae*) as a model system. Thaumatin-like proteins (TLPs) are secreted proteins known to be involved in plant defense. However, their function have not been elucidated. To explore the function of TLPs and their effect on pathogens, we confirmed that knockdown rice lines for the TLP gene *OsPR5* were more susceptible to *M. oryzae* (Ken 53-33) than the wild type (WT, cultivar Moukoto; Figure S1). To identify *M. oryzae* proteins that bind to TLP, we performed an *in vitro* pull-down experiment using OsPR5 tagged with a His epitope (OsPR5-His). We mixed recombinant purified OsPR5-His with secreted proteins from *M. oryzae*, incubated them with His-tag affinity resin (His-resin), and analyzed the fraction bound to the His-resin by SDS-PAGE and silver staining. Three protein bands were visible after pulling down the mixture of *M. oryzae* proteins and OsPR5-His, but not from *M. oryzae* proteins alone (Figure 1A). Liquid chromatography–tandem mass spectrometry (LC-MS/MS) of the three bands revealed a putative *M. oryzae* glucan-binding protein (GBP, MGG_05232) and two functionally uncharacterized proteins, which we named thaumatin-binding protein 24 kDa (ThBP24k, MGG_01944) and thaumatin-binding protein 15 kDa (ThBP15k, MGG_10456). GBP contains a secretion signal peptide and a carbohydrate-binding module family 52 (CBM52, PF10645). ThBP24k and ThBP15k contain secretion signal peptides as well but no other known domains. ThBP24k is presumed to be glycosylated, as its apparent molecular weight determined by SDS-PAGE (∼30 kDa) is higher than the theoretical weight calculated from the amino acid sequence (23.8 kDa). We analyzed the binding kinetics of OsPR5 to each of the three *M. oryzae* proteins (Figure S2). The integrated intensity of GBP, ThBP24k, and ThBP15k rapidly rose following the addition of OsPR5, resulting in stable values. The binding kinetics (*K*_D_) values of OsPR5 to GBP, ThBP24k, and ThBP15k were 7.45×10^−10^ M (± 9.06×10^−11^), 2.41×10^−9^ (± 2.42×10^−10^), and 2.59×10^−8^ M (± 3.68×10^−9^), respectively (Figure S2), suggesting that these protein complexes form rapidly and are stable. The TLPs encoded by Os12g0630100, Os03g0661600, Os03g0663500 and Os12g0568900 highly accumulated in rice leaves infected with *M. oryzae* (Table S1), and all except Os12g0568900 bound to GBP and ThBPs in a pull-down assay (Figure S3), suggesting that multiple TLPs can interact with GBP and ThBPs.

**Figure 1.**
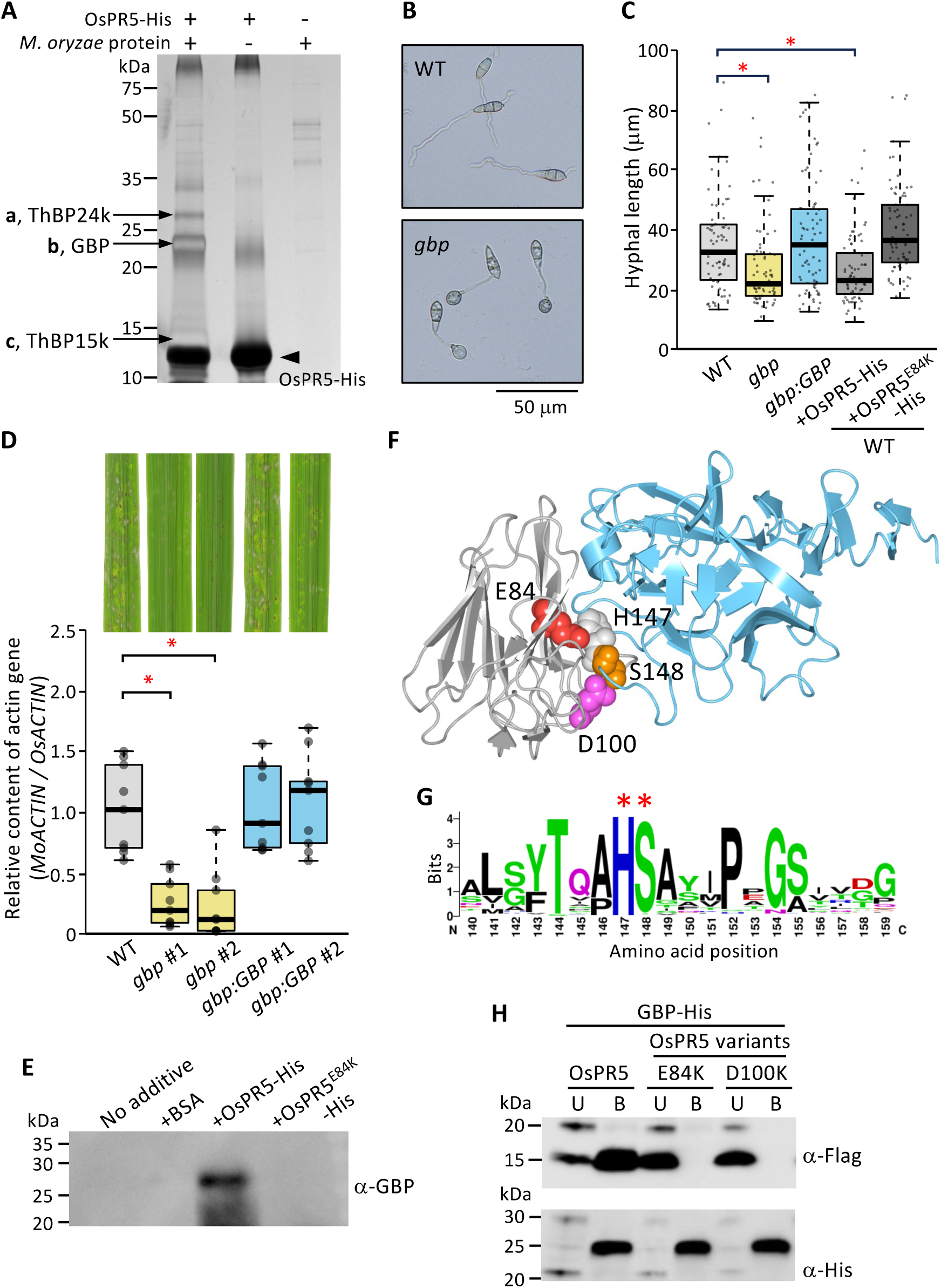
*M. oryzae* GBP is important for efficient infection of the host plant. **(A)** Proteins prepared from an *M. oryzae* (compatible strain Ken53-33) culture filtrate were incubated with recombinant purified OsPR5-His and His-resin. *M. oryzae* proteins and OsPR5-His were individually used as controls. Fractions bound to His-resin were subjected to SDS-PAGE, followed by silver staining. The arrowhead indicates OsPR5-His. The protein bands indicated by arrows (a–c) were analyzed by liquid chromatography–tandem mass spectrometry (LC–MS/MS). **(B)** Representative photograph of germinated hyphae from *M. oryzae* WT and *gbp* conidia. **(C)** Hyphal lengths of *M. oryzae* WT, *gbp*, *gbp*:*GBP*, and WT treated with recombinant purified OsPR5-His or OsPR5^E84K^-His (*n* = 60). Conidia were incubated in 30 μL of 50 mM sodium phosphate buffer (pH 5.5) with 0.2% (w/v) glucose and 0.05% (w/v) yeast extract on a hydrophobic glass slide at 25°C for 7 h. OsPR5-His or OsPR5^E84K^-His (1 μg) was added at time 0. Asterisks indicate a significant difference at *P* < 0.01 according to a two-sided Student’s *t*-test. **(D)** Infection test of *M. oryzae* WT, *gbp*, and *gbp*:*GBP* strains (*n* = 9 each) on rice leaves. The extent of *M. oryzae* infection was evaluated based on relative fungal mass by measuring the ratio of *M. oryzae Actin* genomic DNA (*MoACTIN*) to *O. sativa Actin* genomic DNA (*OsACTIN*) using qPCR. The average ΔΔCt value for WT was set to a ratio of 1. Data are shown as means ± SD from nine independent determinations. Asterisks indicate significant difference at *P* < 0.01 according to a two-sided Student’s *t*-test. Representative photographs of inoculated leaves are shown to the top. **(E)** Release of GBP from the cell wall, as analyzed using *M. oryzae* cultures in the presence of BSA, OsPR5-His, or OsPR5^E84K^-His (1 μg/0.1 mL each). After 8 h of culture, the *M. oryzae* filtrate was analyzed by immunoblotting using anti-GBP antibody. **(F)** Model of the OsPR5 and GBP 3D structures, constructed using AlphaFold3. Based on the complex models, amino acids with short inter-atomic distances between OsPR5 and GBP were extracted by Waals software, as indicated by balls (red; glutamic acid residue in OsPR5, magenta; aspartic acid residue in OsPR5, grey; histidine residue in GBP, orange; serine residue in GBP) in the model. Cyan ribbon model, GBP; Grey ribbon model, OsPR5. **(G)** Protein WebLogo of the amino- acid frequency in proteins homologous to GBP from 113 fungal species. Asterisks indicate the candidate amino acids that interact with E84 and D100 of OsPR5. **(H)** Pull-down assay with Flag-tagged intact OsPR5 or its variants with amino acid substitutions at E84 or D100 and GBP-His using His-resin. Proteins were separated into unbound (U) and bound (B) fractions to His-resin and detected by immunoblotting using anti-Flag and anti-His antibodies.

### GBP is involved in the growth of *M. oryzae* hyphae during infection

The GBP identified above as an OsPR5 interactor has not been characterized. To explore its roles in fungal development and host–pathogen interactions, we conducted biochemical and genetic studies. First, to investigate the cellular localization of GBP, we examined the culture filtrate and cell wall fractions from *M. oryzae* liquid culture by immunoblotting using an antibody against GBP. GBP was primarily present in the cell wall fraction, with a small amount in the culture filtrate (Figure S4A), suggesting GBP is associated with hyphal growth. In a binding assay to cell wall polysaccharides, GBP bound to an *M. oryzae* cell wall preparation and to crystalline- and amorphous-β-1,3- glucan (Figure S4B). *GBP* transcript levels were very low in conidia but markedly increased at 1 and 2 days after inoculation of rice leaves with the fungus, followed by a decline at 3 and 4 days after inoculation (Figure S4C). These results suggest that GBP specifically binds to β-1,3-glucan and might function in cell wall modification during growth of liquid culture and infection to rice plant.

To study the function of GBP, we generated *GBP* knockout (*ko*) mutant strains of *M. oryzae* (*gbp*) and a *GBP* complementation line in the *gbp* mutant background (*gbp:GBP*). Immunoblot analysis confirmed that the *gbp* mutants did not produce GBP, whereas *gbp:GBP* produced this protein to roughly WT levels (Figure S5). We measured cell wall hydrolysis in WT and *gbp M. oryzae* strains by incubating native cell wall fractions prepared from each of these lines with *Trichoderma* sp. endo-β-1,3-glucanase and measured the amount of hydrolyzed sugars (Figure S6). The amount of hydrolyzed sugars from native cell wall preparations was much lower in *gbp* compared to the WT strain, reaching only about 25% of WT levels, indicating that *gbp* cell walls are less susceptible to digestion by endo-β-1,3-glucanase.

We also examined hyphal growth and appressorium formation 7 h after conidium germination in WT and the *gbp* mutant on a hydrophobic glass slide (Figures 1B and S7). The germinated hyphae were significantly shorter in *gbp* compared to the WT. This phenotype returned to WT levels in the *ghp:GBP* strain (Figure 1C). Almost no appressorium formation occurred 7 h after germination in the WT, but more than 70% of conidia formed appressoria in the *gbp* mutant (Figure S8). These results suggest that GBP functions in hyphal growth by providing cell wall–modifying enzymes with access to β- 1,3-glucan in the hyphal cell walls and that GBP modulates appressorium development. We also studied the function of GBP in fungal virulence against rice plants using *gbp* mutants and *gbp:GBP* complementation strains of *M. oryzae* (Ken 53-33). Two g*bp* mutant strains (#1 and #2) caused less severe disease symptoms on rice leaves compared to the WT *M. oryzae* strain (Figures 1D and S9). In agreement with this observation, the relative biomass of the *gbp* mutants was significantly lower than that of the WT *M. oryzae* strain, as judged by quantitative PCR (qPCR) analysis of *ACTIN* genes from *M. oryzae* and rice. In the *gbp:GBP* complementation strains (#1 and #2), disease symptoms and the relative amount of fungal mass in the plants were comparable to those observed following infection with WT *M. oryzae*. These results indicate that *M. oryzae* GBP is involved in cell wall modification during hyphal growth.

### OsPR5 targets GBP and inhibits its function

Treating WT *M. oryzae* conidia with recombinant purified OsPR5-His slowed down hyphal growth (Figure 1C) and enhanced appressorium formation (Figure S8) 7 h after germination, which is a phenocopy of the *gbp* mutant. By contrast, treatment with the point mutant OsPR5^E84K^-His, which cannot bind to GBP (see below), did not affect hyphal growth (Figure 1C). When OsPR5-Flag was added to *M. oryza*e native cell walls, we detected OsPR5 in the cell wall–bound fraction for the WT and *gbp:GBP* strains, but not the *gbp* mutant (Figure S10A), confirming the specific binding of OsPR5 to GBP. Furthermore, GBP was released into the soluble fraction when *M. oryzae* hyphae was grown in the presence of OsPR5-His, but neither BSA nor OsPR5^E84K^-His (Figure 1E), suggesting that OsPR5 specifically binds to and sequesters GBP from fungal cell walls. Previous studies suggested that TLPs bind to purified, crystalline β-1,3-glucan,^26,27^ which we confirmed with six different OsTLPs (Figure S11). However, OsPR5 bound only weakly to protein-free *M. oryzae* native cell wall fractions, which contain amorphous β- 1,3-glucan (Figure S10B). These results suggest that the primary function of OsPR5 is to bind GBP and isolate it from β-1,3-glucan to confer immunity.

To visualize the protein–protein complexes formed by OsPR5 and *M. oryzae* GBP, we predicted their three-dimensional (3D) structures using AlphaFold3 (Figure 1F). In the structural models, the cleft region of OsPR5 is positioned close to *M. oryzae* GBP.

The prediction values obtained for the interface predicted template modeling (ipTM; 0.89) and predicted template modeling (pTM; 0.90) values (Figure S12) demonstrate the reliability of the predicted complex structures.^28^ We predicted 3D models for each of five TLPs (OsPR5, Os12g0630100, Os12g0630500, Os07g0417600, and At1g75040 [AtPR5]) in complex with GBP. We noticed a pair of amino acids in each of the two proteins in the complex with inter-atomic distances < 4Å, which we considered as candidate amino acids involved in the interactions (Data S1). Atoms in a glutamic acid residue (E84) and an aspartic acid residue (D100) of OsPR5 were positioned within short distances of atoms from the H147 and S148 residues of GBP, respectively (Figures 1F and S13). To evaluate the conservation of amino acids important for forming protein complexes, we examined the frequencies of amino acids in homologs of GBP (113 proteins) from a wide range of Ascomycete species (Figures 1G and S14A; Data S2). The amino acids H147 and S148 of GBP are highly conserved in the homologous proteins (H147 conserved in 113/113 proteins and S148 in 111/113 proteins), suggesting their functional significance.

To validate the importance of the interacting amino acid residues predicted above, we generated Flag-tagged OsPR5 variant proteins with amino acid substitutions to perform pull-down binding assays using His-tagged GBP and His-resin. When intact OsPR5-Flag and GBP-His were used in the binding assay, we detected OsPR5-Flag in the bound fraction (Figure 1H). However, when OsPR5^E84K^-Flag or OsPR5^D100K^-Flag was tested for binding to GBP-His, neither OsPR5-Flag variant was present in the bound fraction, indicating that their single amino acid substitutions abrogated binding to GBP. Amino acid substitutions at E84 or D100 to other amino acids also blocked the binding of these OsPR5-Flag variants to GBP (Figure S15). We also performed binding assays using variants of GBP against OsPR5 (Figure S16A). Amino acid substitutions at H147 or S148 of GBP abrogated the binding of these GBP variants to OsPR5, highlighting the functional importance of H147 and S148. Together, our biochemical analysis suggests that the predicted structural models accurately represent these protein complexes and point to the importance of the E84 and D100 residues of OsPR5 for interactions with the H147 and S148 residues of GBP.

### ThBPs suppress the binding of OsPR5 to GBP

ThBP24k and ThBP15k occurred mostly in the culture filtrate of *M. oryzae* liquid culture (Figure S4A), suggesting that the proteins are secreted to rice apoplast and subsequently bind OsPR5 during the infection of *M. oryzae* to rice. *ThBP24k* transcript levels were very low in conidia but markedly increased at 1 and 2 days after inoculation of rice leaves with the fungus, followed by a decline at 3 and 4 days after inoculation(Figure S4C). Notably, *ThBP24k* transcript levels followed the same pattern as those of *GBP*. *ThBP15k* expression was also induced at 1 day after inoculation but remained high. These results suggest that ThBPs may function in host invasion.

We hypothesized that ThBPs bind to OsPR5 and block its binding to GBP. To investigate whether ThBPs inhibit OsPR5 activity, we performed pull-down experiments to detect OsPR5-Flag and GBP-His following an incubation of OsPR5 with ThBPs (Figure 2A). We analyzed both the unbound and bound fractions to His-resin by immunoblotting using anti-OsPR5 and anti-His antibodies. When only OsPR5-Flag and GBP-His were mixed, we detected most of OsPR5-Flag in the bound fraction, indicating that a OsPR5-GBP complex had formed. When OsPR5-Flag was preincubated with ThBP15k or ThBP24k, however, followed by incubation with GBP-His, OsPR5-Flag was present in the unbound fraction, indicating that ThBPs interfered with the binding of OsPR5 to GBP.

**Figure 2.**
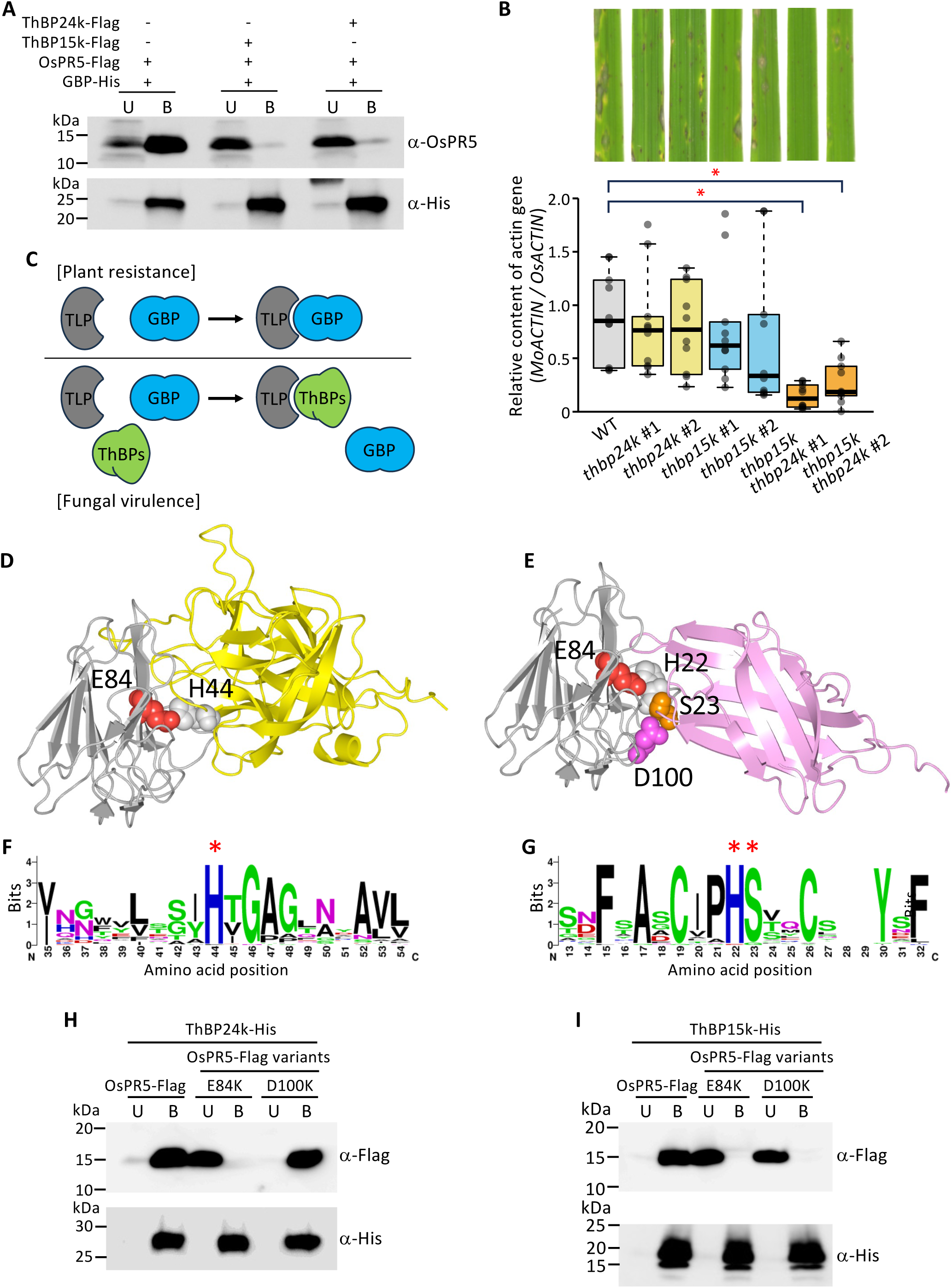
The *M. oryzae* proteins ThBP24k and ThBP15k act as counterdefense mechanisms against the action of plant TLPs. **(A)** Pull-down assay examining the inhibition of OsPR5–GBP complex formation by ThBPs. Protein extract containing OsPR5-Flag produced in *N. benthamiana* leaves was incubated with GBP-His following incubation with ThBP24k or ThBP15k as indicated, and His-resin. OsPR5- Flag and GBP-His were detected by immunoblotting using anti-OsPR5 and anti-His antibodies, respectively. **(B)** Infection test of *M. oryzae* WT, *ThBP24k* knockout (*thbp24k*) mutants, *ThBP15k* knockout (*thbp15k*) mutants and double knockout (*thbp15k thbp24k*) mutants (*n* = 10 each) on rice leaves. The relative amount of *M. oryzae* fungal mass was determined by measuring the ratio of *M. oryzae Actin* genomic DNA (*MoACTIN*) to *O. sativa Actin* genomic DNA (*OsACTIN*) using qPCR. The average ΔΔCt value for WT was set to a ratio of 1. Data are shown as means ± SD from ten independent determinations. Asterisks indicate significant differences at *P* < 0.01 according to a two-sided Student’s *t*-test. Representative photographs of inoculated leaves are shown to the top. **(C)** Diagram of protein–protein interactions between TLP and GBP, ThBP24k, and ThBP15k. **(D, E)** 3D-structural models of OsPR5 and ThBP24k **(D)** or OsPR5 and ThBP15k **(E)**, predicted using AlphaFold3. Based on the models, amino acids with short inter-atomic distances between OsPR5 and *M. oryzae* proteins were extracted by Waals software and are indicated by balls (red; glutamic acid residue in OsPR5, magenta; aspartic acid residue in OsPR5, grey histidine residue in ThBP24k and ThBP15k, orange; serine residue in ThBP15k) in the models. Yellow ribbon model, ThBP24k; Pink ribbon model, ThBP15k; Grey ribbon model, OsPR5. **(F, G)** WebLogos showing the frequency of amino acids in proteins homologous to ThBP24k **(F)** or ThBP15k **(G)** from 53 and 48 fungal species, respectively. Asterisks indicate the candidate amino acids that interact with E84 or D100 of OsPR5. **(H, I)** Pull-down assays with OsPR5-Flag or its variants with amino acid substitutions at E84 or D100 and ThBP24k-His **(H)** or ThBP15k-His **(I)** using His-resin. Proteins were separated into unbound (U) and bound (B) fractions to His-resin and detected by immunoblotting using anti-Flag and anti-His antibodies.

To examine the functions of ThBPs in fungal virulence against rice, we generated single and double knockout *M. oryzae* strains for these *ThBP* genes (*thbp24*, *thbp15k*, and *thbp15k thbp24k*). Immunoblot analysis confirmed that all mutant strains failed to produce the respective proteins (Figure S5). When rice was inoculated with *M. oryzae thbp24k* (#1 and #2) or *thbp15k* (#1 and #2), disease symptoms and relative fungal biomass appeared to be lower compared to inoculation with WT *M. oryzae*, although these differences did not reach statistical significance (Figures 2B and S9). By contrast, rice plants inoculated with the double knockout *thbp15k thbp24k* strains developed less severe disease symptoms and accumulated significantly less relative fungal biomass compared to the controls. These results suggest that rice TLPs target GBP to inhibit its function in cell wall modification during *M. oryzae* invasion; as a counterdefense, the fungus secretes ThBP24k and ThBP15k to inhibit the binding of TLPs to GBP (Figure 2C).

We predicted the 3D structures of the protein–protein complexes of TLPs to the *M. oryzae* proteins ThBP24k and ThBP15k using AlphaFold3, using each of the five TLPs OsPR5, Os12g0630100, Os12g0630500, Os07g0417600, and At1g75040 (AtPR5) complexed to ThBP24k or ThBP15k. The amino acids with the shortest inter-atomic distances between TLPs and ThBP24k were E84 in OsPR5 and H44 in ThBP24k; those between TLPs and ThBP15k were E84 and D100 in OsPR5 facing the histidine (H22) and serine (S23) residues of ThBP15k (Figures 2D, 2E, S12 and S13). We examined the frequencies of amino acid residues in homologs of ThBP24k (53 proteins) and ThBP15k (48 proteins) from a wide range of Ascomycete species (Figures 2F, 2G, S14B and S14C; Data S2). The amino acid residues H44 in ThBP24k, and H22 and S23 in ThBP15k, are highly conserved in the homologous proteins (H44 in ThBP24k: 52/53 proteins; H22 and S23 in ThBP15k: 47/48 proteins and 46/48 proteins, respectively, suggesting their functional significance.

To validate the importance of the predicted interacting amino acid residues, we performed pull-down binding assays of Flag-tagged OsPR5 variants to ThBPs. When OsPR5-Flag and ThBP24k-His were used in a pull-down binding assay, we detected OsPR5-Flag in the bound fraction, whereas OsPR5^E84K^-Flag was present in the unbound fraction (Figure 2H). As a control, we tested OsPR5^D100K^-Flag for its binding to ThBP24k- His, as D100 is not located near the OsPR5–ThBP24k binding interface; we detected OsPR5^D100K^-Flag in the bound fraction in a pull-down assay with ThBP24k-His. When OsPR5-Flag, OsPR5^E84K^-Flag, or OsPR5^D100K^-Flag was incubated with ThBP15k-His, we detected OsPR5-Flag in the bound fraction, whereas OsPR5^E84K^-Flag and OsPR5^D100K^- Flag were present in the unbound fraction (Figure 2I). These results indicate that the E84 and D100 residues of OsPR5 are important for its binding to GBP (Figure 1H) and that these residues are also bound by ThBP24k and ThBP15k, respectively. We also performed binding assays using variants of ThBP24k and ThBP15k against OsPR5 (Figure S16). Amino acid substitutions of H44 in ThBP24k and H22 and S23 in ThBP15 abrogated the binding of these proteins to OsPR5, highlighting the functional importance of these amino acids.

### Evolution of plant TLPs

TLPs are widely distributed in the green lineage from algae to angiosperms.^8–10^ We searched for TLPs using keywords *thaumatin and PF00314* in the Ensembl Plants database for the following species: *Galdieria sulphuraria*, Chlamydomonas (*Chlamydomonas reinhardtii*), *Marchantia polymorpha*, *Physcomitrium patens*, *Selaginella moellendorffii*, rice, maize (*Zea mays*), Arabidopsis, and black cottonwood (*Populus trichocarpa*). Additionally, we performed a BLAST search (E < 1e-4) using OsPR5 amino acid sequence as a query for *Alsophila spinulosa* using the Figshare database.

We grouped the obtained proteins based on their amino acids at the positions equivalent to E84 and D100 in OsPR5, both of which are important for binding to GBP (Figure 3A). We identified a TLP with an amino acid equivalent to E84 but without the D100 counterpart of OsPR5 in Chlamydomonas but not in *G. sulphuraria*, suggesting that this TLP may be the origin of green lineage TLPs. The total number of *TLP* genes per genome increased during evolution. Notably, TLPs with amino acid residues equivalent to E84 and D100 of OsPR5, which we named E-D type TLPs, first appeared in bryophytes (*M. polymorpha*, *P. patens*); their numbers increased in ferns (*A. spinulosa*) and angiosperm plants. We also identified TLPs in rice, maize, Arabidopsis, and *P. trichocarpa* with an equivalent residue to D100 but not E84 of OsPR5 (x-D type), as well as TLPs lacking an equivalent residue to E84 and D100 of OsPR5 (x-x type). However, the E-D type of TLPs accounted for the majority of (65-92%) TLPs in angiosperms. To determine whether the E-D type TLPs of *P. patens* binds to GBP, we performed a pull-down experiment using Flag-tagged *P. patens* TLPs and GBP-His with His-resin (Figure 3B). When Pp3c17_5160-Flag (E-D type) or Pp3c9_14830-Flag (E-x type) was used in the pull-down assay, we detected Pp3c17_5160-Flag and Pp3c9_14830-Flag in the bound and unbound fraction, respectively. These results suggest that the binding of GBPs to E-D type TLPs evolved in bryophytes and that the number of E-D type TLPs increased in plants during evolution, likely because the binding of TLPs to GBP became crucial for plant defense against fungal pathogens.

**Figure 3.**
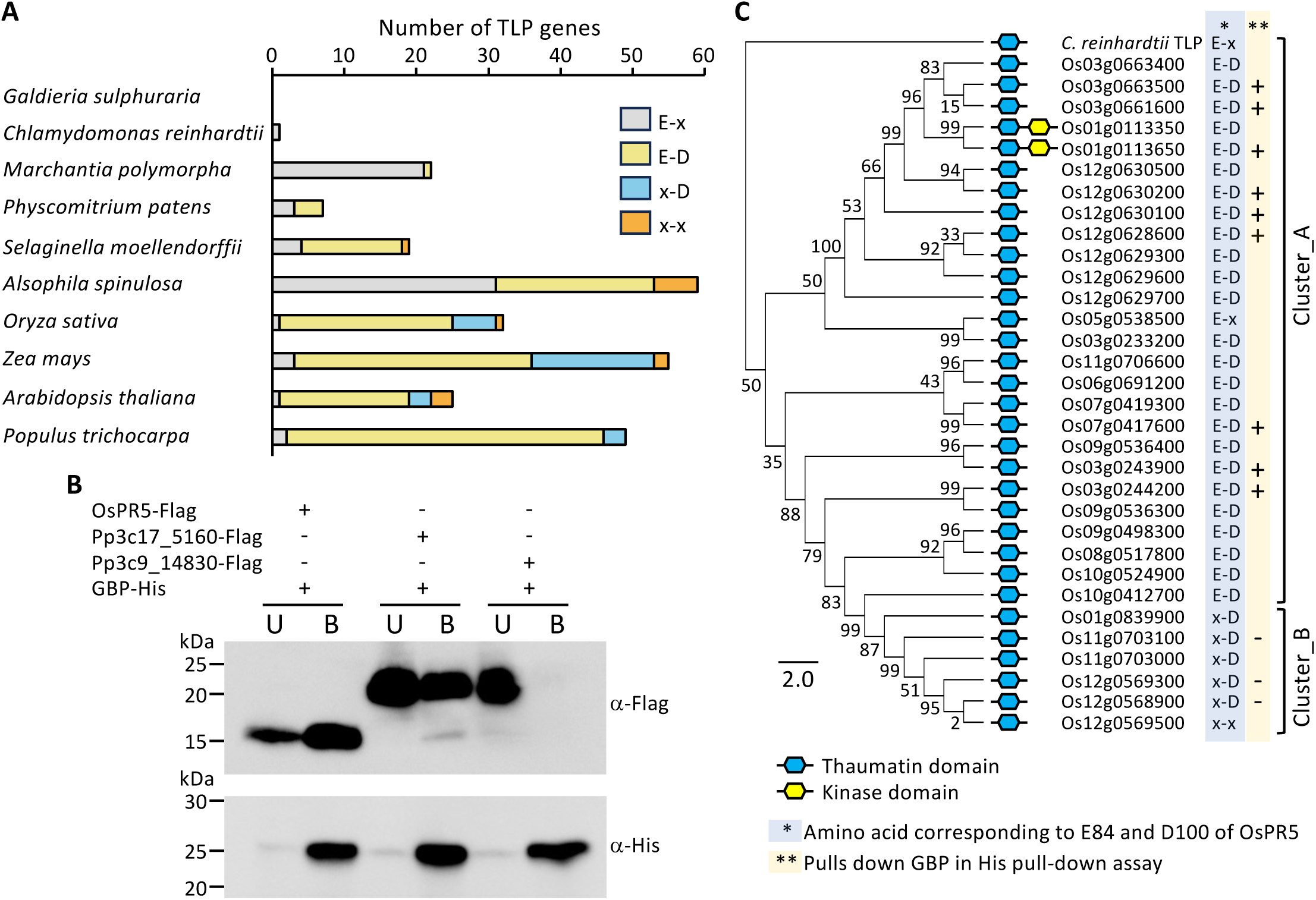
TLPs and ThKs have evolved through gene expansion and alterations in their amino acid sequences. **(A)** Number of genes encoding ThD-containing proteins in *G. sulphuraria*, *C. reinhardtii*, *M. polymorpha*, *P. patens*, *S. moellendorffii*, *A. spinulosa*, *O. sativa, Z. mays, A. thaliana* and *P. trichocarpa*. Proteins were grouped into E-x (light gray), E-D (light yellow), x-D (light blue), and x-x (light orange) types based on the conservation of amino acid residues corresponding to E84 and D100 of OsPR5. **(B)** Pull-down assay testing the binding of an E-D type (Pp05160) or E-x type (Pp14830) TLP from *P. patens* with a Flag-tag, produced in *N. benthamiana* leaves, to GBP-His using His-resin. Proteins were separated into unbound (U) and bound (B) fractions to His-resin and detected by immunoblotting using anti-Flag and anti-His antibodies. **(C)** Phylogenetic tree of rice TLPs and ThKs, reconstructed using the neighbor-joining method. The tree was reconstructed with 1,000 bootstrap replicates. Numbers indicate bootstrap values in percentages. Amino acid residues corresponding to E84 and D100 of OsPR5, and the binding to GBP as determined by pull-down assays, are indicated to the right of the tree. A TLP from *C. reinhardtii* was used as an outgroup to root the tree.

In the rice genome, we identified 30 *TLP* genes and two genes encoding a TLP fused to a kinase protein (*Thaumatin-kinase*, *ThK*) in the RAP-DB database. We reconstructed a phylogenetic tree of rice TLPs and ThKs using the amino acid sequences of the ThD region (Figure 3C and Data S2). All E-D type TLPs are grouped into cluster A in the tree. Cluster B is mainly composed of x-D type TLPs and was derived from E-D type TLPs of cluster A. This finding is in line with the tendency shown in Figure 3A that x-D type TLPs appeared later than E-D type-TLPs in higher plants. We examined the binding of 12 Flag-tagged TLPs produced in *N. benthamiana* leaves to GBP-His by pull-down assays (Figures S3 and S17). Nine OsTLPs from cluster A bound to GBP, while three OsTLPs from cluster B did not. We also examined the binding of TLPs from Arabidopsis, tomato, wheat, and maize to GBP, all of which contain amino acids equivalent to E84 and D100 of OsPR5 (Figure S18). All TLPs tested bound to GBP. These results support the hypothesis that the conserved glutamic acid (E) and aspartic acid (D) residues of TLPs corresponding to OsPR5 E84 and D100 are crucial for binding to GBP and that the function of this type of TLP is widely conserved across the plant kingdom.

The two proteins encoded by the *OsThK* genes are closely related and contain an E-D type ThD (Figure 3C). This observation suggests that the OsThKs originated from E-D type TLPs via the fusion of the TLP domain to a transmembrane domain and serine/threonine kinase domain, thereby acquiring functions in GBP binding and possibly signaling.

### Rice ThK binds to GBP and transduces defense signals

To investigate whether rice ThK has retained the GBP-binding properties of E-D type TLPs, we examined the binding of OsThK1 encoded by Os01g0113650 to GBP and ThBPs (Figure 4A). When we performed pull-down assays of OsThK1-Flag and GBP- His using His-resin, most of OsThK1-Flag was present in the bound fraction. When OsThK1-Flag and ThBP15k-His or ThBP24k-His were used for pull-downs, we detected OsThK1-Flag in the unbound fractions. We obtained identical results when using only the ThD of OsThK1 for pull-downs with GBP and ThBPs (Figure S19). These results indicate that OsThK1 interacts with GBP but not with ThBPs. We searched for *ThK* genes in the Ensembl Plants database and determined that these genes are restricted to land plants such as Poaceae, Brassicaceae, Fabaceae, and Asteraceae plants (Table S2). We tested the binding of Flag-tagged ThKs from maize (ZmThK1-Flag and ZmThK2-Flag) and Arabidopsis (AtPR5K-Flag, AtPR5K2-Flag, and AtThK3-Flag) to GBP-His by pull-down assays (Figure S20). When GBP-His was mixed with ZmThK1-Flag and AtThK3- Flag and pulled-down using His-resin, we detected both ThKs in the fractions that bound to the resin, while other ThKs accumulated in the unbound fractions. These results suggest that GBP-binding ThKs are widespread in higher plants.

**Figure 4.**
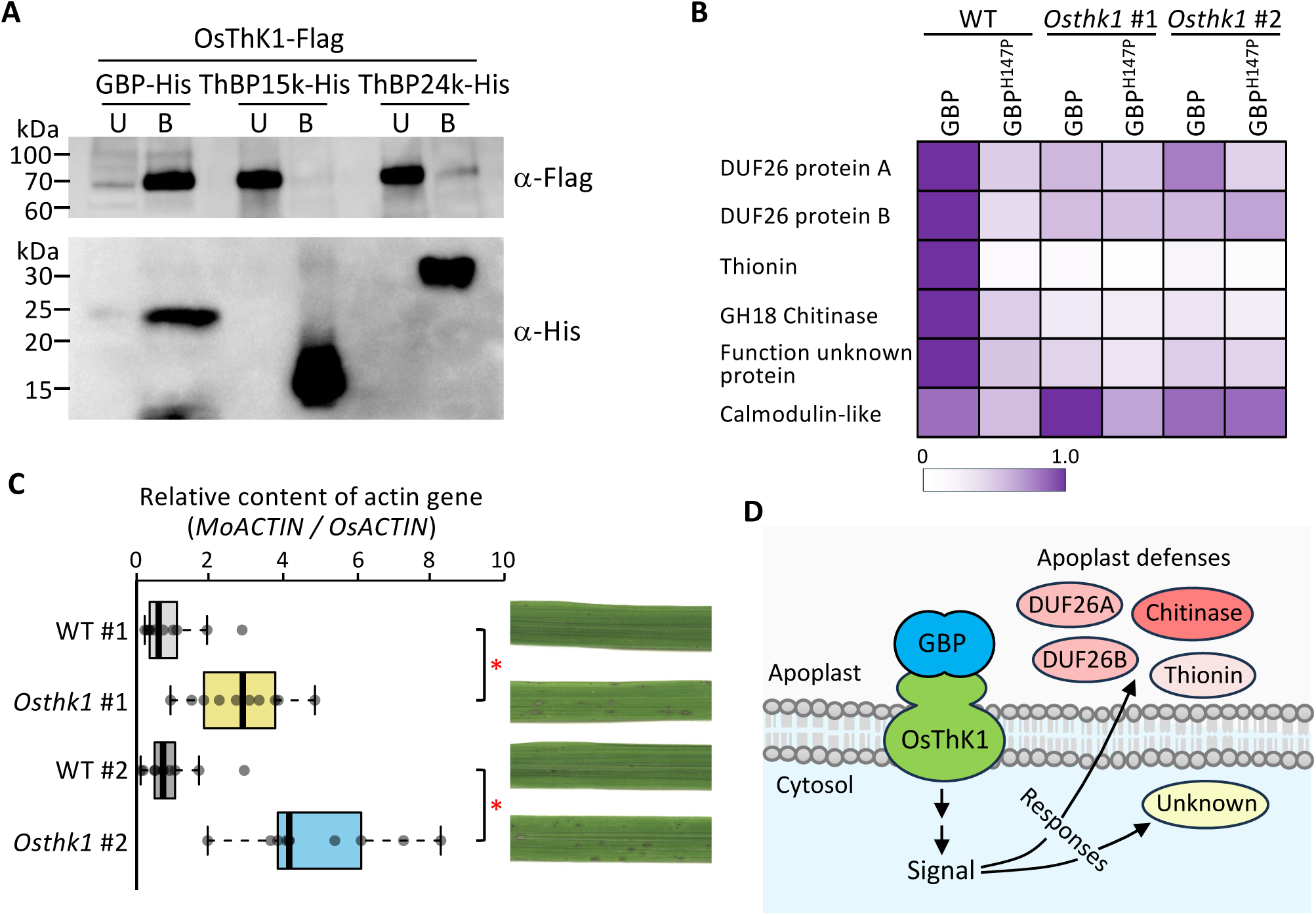
OsThK1 induces defense signaling to enhance defense responses against *M. oryzae* via GBP perception. **(A)** Pull-down assay testing the binding of a Flag-tagged rice Thaumatin-kinase 1 (OsThK1- Flag) produced in *N. benthamiana* leaves to GBP-His, ThBP15k-His, or ThBP24k-His, using His-resin. Proteins were separated into unbound (U) and bound (B) fractions to His-resin by SDS-PAGE. OsThK1-Flag and *M. oryzae* proteins were detected by immunoblotting using anti-Flag and anti-His antibodies. **(B)** Gene expression levels of the candidate genes identified downstream of ThK1. Rice callus prepared from WT and *OsThK1*-knockout mutant (*Osthk1* #1 *and* #2) plants were treated with purified GBP-His or GBP^H147P^-His (*n* = 3 for each treatment) for 6 h. Relative expression levels of DUF26 protein-A (*Os03g0277700*), DUF26 protein-B (*Os03g0312300*), Thionin (OsThionin38, *Os03g0700100*), GH18 chitinase (*Os05g0248200*), Calmodulin-like (*Os05g0491000*), and Function unknown protein (*Os01g0847100*) are shown in a heat map. *OsACTIN* was used as an internal control to normalize gene expression levels. The expression levels of untreated rice callus (time 0) were set to a ratio of 1. **(C)** *M. oryzae* infection test performed using T_1_ *Osthk1* and WT control progenies (*n* = 10 each) derived from self-pollination of T_0_ heterozygous *Osthk1* plants. The relative amount of *M. oryzae* fungal mass was determined by measuring the ratio of *M. oryzae Actin* genomic DNA (*MoACTIN*) to *O. sativa Actin* genomic DNA (*OsACTIN*) using qPCR. The average ΔΔCt value for WT was set to a ratio of 1. Data are shown as means ± SD from ten independent determinations. Asterisks indicate a significant difference at *P* < 0.01 according to a two-sided Student’s *t*-test. Representative photographs of inoculated leaves are shown to the right. **(D)** Diagram of the OsThK1 signaling pathway leading to the production of apoplastic defense-related proteins.

We further examined whether OsThK1 participates in the induction of defense-related genes using callus from WT and *OsThK1-ko* (*Osthk1*) generated by clustered regularly interspaced short palindromic repeats (CRISPR)/CRISPR-associated nuclease 9 (Cas9)- mediated gene editing (Figure S21 and Table S3). We generated two *Osthk1* mutants (*Osthk1* #1 and *Osthk1* #2) with different mutations in *OsThk1* gene (Figure S21). We incubated WT and *Osthk1* #1 callus with purified GBP-His, chitin, or BSA for 6 h and examined gene expression by transcriptome deep sequencing (RNA-seq) (Data S4). We identified six genes that were upregulated in WT but less strongly in *Osthk1* callus after GBP treatment (Figure S22). These six genes encode the following: two domain of unknown function 26 (DUF26)-containing proteins (Os03g0277700 and Os03g0312300); a thionin (OsTHI37; Os03g0700100); a glycoside hydrolase family18 (GH18) chitinase (Os05g0248200); a calmodulin-like protein (Os05g491000); and a protein of unknown function (Os01g0847100). To validate the RNA-seq data, we treated WT, *Osthk1* #1 and *Osthk1* #2 callus with purified GBP-His or GBP^H147P^-His for 6 h and analyzed expression levels by RT-qPCR (Figures 4B and S22). The expression levels of all genes except for the calmodulin-like protein gene were significantly higher in WT callus after GBP-His treatment than after GBP^H147P^-His treatment. The expression levels of the five genes were significantly lower in *Osthk1* #1 and *Osthk1* #2 callus after GBP- His treatment than in WT. These results suggest that these five genes function downstream of OsThK1.

We also investigated the role of OsThK1 in plant defense against blast fungus using WT and *Osthk1* rice plants derived from self-pollination of T_0_ heterozygous *Osthk1* #1 and *Osthk1* #2 plants. We inoculated the plants with a compatible isolate of *M. oryzae* and examined disease symptoms (Figure 4C). In parallel, we evaluated the relative fungal biomass in *Magnaporthe*-inoculated rice leaves. We observed more pronounced disease symptoms in *Osthk1* plants than in the controls, and more fungal biomass accumulated in *Osthk1* than in the WT. These results suggest that OsThK1 binds to GBP and induces the expression of at least five genes encoding proteins that function in the apoplast and cytosol, thereby enhancing plant defense (Figure 4D).

## DISCUSSION

### E-D type TLPs target GBP to inhibit its activity

*M. oryzae* GBP is secreted and binds specifically to β-1,3-glucan in the cell wall. Hydrolysis of the *gbp* cell wall in response to endo-1,3-β-glucanase treatment was strongly lower than that in the WT strain, suggesting that the *gbp* cell wall is less susceptible to β-1,3-glucan degrading enzymes (Figure S6). In addition, hyphal growth was slower in both *gbp* and WT cultures treated with recombinant purified OsPR5 (Figure 1C). Fungal growth depends on both cell division and hyphal expansion, which require a drop in cell wall strength via enzymatic degradation of cell wall polysaccharides. Our results suggest that GBP plays roles in cell wall modification and the formation of the β- 1,3-glucan network required for cell growth and that OsPR5 binds to GBP as its target molecule to inhibit its action.

Plant TLPs are found in a wide range of species of the green lineage, from algae to angiosperms, and are involved in defense against pathogens, as evidenced by the enhanced and compromised resistance against pathogens observed in plants overexpressing or knocked down for *TLP* genes, respectively.^29,30^ Nevertheless, the mode of action of TLPs had been unclear until now. In this study, we identified *M. oryzae* GBP, ThBP24k, and ThBP15k as interacting proteins of OsPR5 by pull-down assays. Understanding how these proteins interact with each other sheds light on the apoplastic molecular battle that influences pathogen infection and plant defense.

According to the structural model of the OsPR5–GBP complex predicted by AlphaFold3, GBP is docked in the cleft region of OsPR5. Furthermore, we propose that interactions of E84 and D100 in OsPR5 with H147 and S148 in GBP are crucial for OsPR5–GBP complex formation based on an analysis of inter-atomic distances in the complex and a binding test to a series of OsPR5 and GBP variants with single amino-acid substitutions (Figures 1F and 1H). The amino acid residues responsible for protein– protein interactions are highly conserved among plant TLPs named as E-D type TLPs.

Among the 113 GBP homologous proteins, a total of 14 amino acid residues including H147 and S148 are highly conserved (>95%), most of which are likely required for maintaining β-1,3-glucan binding surface (Figure S23); however, the canonical two amino acid residues H147 and S148 are targeted by E-D type TLPs for binding (Figure 5).

**Figure 5.**
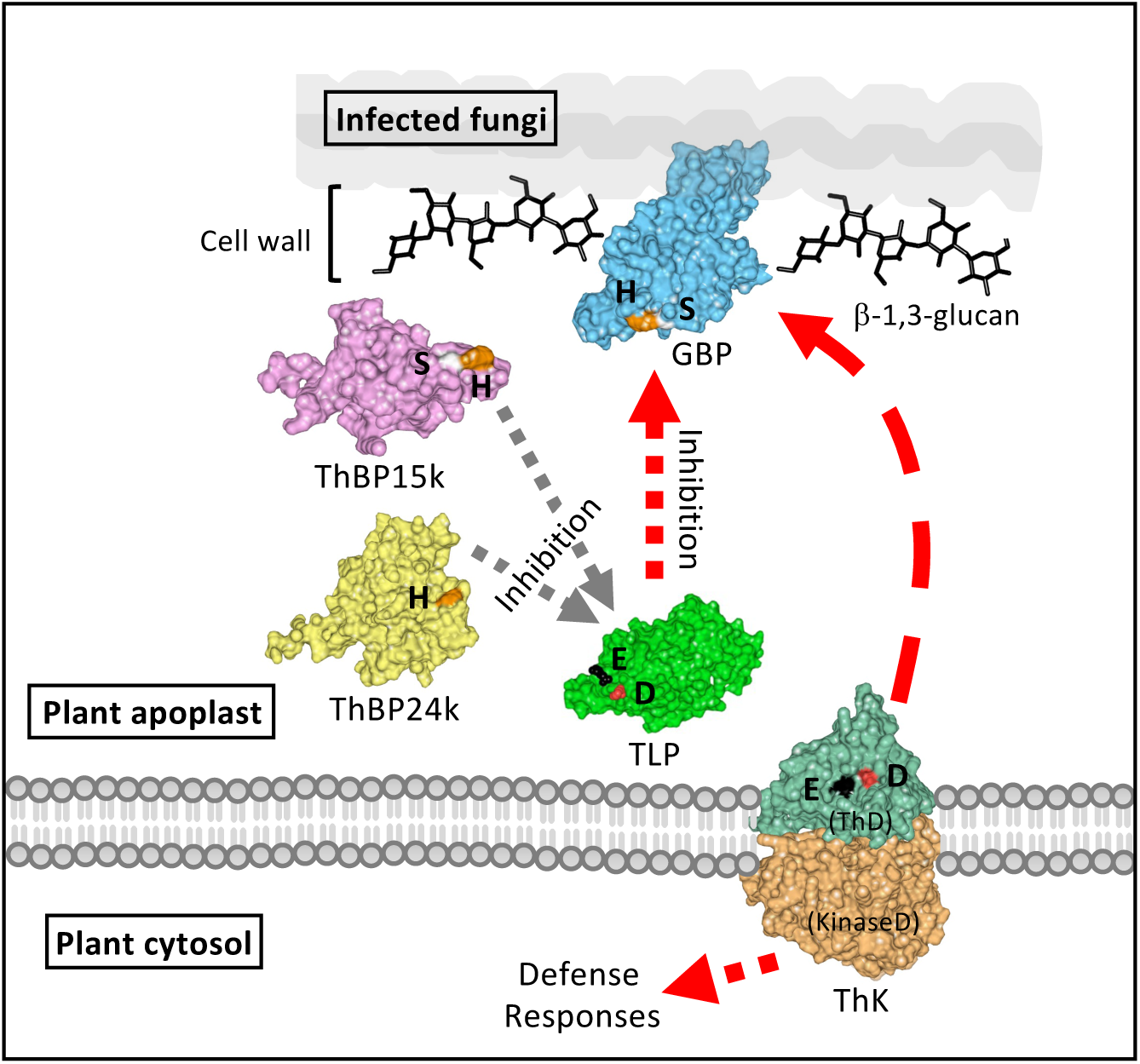
Proposed model of the interactions between Plant E-D type TLPs and fungal GBP during the infection of rice by blast fungus. Plant E-D type TLPs inhibit the action of fungal GBP. As a countermeasure, fungal ThBPs bind to E-D type TLPs to interfere with the binding of TLPs to GBP. Plant ThKs containing an E-D type thaumatin domain perceive GBP to trigger defense signaling, thereby enhancing immunity.

Several TLPs show endo-1,3-β-glucanase activity and/or antifungal activity, but these properties are not always correlated.^9^ In the current study, the β-1,3-glucanase activities of 11 rice TLPs were far lower than that of commercial β-1,3-glucanase (1.5 mU) (Figure S24). A catalytic Glu-Asp (E-D) pair in the acidic cleft region of TLPs is predicted to be required for β-1,3-glucanase activity.^31^ However, neither E-D type TLPs nor x-D type TLPs in rice possess much if any hydrolytic activity. Therefore, the conservation of these amino acid residues among rice TLPs is not sufficient to support their endo-1,3-β- glucanase activity. Since E-D type TLPs tightly bind to GBP in the native fungal cell wall (Figure S10), they may not be able to access β-1,3-glucan for binding and hydrolysis. Our findings suggest that the major function of E-D-type TLPs is to bind to and sequester GBP from the fungal cell wall to confer immunity.

### ThBPs abolish the binding of E-D type TLPs to GBP as a countermeasure

Plant pathogens appear to have developed strategies for evading the threat of TLPs. *A. alternata* Alta1 and *V. dahlia* PevD1 are extracellular effectors that inhibit the hydrolytic activity and antifungal activity of PR5s.^24,25^ These proteins share low sequence similarity (27%) but are structurally similar. Protein–protein docking calculations based on the experimental X-ray structure of PevD1 and the predicted structure of GhPR5 suggested that a region corresponding to aa 113–155 of PevD1 interacts with the electrostatically negative cleft of GhPR5. *M. oryzae* ThBP24k and ThBP15k identified in this study are also two extracellular effectors that are secreted into the apoplastic space during the infection of rice plants and bind to TLPs. Interactions of H44 of ThBP24k with E84 of OsPR5, and H22 and S23 of ThBP15k with E84 and D100 of OsPR5, are important for protein–protein complex formation (Figures 2D, 2E, 2H and 2I). ThBPs might serve as inhibitor proteins of E-D type TLPs. Indeed, ThBP24k and ThBP15k abolished the interaction of OsPR5 with GBP (Figure 2A). E-D type TLPs must maintain the D and E residues in suitable positions for GBP binding, explaining how E-D type TLPs are unable to avert the attack of ThBPs (Figure 5). Neither Alta1, PevD1, nor a *M. oryzae* PevD1-homolog (MGG_02678) possesses the corresponding amino acid residues of ThBPs, indicating that these proteins function in a different manner from ThBPs. Plant pathogens may have therefore evolved a variety of TLP-binding proteins to overcome the threat of TLP actions.

### The functions of TLPs in plant–pathogen interactions have diversified

Based on the Ensembl Plants database, an E-x type TLP is present in Chlamydomonas, suggesting a possible origin for TLPs in the green lineage. In the bryophytes *M. polymorpha* and *P. patens*, the number of E-x type TLPs increased, and subsequently, the E-D type TLPs appeared. The E-D type TLP Pp3c17_5160 binds to GBP, suggesting that the binding of GBP to TLPs first appeared in bryophytes. Thereafter, the number of E-D type TLPs markedly increased in ferns and higher plants (Figure 3A). These observations suggest that E-D type TLPs play significant roles in defense responses against plant pathogens. In higher plants, x-D type TLPs incapable of binding to GBP have emerged. The detailed functions of this class of TLPs should be analyzed in future studies.

Another important event in the evolutionary history of TLPs is the emergence of the receptor-like kinases (RLKs) ThKs. ThKs have a modular structure, with the N-terminal extracellular ThD connected to a cytoplasmic serine/threonine kinase-domain by a transmembrane-spanning domain. Whereas TLPs are widespread in plants, ThKs are restricted to land plants (Table S2). Thus far, the few examples of characterized Thks play roles in abiotic stress responses. The Arabidopsis ThK PR5-LIKE RECEPTOR KINASE (AtPR5K) shows auto-phosphorylation activity, indicative of serine/threonine kinase activity.^32^ Its homolog AtPR5K2 interacts with the protein phosphatases ABA-INSENSITIVE 1 (ABI1) and ABI2, and overexpressing *AtPR5K2* enhanced drought tolerance in transgenic Arabidopsis plants.^33^ However, the involvement of ThKs in defense responses against plant pathogens has not been addressed. In the current study, combined analyses of gene expression by RNA-seq and RT-qPCR identified five candidate genes that may function downstream of OsThK1 signaling (Figures 4B and S22). The perception of pathogen-derived GBP by OsThK1 significantly contributes to plant defense in both the apoplast and cytosol (Figures 4D and Figure 5). How OsThK1 is activated upon binding to GBP should be addressed in the future. Notably, OsThK1 is not bound by ThBPs, likely due to steric hindrance (Figure 4A), thereby avoiding interference from these pathogen proteins.

Based on a phylogenetic tree of rice OsTLPs and OsThKs, OsThKs may have originated from E-D type TLPs, suggesting that OsThKs were generated via the fusion of a ThD from an E-D type TLP to a transmembrane domain and serine/threonine kinase domain (Figure 3C). It would be interesting to explore how frequently such fusions of plant apoplastic defense proteins with signaling modules have occurred during evolution.^34^ We previously observed a similar fusion of a plant domain to a nucleotide-binding leucine rich repeat (NLR)-type cytoplasmic receptor, where pathogen effector target domains were integrated with NLR sensors.^35^ Such non-homologous recombination-mediated domain shuffling between proteins involved in host–pathogen interactions and signaling modules may thus be prevalent in plants, providing the opportunity for natural selection to act in the evolution of efficient defense responses against rapidly evolving pathogens.

### Possible application of our findings for controlling pathogen spread

In this study, we revealed a previously unknown molecular mechanism for the interactions of the plant proteins OsTLPs and OsThK1 with the *M. oryzae* proteins GBP, ThBP24k, and ThBP15k, which was established during the coevolution between plants and plant pathogens (Figure 5). E-D type TLPs target fungal GBP to inhibit its function, but ThBPs interfere with the action of TLPs. OsThK1 binds to GBP via the ThD and transduces signals to induce the accumulation of defense-related proteins. Our findings could be used to generate pathogen-resistant plants with enhanced accumulation of TLPs and ThKs to help improve crop yields. Specifically, similar to the study of cross-family transfer of EFR receptor gene,^36^ the utilization of ThKs by gene transfer and crossbreeding could enhance pathogen resistance in crops because many plant species do not possess *ThK* genes.

## MATERIALS AND METHODS

### Microorganisms and growth conditions

After the emergence of 7th leaf, the 5th and 6th leaves were homogenized in 100 mM sodium phosphate buffer (pH7.0). The resulting pellet obtained by centrifugation at 20,000 x *g* for 10 min at 4°C were used as the rice cell wall fraction for supplementing liquid culture. The fungus *Magnaporthe oryzae* (pathovar Ken53-33) was cultured on oatmeal agar plates or yeast glucose (YG) medium (0.5% [w/v] yeast extract, 2% [w/v] glucose) with or without 0.1% [w/v] rice cell wall fraction at 25°C. Conidia formation was induced by incubation under dark-blue light at 28°C for 4 days. Agrobacterium (*Agrobacterium tumefaciens*) strain GV3101 carrying specific plasmids was grown in YEB medium (0.5% [w/v] yeast extract, 1% [w/v] peptone, 0.5% [w/v] beef extract, 0.5% [w/v] sucrose) supplemented with rifampicin (50 μg/mL) and kanamycin (100 μg/mL) at 28°C.

### Plant materials and growth conditions

The rice (*Oryza sativa*) cultivar ‘Moukoto’ and *Nicotiana benthamiana* plants were grown in soil at 28°C or 25°C, respectively. Rice suspension cells were cultured at 28°C with rotation at 140 rpm and transferred to fresh medium every 10 days as described previously.^37^

### Recombinant protein preparation

Plant TLPs were produced in rice suspension cells and *N. benthamiana* leaves as described previously.^34,38^ His-tagged proteins were purified using His-tag affinity resin (His-resin, Takara Bio). GBP-His and its variants were produced in *Escherichia coli* (*Origami*) cells harboring the pCold2 vector (Takara Bio) in which the respective coding sequence without secretion signal peptide coding sequence had been cloned according to manufacturer’s instructions. Recombinant ThBP24k, ThBP15k, and their variants were produced in *M. oryzae* as described previously.^39^

### Pull-down assays to identify OsPR5-binding proteins

Recombinant OsPR5-His purified from rice suspension cells was incubated with a crude protein preparation from a *M. oryzae* culture filtrate in binding buffer (50 mM sodium phosphate buffer, pH 7.5, containing 150 mM NaCl) for 15 min at 4°C as described previously.^34^ The mixture was then incubated with His-resin for 30 min at 4°C. After washing the His-resin with binding buffer containing 5 mM imidazole, proteins bound to the resin were eluted using binding buffer containing 200 mM imidazole. Eluted proteins were analyzed by SDS-PAGE and visualized using the SilverQuest Silver Staining Kit (Thermo Fisher Scientific).

### Peptide identification

Protein bands separated by SDS-PAGE were excised and subjected to in-gel trypsin digestion as described previously.^40^ Peptide identification was performed as described previously.^41^ Digested peptides were loaded onto a Magic C18 AQ nano column (0.1 x 150 mm, MICHROM Bioresources, Inc.) using an ADVANCE UHPLC system (MICHROM Bioresources, Inc.) equilibrated with 0.1% (v/v) formic acid in acetonitrile and eluted using a linear gradient of 5–45% (v/v) acetonitrile at a flow rate of 500 nL/min. Mass spectrum analysis was performed using an LTQ Orbitrap XL mass spectrometer (Thermo Fisher Scientific) with Xcalibur software ver. 2.0.7 (Thermo Fisher Scientific). Peptides were identified using a MASCOT MS/MS ion search (http://www.matrixscience.com/home.html) in error tolerance mode (one amino acid substitution allowed) with the NCBI database. Search parameters were as follows: taxonomy, plants; max missed cleavages, 0; fixed modifications, carbamidomethyl; peptide tolerance, ± 5 ppm; fragment mass tolerance, ± 0.6 Da.

### Binding assay with His-tagged proteins

For pull-down assays, crude protein preparations containing His-tagged or Flag-tagged proteins were incubated for 15 min at 4°C. The mixture was then incubated with His-resin for 15 min at 4°C. Following centrifugation at 2,400 x *g* for 1 min at 4°C, the supernatant was collected as the unbound fraction. Proteins bound to the His-resin were eluted with binding buffer containing 200 mM imidazole and collected as the bound fraction. Both fractions wereanalyzed by SDS-PAGE, followed by immunoblotting using anti-His (Qiagen, ID. 34460) and anti-Flag (MBL Life Science, Code No. PM020-7) antibodies. For binding to β-1,3-glucan, His-tagged TLPs were incubated with water- insoluble β-1,3-glucan in 100 mM sodium phosphate buffer (pH 6.0) containing 150 mM NaCl for 30 min at 4°C. The supernatant, obtained by centrifugation at 2,400 x *g* for 1 min at 4°C, was collected as the unbound fraction. The pellet was washed with the same buffer, and the bound proteins were eluted by boiling in SDS-PAGE sample buffer. The protein fractions unbound and bound to β-1,3-glucan were subjected to immunoblotting using anti-His antibodies.

### Protein detection

Proteins were detected by silver staining and immunoblotting using antibodies against peptide epitope tags, as well as specific antibodies against OsPR5, GBP, ThBP24k and ThBP15k. The anti-OsPR5, anti-GBP, anti-ThBP24k and anti-ThBP15k antibodies were generated in mice using the following synthetic peptides (*N*-SGQKPLTLAEFTIGGSQC- *C* for anti-OsPR5, *N*-SPSVHGKAIANSGRC-*C* for anti-GBP, *N*- CQRPDEFDLPANDKS-*C* for anti-ThBP24k, and *N*-CINSRTNQDYCHKIP-*C* for anti- ThBP15k). Detection was performed using HRP-conjugated secondary anti-mouse IgG antibody produced in goat (MBL Life Science, Code No. 330). Protein concentration was determined using the Bradford Protein Assay Kit (Thermo Fischer Scientific), with bovine serum albumin (Sigma-Aldrich) as the standard.

### Binding kinetics assay

Binding kinetics of OsPR5 to GBP, ThBP24k, or ThBP15k was evaluated by measuring integrated signal intensity using a BLItz instrument and BLItz Pro software (ForteBio) as described previously.^42^ Recombinant purified OsPR5-His (10 μL, 2.0 μM) was immobilized onto a His-tag biosensor and incubated for 10 min at 25°C. A baseline of binding response was obtained by incubating the sensor with binding buffer (50 mM sodium phosphate buffer, pH 7.5, 150 mM NaCl) for 10 min. The association phase was initiated by immersing the sensor in a solution containing GBP or ThBPs (10 μL, 2.0 μM) for 2 min. Protein dissociation was measured for 2 min by incubating the sensor with binding buffer. Time zero was defined as the starting point of protein association. The fold increase in integrated intensity was calculated by dividing each trajectory by the value at time zero.

### Generation of variant proteins

DNA substitutions were introduced by PCR using DNA primers with the necessary mutations as listed in Table S4 The resulting variant proteins were produced as described above.

### Cell wall hydrolysis by endo-β-1,3-glucanase

*M. oryzae* wild-type (WT) and *gbp* mutant (see below) strains were cultured in YG medium for 3 days. Hyphae were harvested and sonicated in 100 mM sodium phosphate buffer (pH 7.0). The resulting pellets obtained by centrifugation at 10,000 x *g* for 5 min at 4°C were washed with the same buffer and used as a cell wall preparation. The cell wall preparations were treated with *Aspergillus oryzae* endo-1,3-β-glucanase (2.0 units/mL, Megazyme) for 18 h at 30°C. The amount of released sugar in the supernatant obtained by centrifugation at 10,000 x *g* for 5 min at 4°C was measured by the PAHBAH (*p*-hydroxybenzoic acid hydrazide) method.^43^

### Measurement of *M. oryzae* hyphal length

Conidia (0.5 × 10^5^) from *M. oryzae* WT, the *gbp* mutant, or the *gbp:GBP* complementation strain were resuspended in 20 mM sodium phosphate buffer (pH 5.5) and placed on a glass slide. BSA, OsPR5, or OsPR5^E84K^ (1.0 μg each) was added to the conidial suspension. After 7 h of incubation, hyphae from germinated conidia were photographed, and their lengths were measured using ImageJ software.

### β-1,3-Glucanase activity assay

His-tagged TLPs were produced in *N. benthamiana* leaves and purified using His-resin. Each prepared protein (0.2 μg) was incubated with 0.1% (w/v) laminarin (Sigma-Aldrich) in 100 mM sodium phosphate buffer (pH 6.0) at 30°C for 18 h. Hydrolyzed products were quantified using the PAHBAH method as above.

### Generation of transgenic *M. oryzae* strains and rice plants

Knockout mutants of *GBP*, *ThBP24k*, and *ThBP15k* were generated by the TALEN method as described previously.^44^ *M. oryzae* mutants were selected on potato-dextrose agar plates containing 250 μg/mL bialaphos (FUJIFILM WaKo, 022-15413). A 298-bp *OsPR5* DNA fragment corresponding to nucleotides 411–708 of open reading frame was cloned into the pANDA vector for RNA interference as described previously.^45^ Knockout rice mutants for *OsThK1* were generated using single guide RNAs (sgRNAs) recognizing the target DNA sequences 5’-tcagacgctcgctgagttcacgg-3’ and 5’-ccggccaacatcacgtcgcagtg- 3’ by the CRISPR/Cas9 method as described previously.^46^ The rice cultivar Moukoto was used for Agrobacterium-mediated transformation as described previously.^37^ Transgenic rice plants were selected on medium containing 300 μg/mL hygromycin B; genomic DNA was extracted from the plants by NucleoSpin Plant II (Macherey-Nagel) and used as a template for PCR amplification and Sanger sequencing of the *OsThK1* locus to verify the presence of mutations (Figure S21). Genes sharing high similarity (>70%) to the target DNA sequences were identified by BLAST search using the RAP-DB database (Table S3). Six candidate genes with potential off-target sites were amplified by PCR and their DNA sequences were verified by Sanger sequencing to confirm the absence of off-targets mutations.

### *M. oryzae* inoculation of rice plants

Leaves of rice plants (cultivar Moukoto) at the 6-leaf stage were spray-inoculated with *M. oryzae* spores (1.0–3.0 × 10^5^; compatible strain Ken53-33) in 20 mM sodium phosphate buffer with 0.01% (v/v) Tween-20.

### Quantitative PCR and reverse-transcription quantitative PCR

Genomic DNA and total RNA were extracted from rice leaves (cultivar Moukoto), rice suspension culture cells, and *M. oryzae* hyphae. Quantitative PCR (qPCR) and RT-qPCR (RT-qPCR) were conducted using SYBR GreenER qPCR Super Mix (Invitrogen) and specific primers (Table S4) in a StepOnePlus Real-Time PCR system (Applied Biosystems). For qPCR, the relative *M. oryzae* DNA content was calculated using the ΔΔCT method, with *M. oryzae Actin* genomic DNA (*MoACTIN*, MGG_03982) as the target gene and rice *Actin* genomic DNA (*OsACTIN*, Os03g0718100) as the reference, as described previously (Takeda et al., 2022). The average ΔΔCt value for WT was set to a ratio of 1. Data are presented as means ± standard deviation (SD) from independent determinations. For RT-qPCR, expression levels were normalized to *MoACTIN.* Data are presented as means ± standard deviation (SD) from independent determinations.

### RNA-seq analysis of rice callus

Rice callus was treated with 2 μg/mL BSA, 4 μg/mL chitin, or 2 μg/mL GBP at 28°C for 6 h. Total RNA was extracted from treated callus using the SV Total RNA Isolation System (Promega). One microgram of total RNA was used to prepare sequencing libraries with an NEB Next Ultra II Directional RNA Library Prep Kit for Illumina (NEB). The libraries were sequenced by paired-end sequencing (2 × 150 PE) using a Illumina Nova Seq platform (Illumina). The quality of the RNA-seq data was evaluated using FastQC (http://www.bioinformatics.babraham.ac.uk/projects/fastqc/). Hisat2 was used to align the RNA-seq reads against the *O. sativa* (cultivar Nipponbare) reference genome downloaded from RAP-DB (https://rapdb.dna.affrc.go.jp/).^47^ Expression levels were measured as Transcripts Per Million (TPM) using TPMCalculator.^48^

### DNA sequencing

DNA was amplified from cDNA by PCR using PrimeStar GXL DNA polymerase (Takara Bio) and verified by Sanger sequencing on a 3130 Genetic Analyzer (Applied Biosystems). The primers used in this study are listed in Table S4.

### Structural modeling of protein complexes

Modeling of protein complexes of TLPs with GBP, ThBP24k, or ThBP15k was carried out using the AlphaFold3 server (https://alphafoldserver.com, accessed in December, 2024). The display of steric structures and measurement of interatomic distances were performed using Waals software (Altif Laboratories Inc.) and UCSF ChimeraX (https://www.cgl.ucsf.edu/chimerax/). Amino acid interactions of protein-protein complexes were visualized using LigPlot+ v.2.2 (https://www.ebi.ac.uk/thornton-srv/software/LigPlus/). Molecular docking of GBP and β-1,3-glucan hexaose was performed using the Protenix server (protenix-server.com).

### Search for homologous proteins of TLP, GBP and ThBPs

Amino acid sequences containing the thaumatin domain (ThD, Pfam PF00314) in *G. sulphuraria*, *C. reinhardtii*, *M. polymorpha*, *P. patens*, *S. moellendorffii*, *O. sativa, Z. mays*, *A. thaliana*, and *P. trichocarpa* were retrieved using keywords ‘thaumatin’ and ‘Pfam: PF00314’ from the Ensembl Plants (https://plants.ensembl.org/index.html) database. Amino acid sequences from *A. spinulosa* were obtained by BLAST search (e < 1e-4) using the OsPR5 amino acid sequence as the query in the Figshare database (https://figshare.com). Homologous protein sequences of GBP and ThBPs were retrieved by BLAST search (E < 1e-5) using GBP and ThBPs as queries from the NCBI database (https://www.ncbi.nlm.nih.gov).

### Phylogenetic tree reconstruction

Amino acid sequences were aligned using CLUSTALW (https://www.genome.jp/tools-bin/clustalw), and a maximum-likelihood phylogenetic tree was reconstructed with 1,000 bootstrap replicates. The amino acid sequences used for tree reconstruction are listed in Data S2.

### WebLogo construction

An amino acid sequence logo was constructed using homologous sequences of GBP, ThBP24k, and ThBP15k from 113, 53, and 48 fungal species, respectively, using WebLogo (https://weblogo.berkeley.edu/logo.cgi).

## ACKNOWLEDGMENTS

We thank Sophien Kamoun for valuable comments to improve the manuscript.

