## Supplemental file for "Multipartite coevolution shapes plant apoplastic immunity against rice blast fungus"

Supplementary data

Supplementary Figures (Figure S1-24)

Supplementary Tables (Table S1-4)

Supplementary Figure S1

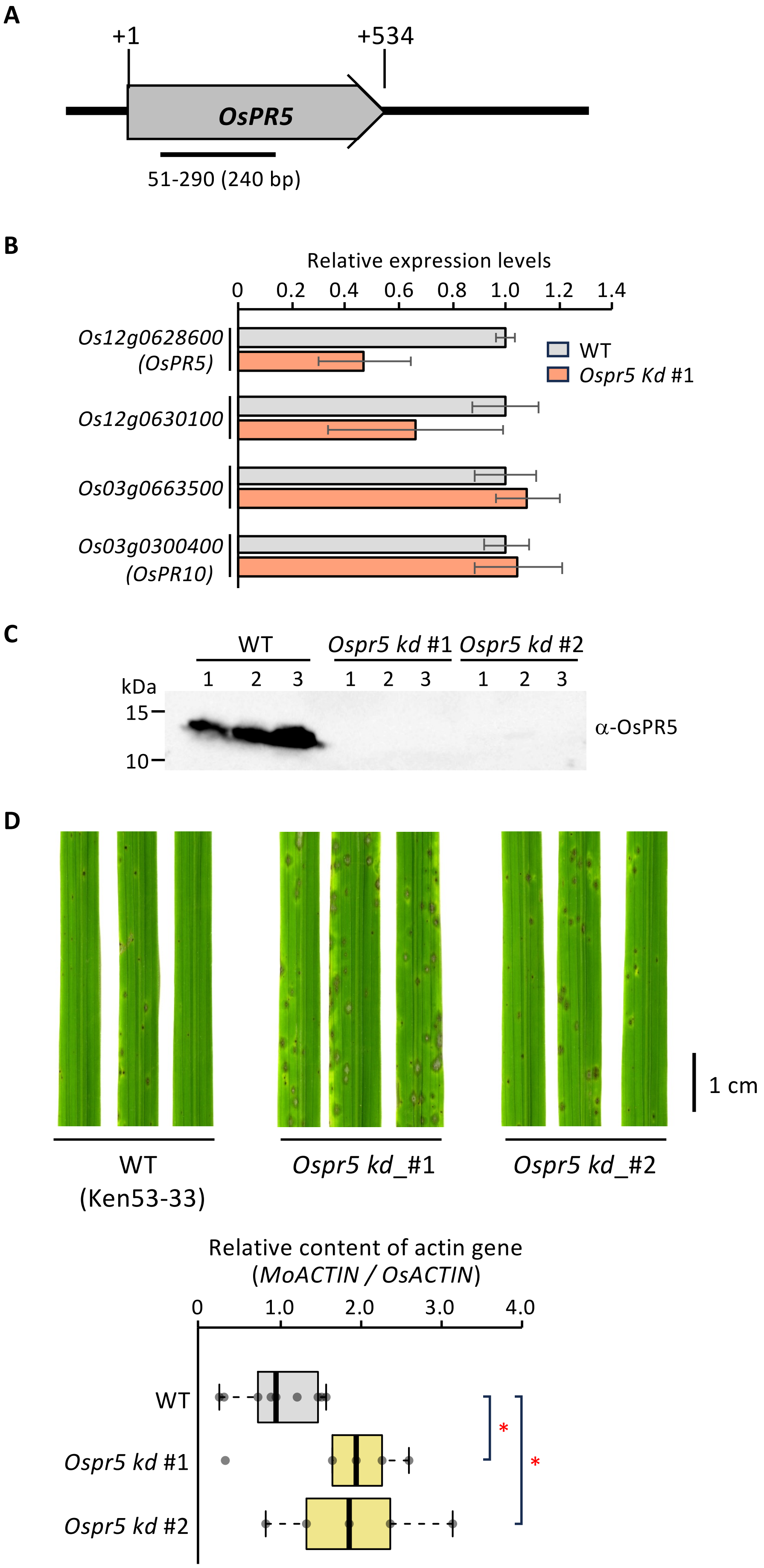

**Figure S1. *OsPR5* knockdown rice lines show diminished resistance to *M. oryzae infection***

**(A)** *OsPR5* knockdown mutants (*Ospr5 kd* #1 and #2) in the rice cultivar Moukoto were generated using RNA interference (RNAi)-mediated gene silencing with a pANDA vector containing a 240-bp segment of the *OsPR5* open reading frame. **(B)** Rice wild type (WT) and *Ospr5 kd* #1were inoculated with *M. oryzae* (Ken 53-33), and the leaves were used for gene expression analysis of Os12g0628600 (*OsPR5*) and *OsPR5*-like *TLP*s Os12g0630100, Os03g0663500, and Os03g0300400 (*OsPR10*, as a control), as analyzed by RT-qPCR. Rice Actin gene was used as an internal control to normalize gene expression levels. The average $\Delta\Delta$Ct value for WT was set to a ratio of 1. Data are shown as means $\pm$ standard deviation (SD) from three independent determinations. **(C)** Immunoblot analysis of OsPR5 accumulation in the leaves of WT and *Ospr5 kd* (#1 and #2) 5 days after *M. oryzae* inoculation, using an anti-OsPR5 antibody. **(D)** Representative photographs of rice leaves from WT ($n$ = 9) and *Ospr5* (#1 and #2, $n$ = 5 each) plants spray-inoculated with *M. oryzae*. Disease symptoms and *M. oryzae* fungal mass were assessed in rice leaves at 5 days post-inoculation. The relative amount of *M. oryzae* fungal mass in the leaves was quantified by measuring the ratio of *M. oryzae Actin* genomic DNA (*MoACTIN*) to *O. sativa Actin* genomic DNA (*OsACTIN*) using qPCR. The average $\Delta\Delta$Ct value for WT was set to a ratio of 1. Data are shown as means $\pm$ SD from independent determinations. Asterisks indicate significant differences at $P < 0.01$, as determined by a two-sided Student's t-test.

### Supplementary Figure S2

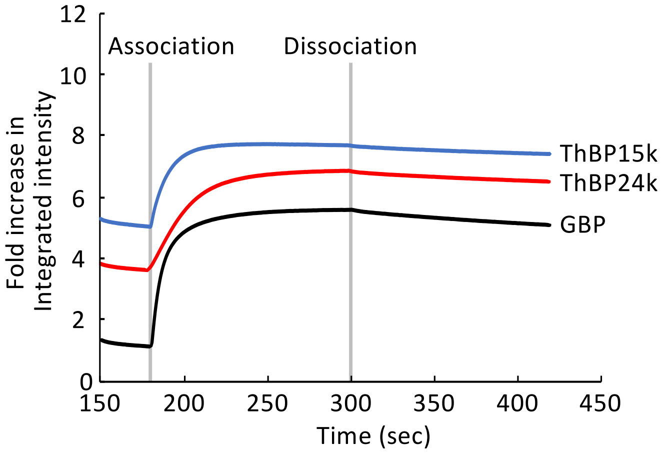


Binding kinetics of OsPR5 and *M. oryzae* proteins

| | $K_a$ | $K_d$ | $K_D$ |
|---|---|---|---|
| OsPR5 / GBP | $2.02E^{6} \pm 5.74E^{3}$ | $1.51E^{-3} \pm 2.01E^{-5}$ | $7.45E^{-10} \pm 9.06E^{-11}$ |
| OsPR5 / ThBP24k | $4.58E^{5} \pm 1.91E^{3}$ | $1.11E^{-3} \pm 9.18E^{-6}$ | $2.41E^{-9} \pm 2.42E^{-10}$ |
| OsPR5 / ThBP15k | $5.63E^{4} \pm 3.31E^{2}$ | $1.46E^{-3} \pm 1.56E\text{-}^{5}$ | $2.59E^{-8} \pm 3.68E^{-9}$ |

**Figure S2. Integrated intensity representing the binding kinetics of OsPR5 to *M. oryzae* proteins**

The association of purified GBP-His, ThBP24k-His, and ThBP15-His with OsPR5-His was measured from 180 sec to 300 sec. Binding buffer was used to measure protein dissociation from 300 sec to 420 sec. Fold increase in integrated intensity was calculated by dividing each trajectory by the value at 180 sec. The figure shows representative traces from three independent experiments. Values of the binding kinetics are summarized in Table.

Supplementary Figure S3

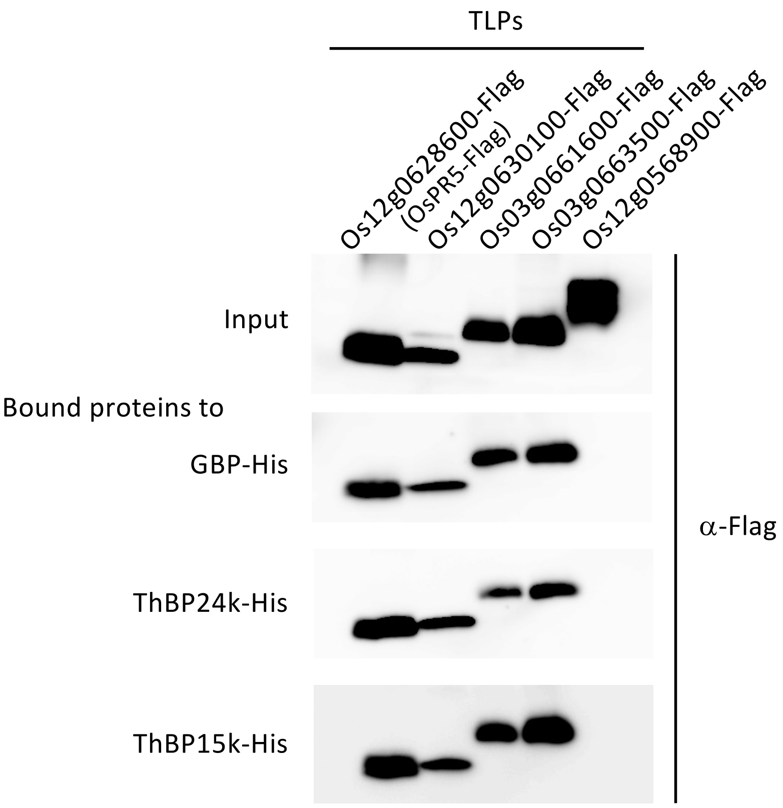


**Figure S3. Various TLPs bind to GBP, ThBP24k, and ThBP15k**

OsTLPs encoded by Os12g0628600, Os12g0630100, Os03g0661600, Os03g0663500, and Os12g0568900, which all accumulate to high levels in rice leaves infected with *M. oryzae* (Table S1), were produced with a Flag-tag in *N. benthamiana* leaves. TLP-Flag proteins were examined for binding to recombinant GBP-His, ThBP24k-His, and ThBP15k-His by pull-down assays using His-resin. The fractions bound to His-resin were analyzed by immunoblotting using an anti-Flag antibody.

Supplementary Figure S4

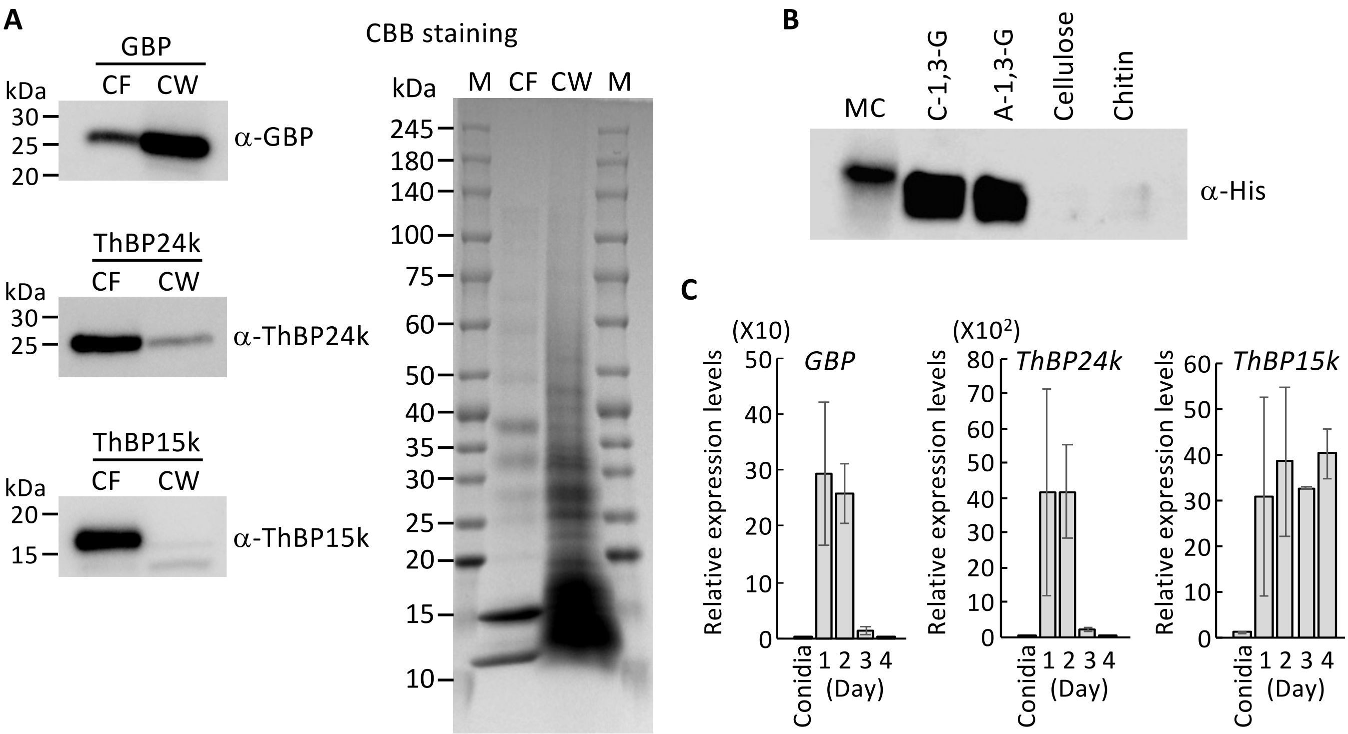


**Figure S4. The localization, binding properties and gene expression of GBP, ThBP24k, and ThBP15k define their extracellular interactions with TLPs**

**(A)** *M. oryzae* was cultured for 3 days in YG medium containing rice cell wall material. The culture was separated into culture filtrate (CF) and cell wall (CW) fractions. Proteins equivalent to 0.5 mL of *M. oryzae* culture were loaded to SDS-PAGE. GBP, ThBP24k, and ThBP15k accumulation was determined by immunoblotting using anti-GBP, anti-ThBP24k, and anti-ThBP15k antibodies. Proteins of CF and CW were stained with Coomassie brilliant blue R-250 (CBB) to show total protein loaded for gel electrophoresis. M indicates protein molecular marker. **(B)** GBP-His overexpressed in *M. oryzae* was used in binding experiments with an *M. oryzae* cell wall fraction (MC), crystalline β-1,3-glucan (C-1,3-G), alkaline-swollen β-1,3-glucan (A-1,3-G), cellulose, and chitin. Bound proteins were analyzed by immunoblotting using an anti-His antibody. **(C)** Relative gene expression levels of GBP, ThBP24k, and ThBP15k during fungal infection to rice. The gene expression levels of GBP, ThBP24k, and ThBP15k were analyzed in conidia and rice leaves after 1-4 days of *M. oryzae* inoculation ($n$ = 3 each) by RT-pPCR. *MoACTIN* was used as an internal control to normalize gene expression levels. Data are shown as means ± SD from three independent determinations.

### Supplementary Figure S5

**A**

Immunoblotting

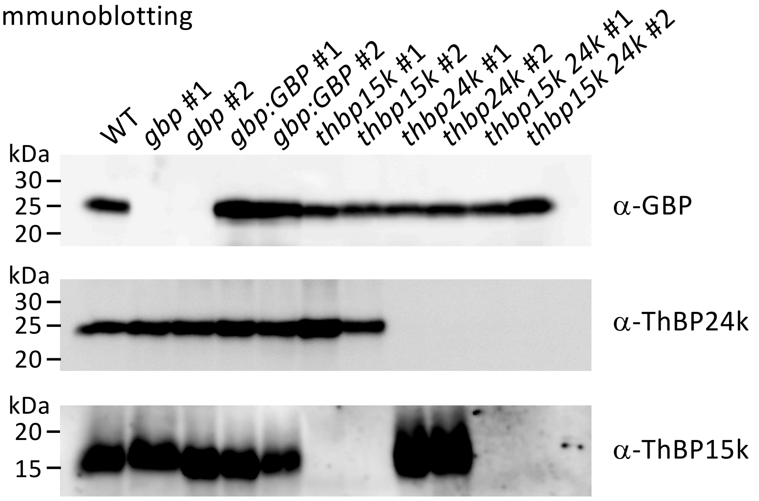


**B**

CBB staining

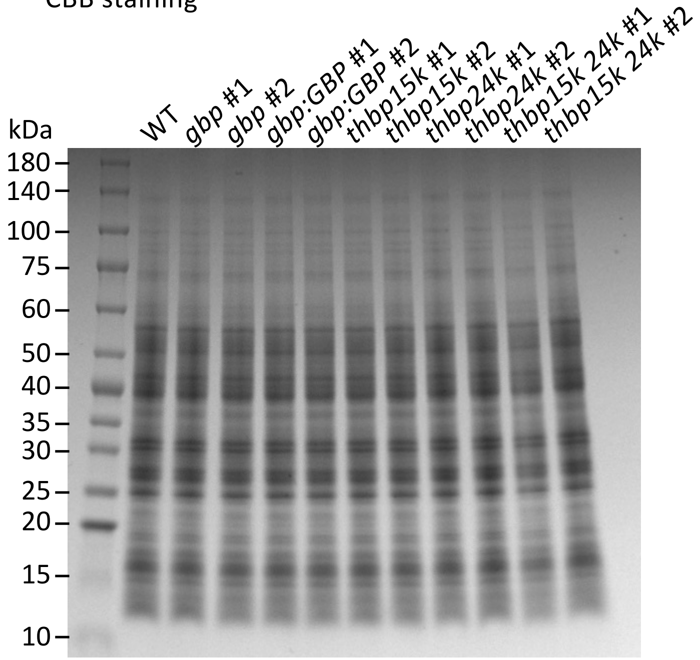


**C**

CBB staining

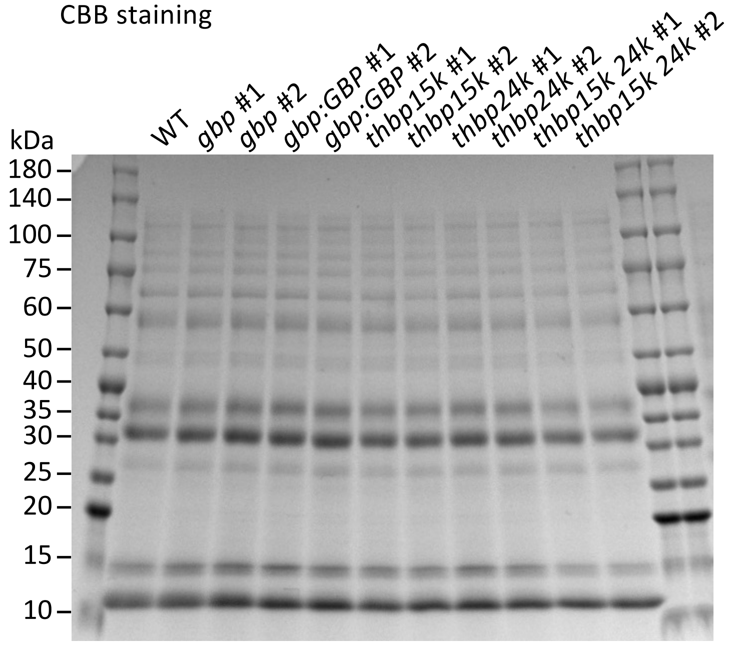

**Figure S5. *M. oryzae* knockout mutants do not produce their corresponding proteins**

**(A)** *M. oryzae* WT and gene-knockout mutant strains were cultured in YG medium containing rice cell wall material for 3 days. The cell wall fraction was used for immunoblot analysis of GBP, and the culture filtrate was used for immunoblot analysis of ThBP24k and ThBP15k. Proteins equivalent to 0.5 mL of *M. oryzae* culture were loaded to SDS-PAGE. Immunoblotting was performed using the corresponding antibodies. **(B, C)** Protein preparations of cell wall fraction **(B)** and culture filtrate fraction **(C)** were stained with CBB to verify equal loading of the protein.

Supplementary Figure S6

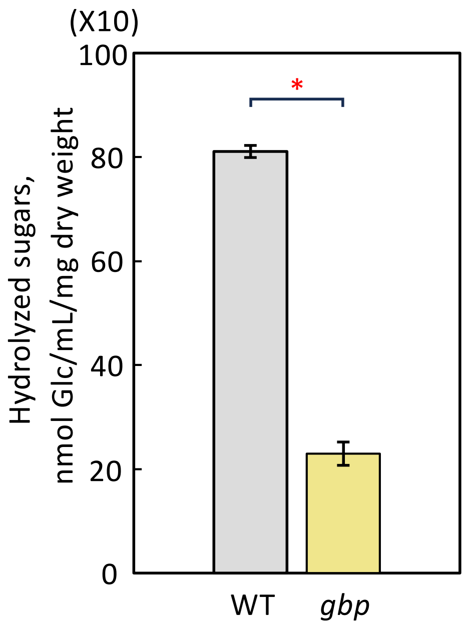


**Figure S6. Susceptibility of cell wall with β-1,3-glucan degrading enzyme**
Cell wall fractions from *M. oryzae* wild-type (WT) and *gbp* mutant strains were treated with *Trichoderma* sp. endo-β-1,3-glucanase (2.0 units/mL) for 18 h at 30°C. The amount of released sugar in the supernatant was determined by measuring the increased reducing power. Means ± SD of three measurements are shown for WT and *gbp*.

Supplementary Figure S7

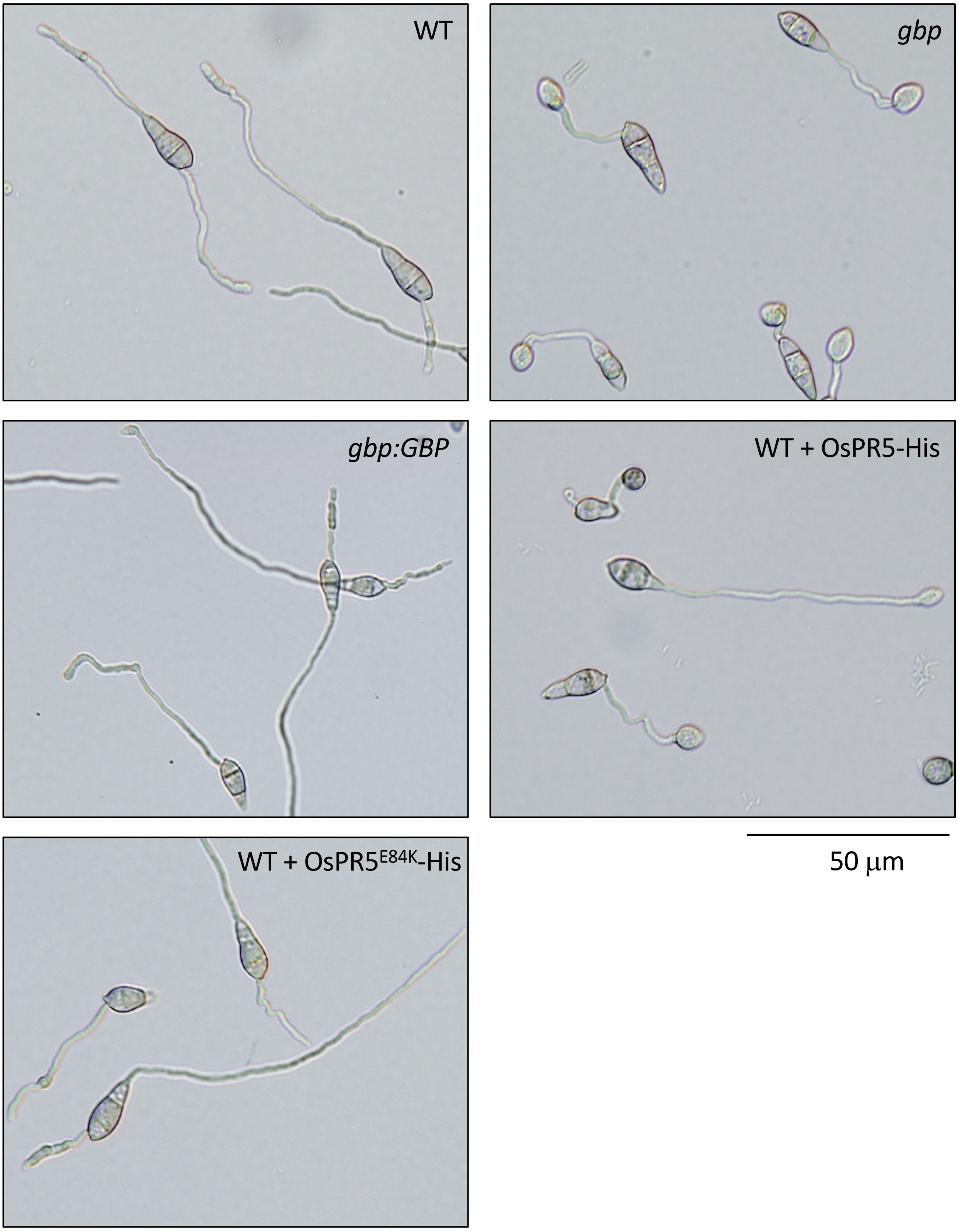


**Figure S7. *GBP* knockout mutants exhibit compromised fungal growth**
Conidia from WT, *gbp*, *gbp:GBP*, and WT *M. oryzae* strains treated with recombinant purified OsPR5-His or OsPR5$^{E84K}$-His were incubated in 50 mM sodium phosphate buffer (pH 5.5) for 7 h. The hyphae were photographed, and their lengths were measured using ImageJ software.

Supplementary Figure S8

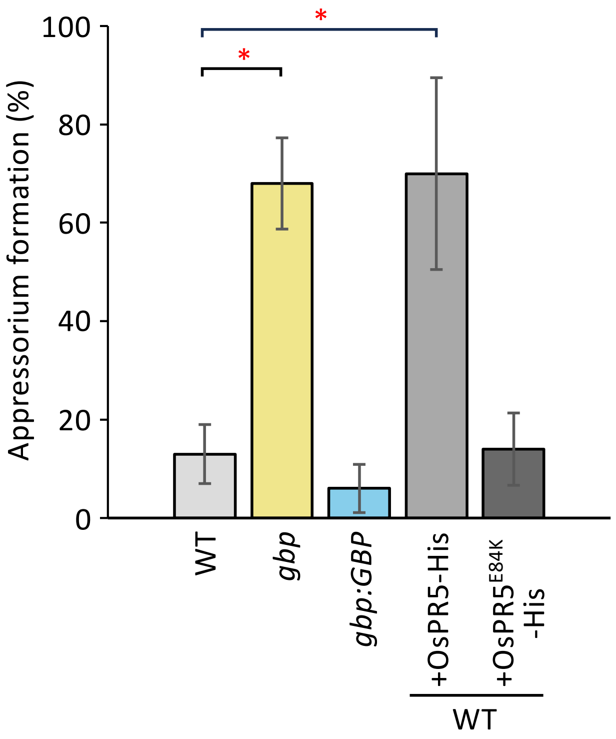


**Figure S8. *M. oryzae* appressorium formation**

Conidia of *M. oryzae* WT, *gbp*, *gbp*:*GBP*, and WT in the presence of recombinant purified OsPR5-His or OsPR5$^{E84K}$-His were incubated in 30 µL of 50 mM sodium phosphate buffer (pH 5.5) with 0.2% (w/v) glucose and 0.05% (w/v) yeast extract on a hydrophobic glass slide at 25˚C for 7 h. Germinated hyphae with or without appressorium were observed under microscope ($n$ = 5, each 20 hyphae observed). Asterisks indicate a significant difference at $P < 0.01$ according to a two-sided Student's $t$-test. Data are shown as means ± SD from five independent determinations.

Supplementary Figure S9

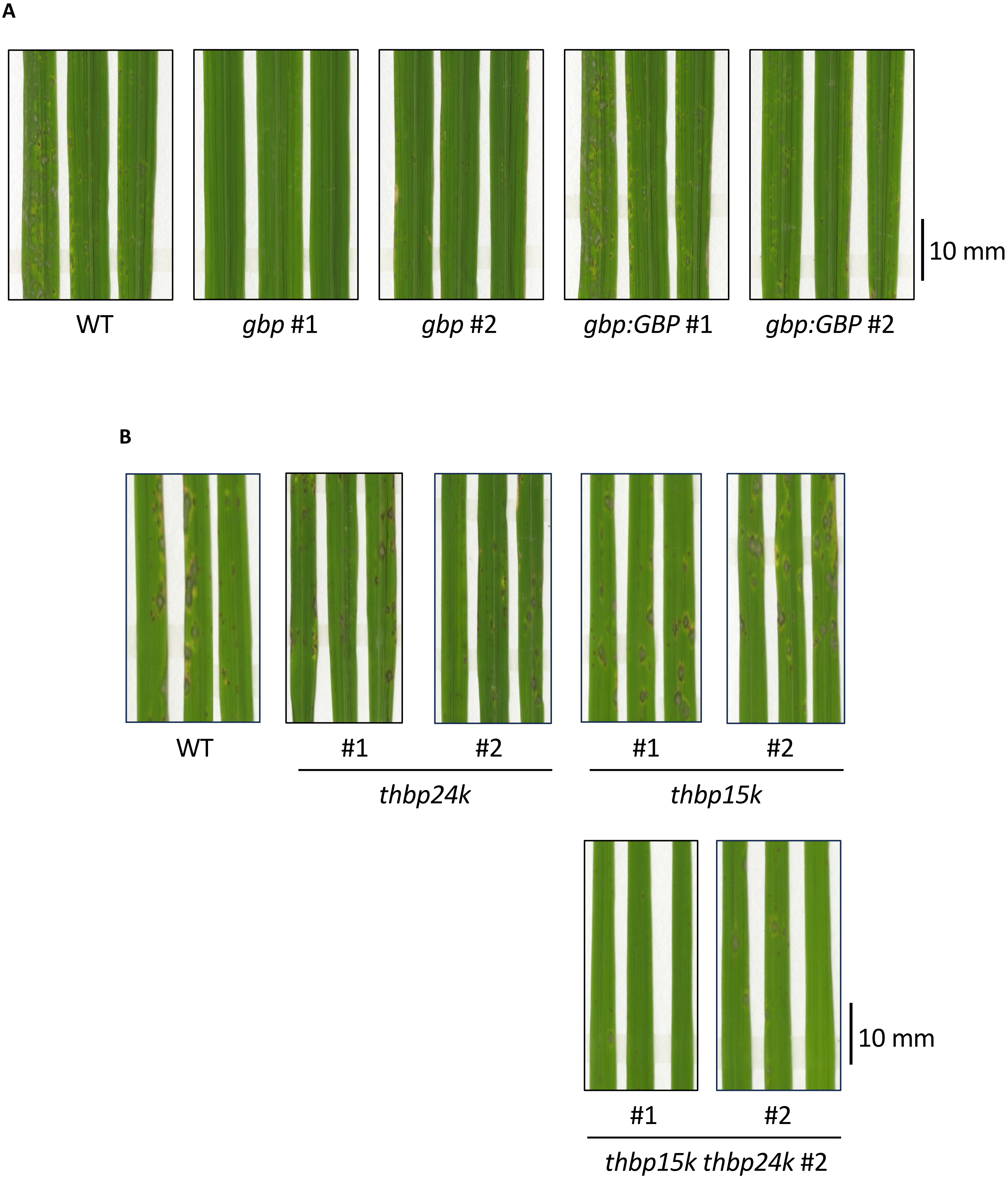

C

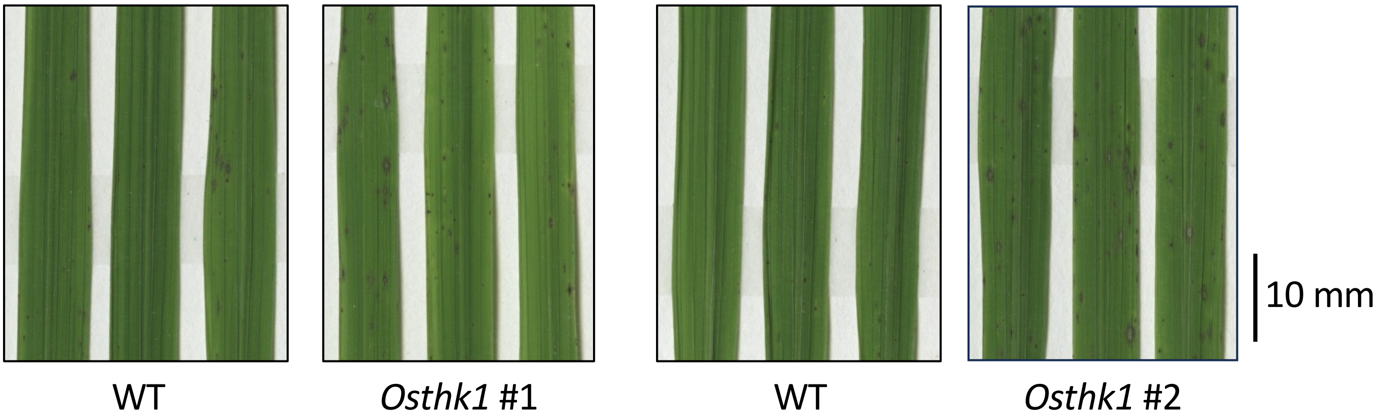


**Figure S9. Apoplastic proteins affect on fungal infection and rice plant thaumatin kinase (Thk1) is required for the defense**
**(A-C)** Representative photographs of inoculated rice leaves as shown in Figure 1G **(A)**, Figure 2B **(B)**, and Figure 4C **(C)**.

#### Supplementary Figure S10

**A**

Binding of OsPR5-Flag protein to the native cell wall of the indicated *M. oryzae* isolates

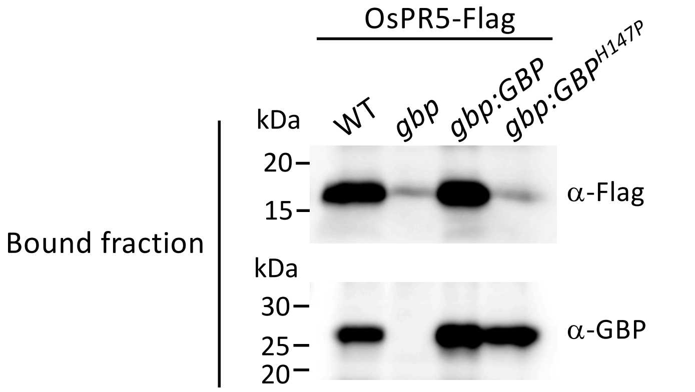


**B**

Binding of OsPR5-Flag protein to the protein-free native cell wall of the indicated *M. oryzae* isolates

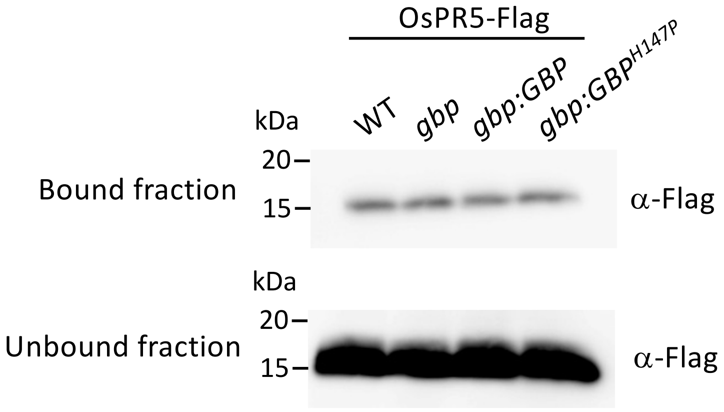


**Figure S10. OsPR5 binds strongly to GBP but binds only weakly to β-1,3-glucan in the *M. oryzae* native cell wall**

**(A)** Testing the binding of OsPR5-Flag to native cell wall fractions prepared from *M. oryzae* WT, *gbp*, *gbp:GBP*, and *gbp:GBP*$^{H147P}$ strains. *M. oryzae* hyphae grown in YG medium for 3 days were sonicated and washed with 50 mM sodium phosphate buffer (pH 6.0). The pellet obtained by centrifugation was used as the native cell wall fraction. OsPR5-Flag produced in *N. benthamiana* leaves was incubated with the native cell wall fraction and washed with the same buffer to remove unbound protein. Bound proteins were eluted with 2% (w/v) SDS containing 5% (v/v) 2-mercaptoethanol and analyzed by immunoblotting using an anti-Flag antibody. **(B)** Testing the binding of OsPR5-Flag to a protein-free native cell wall. The native cell wall fraction was treated with 2% (w/v) SDS containing 5% (v/v) 2-mercaptoethanol to remove bound proteins and washed with 50 mM sodium phosphate buffer (pH 6.0). OsPR5-Flag was incubated with the treated cell wall fraction, and bound proteins were analyzed by immunoblotting using an anti-Flag antibody. OsPR5 does not bind to the GBP$^{H147P}$ variant, as shown in Figure S16. OsPR5-Flag bound to native cell wall fractions from *M. oryzae* WT and *gbp:GBP*, but bound only weakly to the protein-free native cell wall. These results indicate that OsPR5 binds to GBP but binds only weakly to β-1,3-glucan in the *M. oryzae* cell wall.

Supplementary Figure S11

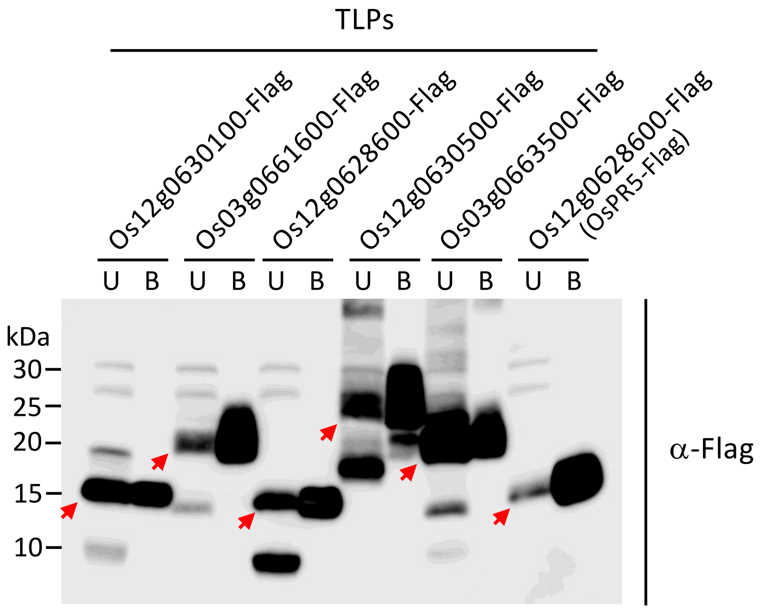


**Figure S11. Various TLPs bind to β-1,3-glucan *in vitro***

Six rice TLPs were produced with a Flag-tag in *N. benthamiana* leaves. TLP-Flag proteins were examined *in vitro* for binding to pure crystalline β-1,3-glucan. Proteins were separated into unbound (U) and bound (B) fractions to pure crystalline β-1,3-glucan by SDS-PAGE and detected by immunoblotting using an anti-Flag antibody.

#### Supplementary Figure S12

**A** OsPR5 vs. GBP (ipTM=0.89, pTM=0.90)

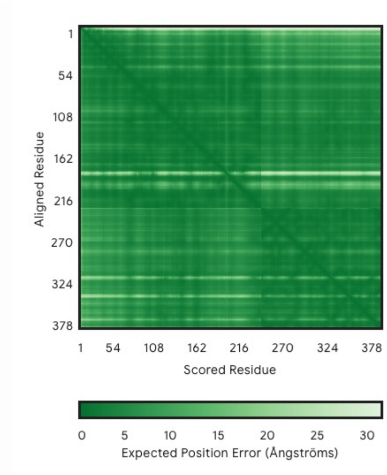


**B** OsPR5 vs. ThBP24k (ipTM=0.88, pTM=0.88)

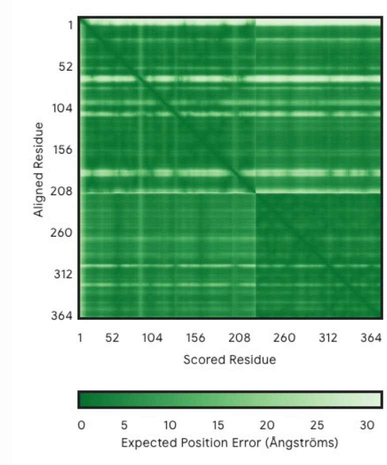


**C** OsPR5 vs. ThBP15k (ipTM=0.89, pTM=0.87)

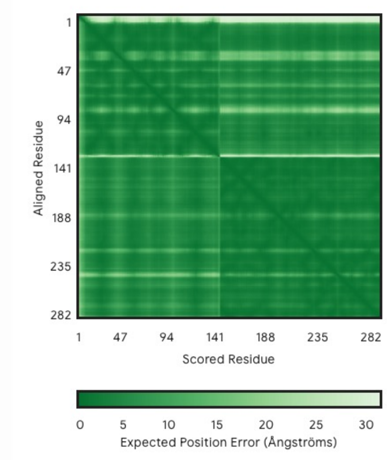


**Figure S12. The predicted 3D structures of the protein–protein complex between OsPR5 and *M. oryzae* proteins are highly reliable**

Predicted ipTM, pTM, and aligned error (right) for the structures of the OsPR5–GBP **(A)**, OsPR5–ThBP24k **(B)**, and OsPR5–ThBP15k **(C)** complexes.

Supplementary Figure S13

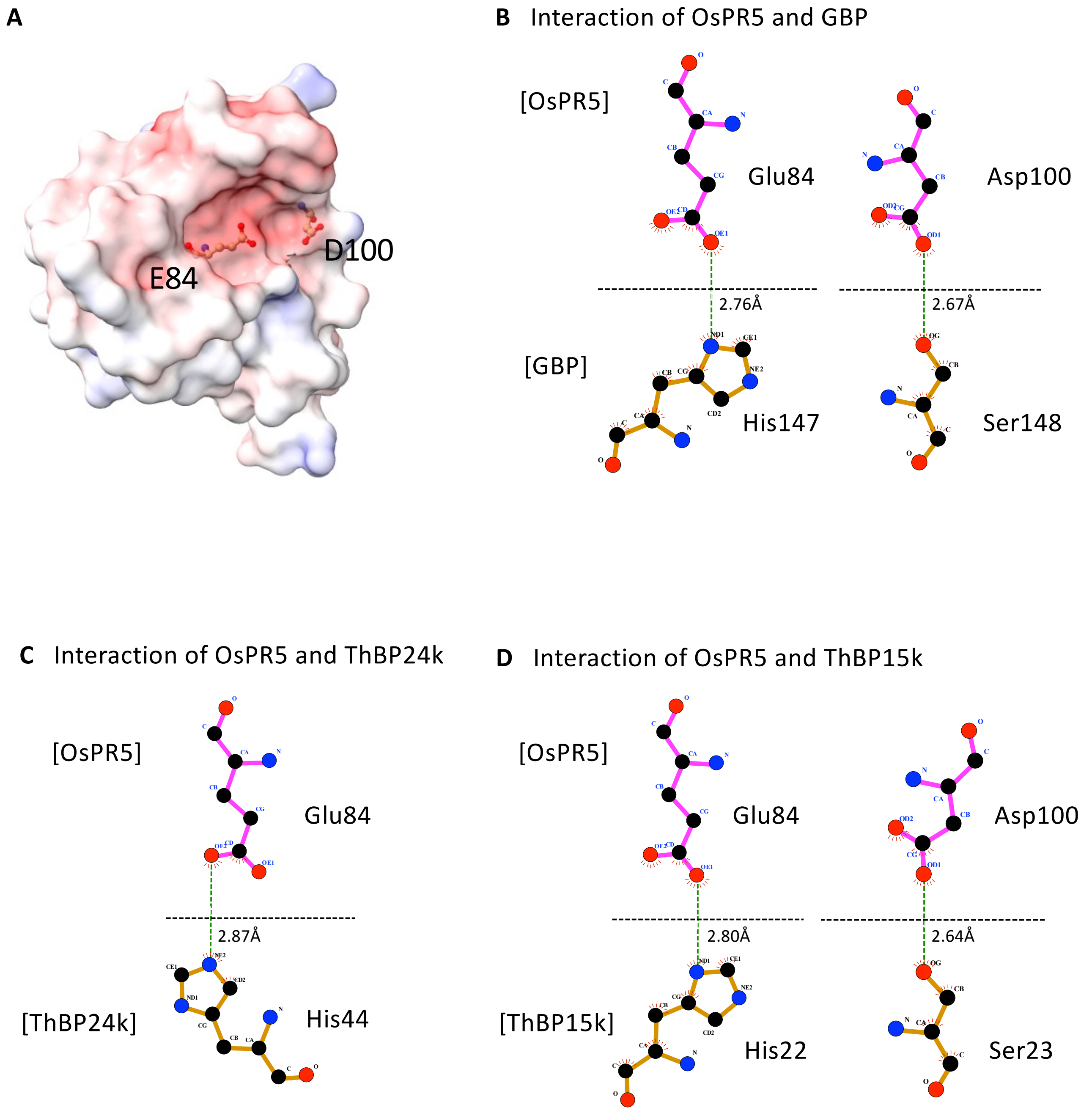


**Figure S13. Amino acid interaction diagrams**

**(A)** Diagram showing the Poisson-Boltzmann electrostatic potential mapped onto the molecular surface of OsPR5. The E84 and D100 amino acid residues of OsPR5 are highlighted as the amino acids in the cleft region with a negative charge. **(B–D)** Amino acid interactions between OsPR5 and GBP **(B)**, OsPR5 and ThBP24k **(C)**, and OsPR5 and ThBP15k **(D)**, visualized using LigPlot+ v.2.2 (https://www.ebi.ac.uk/thornton-srv/software/LigPlus/).

### Supplementary Figure S14

**A**

Phylogenetic tree of GBP homologous proteins

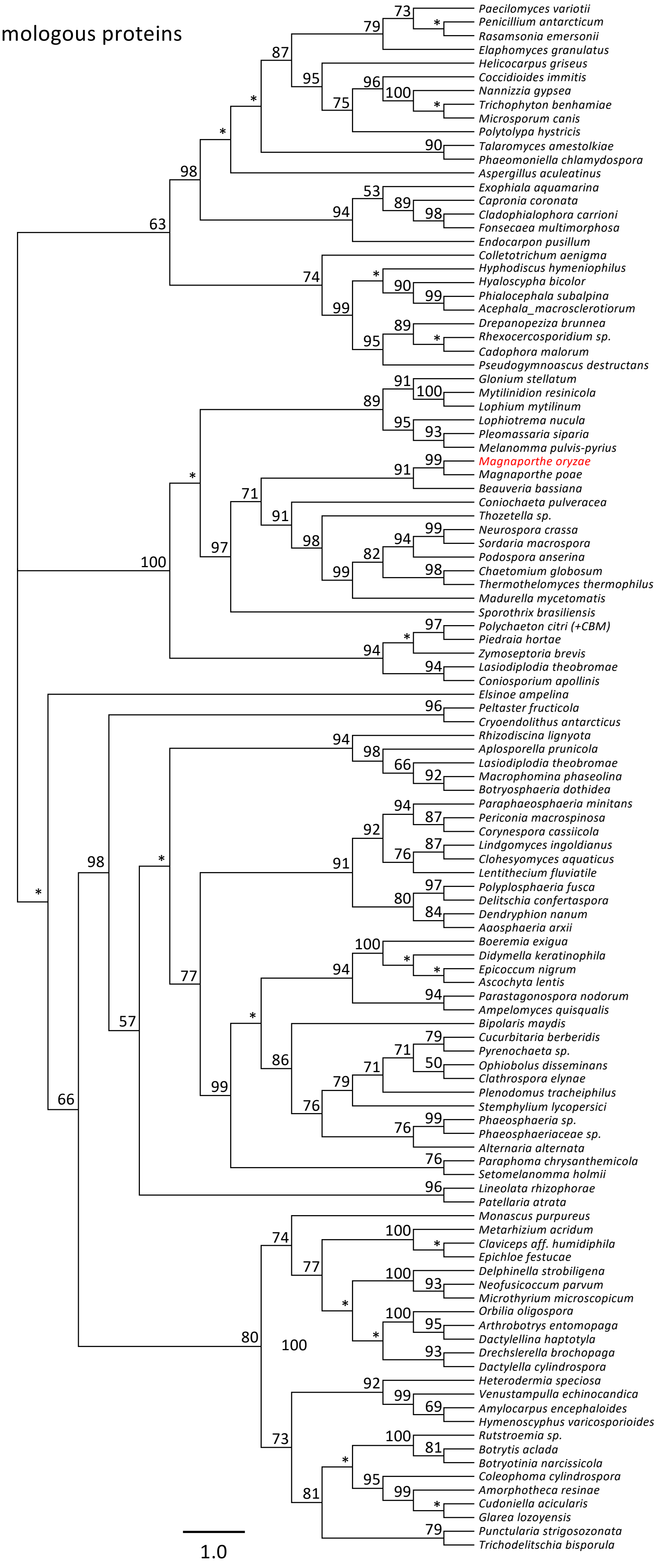

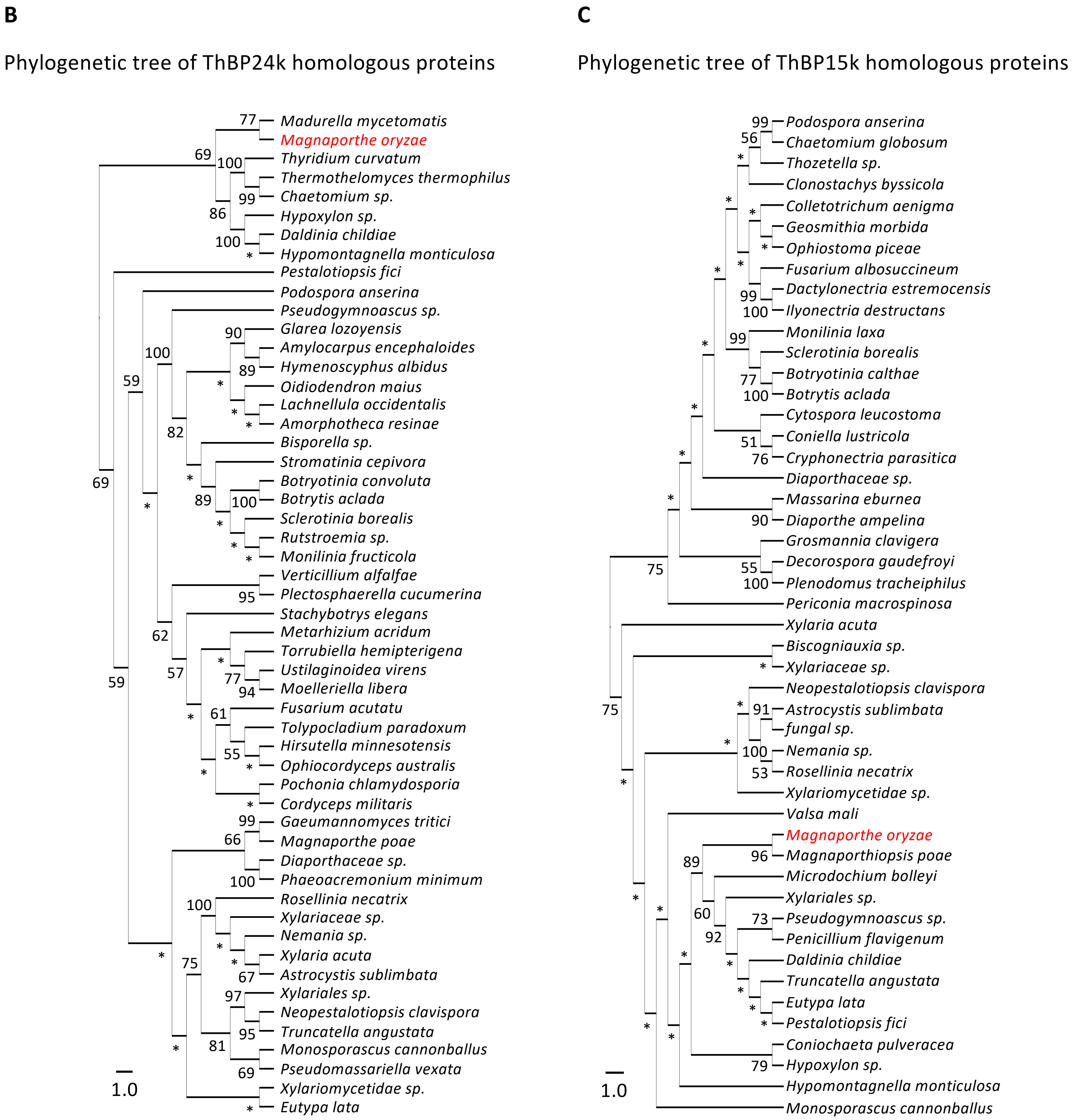


**Figure S14. Phylogenetic trees of GBP, ThBP24k, and ThBP15k homologous proteins**

**(A–C)** Maximum-likelihood trees of GBP homologous proteins **(A)**, ThBP24k homologous proteins **(B)**, and ThBP15k homologous proteins **(C)**. The trees were reconstructed with 1,000 bootstrap replicates. Asterisks indicate support values < 50. The proteins from *M. oryzae* are highlighted in red.

Supplementary Figure S15

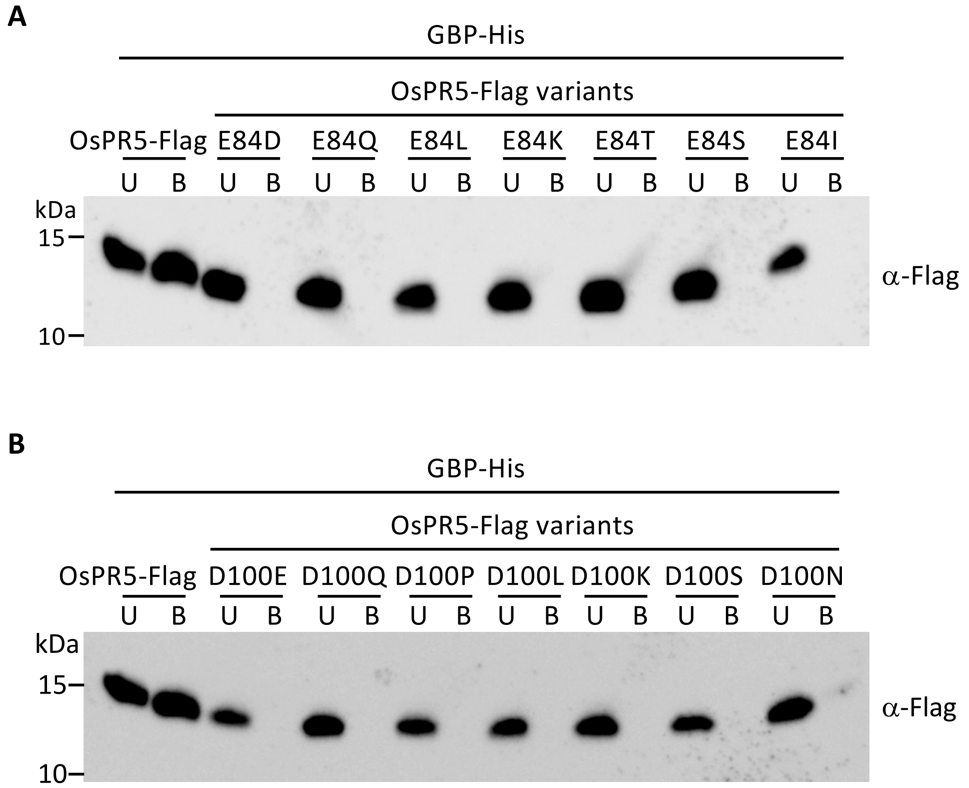


**Figure S15. OsPR5 point mutants with amino acid substitutions at E84 or D100 do not bind to GBP**

**(A, B)** Intact OsPR5-Flag and the indicated OsPR5-Flag variants with amino acid substitutions at E84 **(A)** or D100 **(B)** were produced in *N. benthamiana* leaves. Prepared OsPR5-Flag variants were used for pull-down assays with GBP-His using His-resin. Proteins were separated into unbound (U) and bound (B) fractions and detected by immunoblotting using an anti-Flag antibody.

Supplementary Figure S16

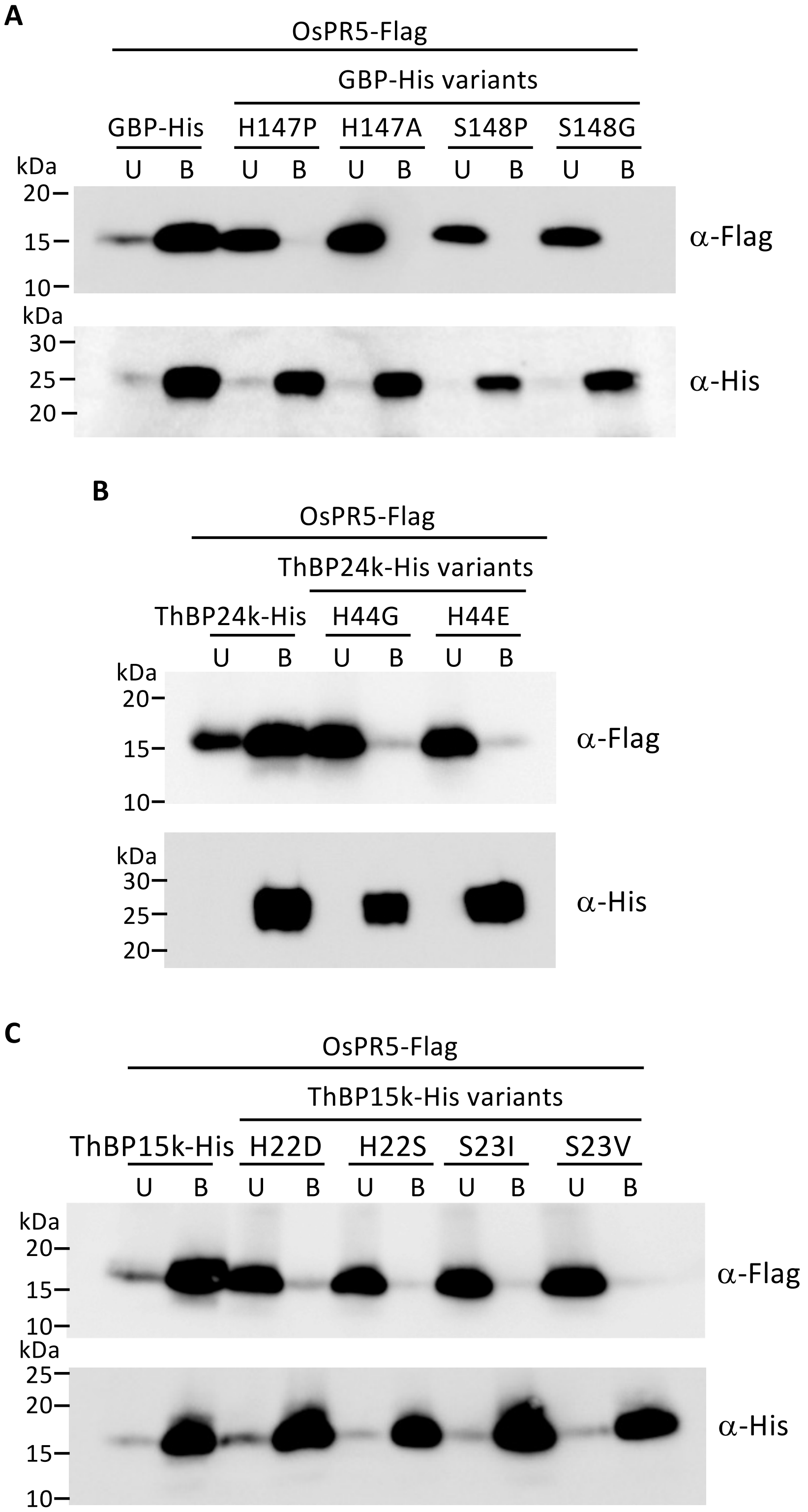


**Figure S16. Amino acid substitutions at the binding sites of *M. oryzae* proteins abolish their binding to OsPR5**

***(A–C)*** *M. oryzae* variant proteins with amino acid substitutions at H147 or S148 of GBP **(A)**, H44 of ThBP24k **(B)**, and H22 or S23 of ThBP15k **(C)** were produced with a His-tag. The proteins were used for pull-down assays with OsPR5-Flag using His-resin. Proteins were separated into unbound (U) and bound (B) fractions and detected by immunoblotting using anti-Flag and anti-His antibodies.

Supplementary Figure S17

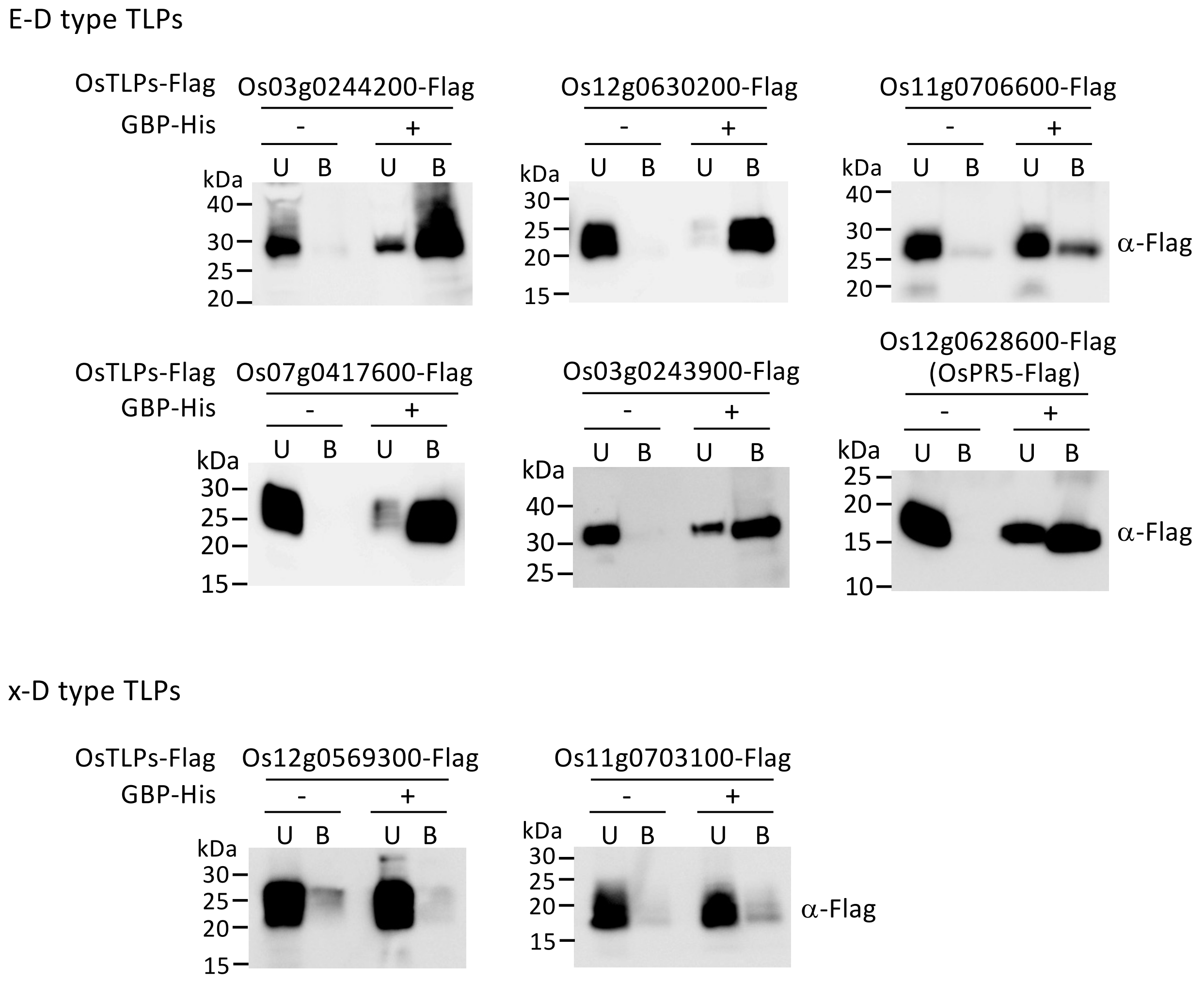


**Figure S17. Various rice E-D type TLPs bind to GBP**

Rice E-D and x-D type TLPs were produced with a Flag-tag in *N. benthamiana* leaves. TLP-Flag proteins were examined for binding to GBP-His by pull-down assays using His-resin. Proteins were separated into unbound (U) and bound (B) fractions from His-resin and detected by immunoblotting using an anti-Flag antibody.

Supplementary Figure S18

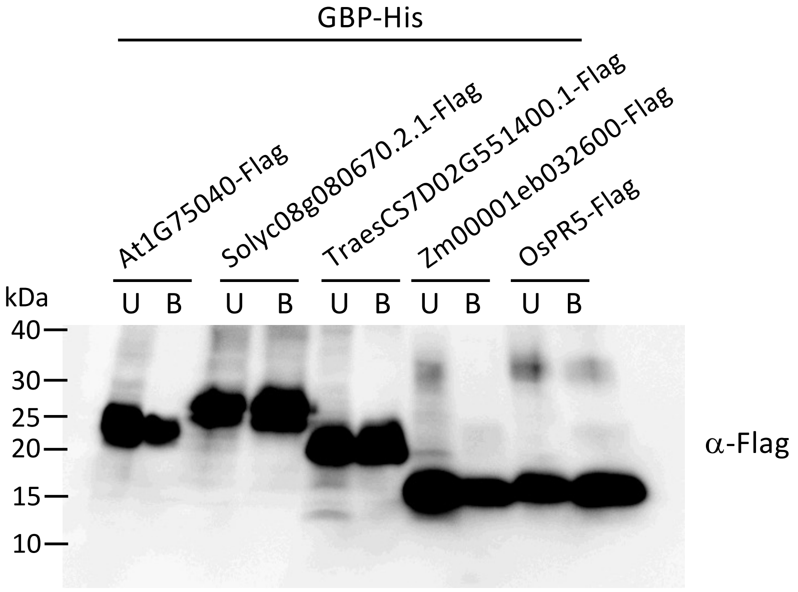


**Figure S18. Various plants possess TLPs that bind to GBP**

E-D type TLPs from Arabidopsis (encoded by At1g75040), tomato (*Solanum lycopersicum* , encoded by Solyc08g080670.2.1), wheat (*Triticum aestivum, encoded by* TraesCS7D02G551400.1), maize (*Zea mays*, encoded by Zm00001eb032600), and rice (OsPR5) were produced with a Flag-tag in *N. benthamiana* leaves. TLP-Flag proteins were examined for binding to GBP-His by pull-down assays using His-resin. Proteins were separated into unbound (U) and bound (B) fractions from His-resin and detected by immunoblotting using an anti-Flag antibody.

Supplementary Figure S19

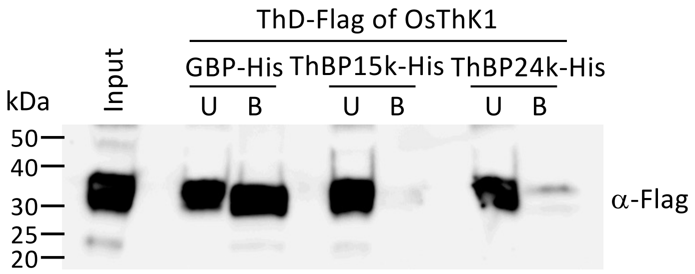


**Figure S19. The thaumatin domain of OsThK1 binds to GBP**

The extracellular region (247 amino acids) containing the thaumatin domain (ThD) of OsThK1 was produced with a Flag-tag in *N. benthamiana* leaves. ThD-Flag was tested for binding to GBP-His, ThBP15k-His, and ThBP24k-His by pull-down assays using His-resin. Proteins were separated into unbound (U) and bound (B) fractions from His-resin and detected by immunoblotting using an anti-Flag antibody.

Supplementary Figure S20

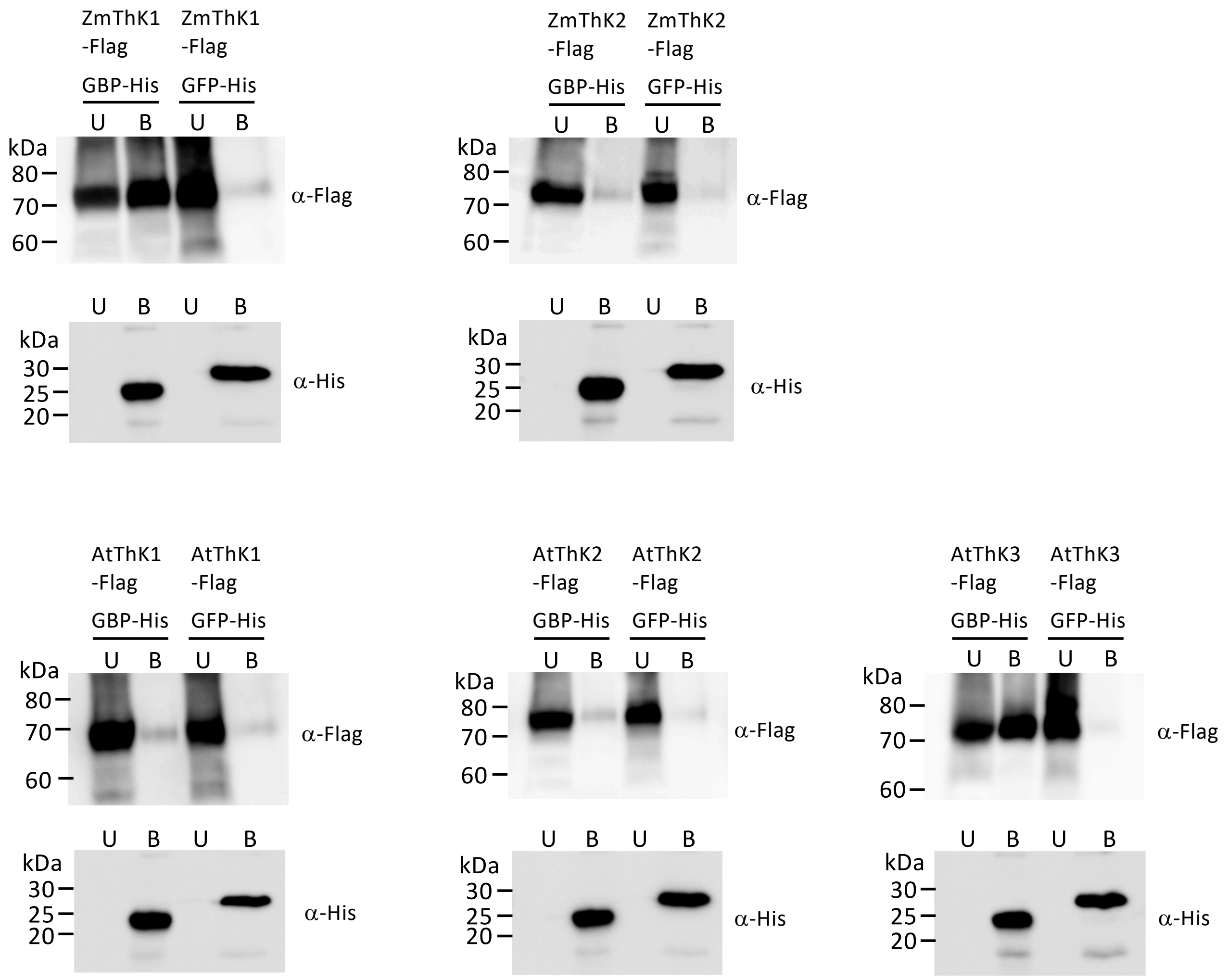


**Figure S20. ThKs from maize and Arabidopsis bind to GBP**

ThKs from maize (ZmThK1; Zm00001eb201520, ZmThK2; Zm00001eb202030) and Arabidopsis (AtThK1; AT1G70250, AtThK2; AT4G18250, AtThK3; AT5G38280) were produced with a Flag-tag in *N. benthamiana* leaves. ThK-Flag proteins were assessed for binding to GBP-His by pull-down assays using His-resin. GFP-His was used as a negative control. Proteins were separated into unbound (U) and bound (B) fractions from His-resin and detected by immunoblotting using anti-Flag and anti-His antibodies.

Supplementary Figure S21

**A**

| Line name | Sequence | Modification |
|---|---|---|
| | (sgRNA.OsThK1-gRNA1) | |
| WT (Moukoto) | CACCTCAGACGCTCGCTGAG-TTCACGGTGGA | |
| *Osthk1* #1 (homo) | CACCTCAGACGCTCGCTGAGTTTCACGGTGGA | 1bp (T) insertion |
| | (sgRNA.OsThK1-gRNA2) | |
| WT (Moukoto) | CTGCCCGGCCAACATCACGTCGCAGTGCCCC | |
| *Osthk1* #2 (homo) | CTGCCCGGCC----TCACGTCGCAGTGCCCC | 4bp (AACA) deletion |

**B**

*Osthk1* #1 (homo)

1bp insertion

5’-CACCTCAGACGCTCGCTGAGTTTCACGGTGGA-3’

3’-GTGGAGTCTGCGAGCGACTCAAAGTGCCACCT-5’

Sequenced

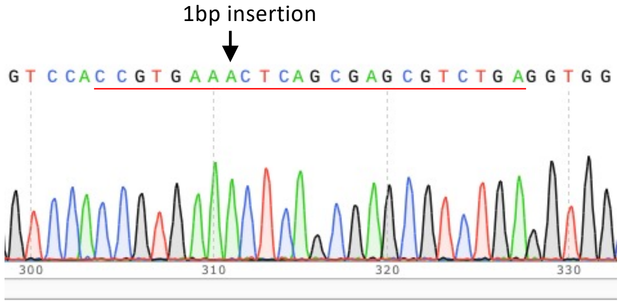


*Osthk1* #2 (homo)

4bp deletion

5’-CTGCCCGGCC----TCACGTCGCAGTGCCCC-3’

3’-GACGGGCCGG----AGTGCAGCGTCACGGGG-5’

Sequenced

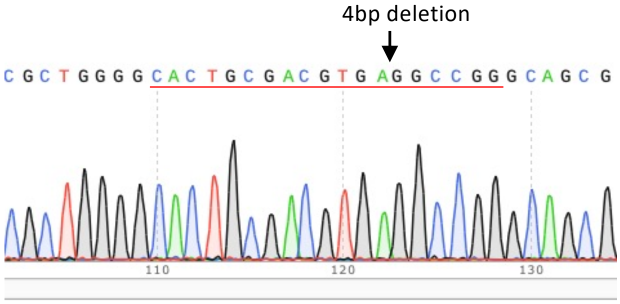


**Figure S21. Verification of DNA mutations in *Osthk1***
**(A)** A table showing locations of targeted DNA sequences used for CRISPR/Cas9 mutagenesis for OsThK1 (red), and the resulting nucleotide changes. PAM sequences are indicated with blue. **(B)** Verification of DNA mutations in genomic DNA in *Osthk1* #1 (upper panel) and #2 (lower panel).

### Supplementary Figure S22

**A**

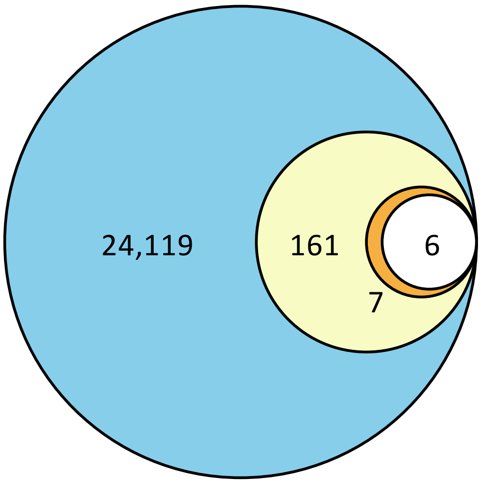


Candidate 6 genes in a downstream of OsThK1

| Gene ID | Annotation |
|---|---|
| Os03g0277700 | DUF26 protein A |
| Os03g0312300 | DUF26 protein B |
| Os03g0700100 | OsThionin38 |
| Os05g0248200 | GH18 chitinase |
| Os01g0847100 | Function unknown protein |
| Os05g0491000 | Calmodulin-like |

**B**

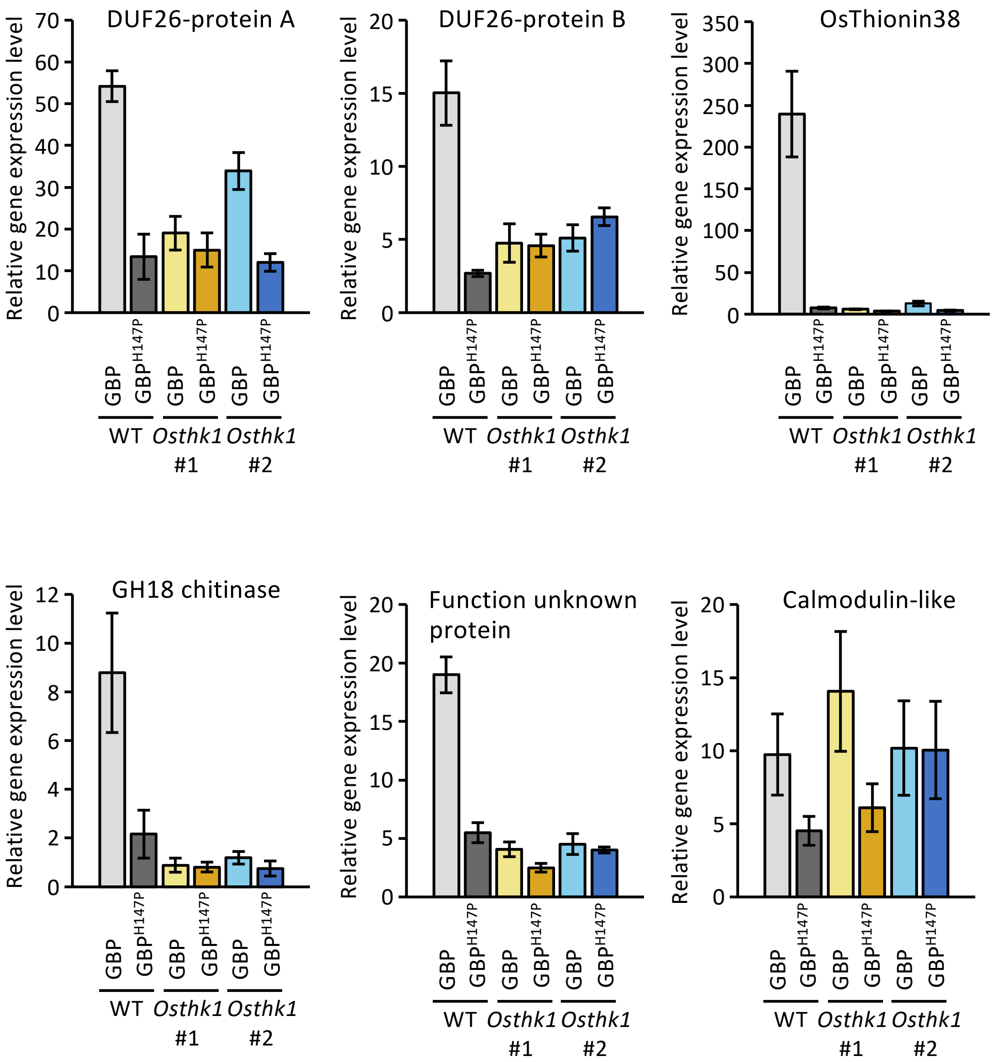

**Figure S22. Gene expression analysis by RNA-seq identifies six defense-related genes downstream of ThK1**

**(A)** Three independent replicates of rice callus prepared from WT and *Osthk1* mutants were treated with GBP, chitin, or 10 mM sodium phosphate buffer (pH 7.0) as control for 6 h. Total RNA extracted from treated callus was used for RNA-seq analysis (Data S2). A total of 24,119 genes with TPM values greater than 3.0 were identified (blue circle). To obtain genes specifically expressed in response to GBP treatment, we analyzed the data as follows: first, we selected the genes more abundantly ( > 3.5 times) expressed in GBP-treated WT cells compared to non-treated WT cells, which resulted in the selection of 161 genes (yellow circle) ; second, focusing on the 161 genes, we selected the genes more abundantly ( > 2 times) expressed in GBP-treated WT cells compared to GBP-treated *Osthk1* cells, resulting in the selection of 7 genes (orange circle). Finally, we selected the genes that show lower ( < 5 times) induction by chitin treatment of WT cells, resulting in 6 genes. These 6 genes were regarded as those functioning downstream of OsThK1(see table). **(B)** Gene expression analysis of candidate genes downstream of OsThK1 was carried out by RT-qPCR using rice callus prepared from WT and *Osthk1* (#1 and #2) treated with purified GBP or $GBP^{H147P}$ for 6 h. *OsACTIN* was used as an internal control to normalize gene expression levels. Gene expression was evaluated as a relative expression level in which the expression levels of untreated rice callus at time 0 was set to a ratio of 1. Data are shown as means ± SD from three independent determinations.

#### Supplementary Figure S23

*Magnaporthe oryzae* GBP amino acid sequence

```
ALAEKCGDAN YDPKSYICHN GNFLCPVVNG EGLSYCNGAC YSKFMYQCNS
                                                    **
GTLSQLPLLE QGSKFTLTAW NPASPSVHGK AIANSGRHWW VGGETSSYCP
        *      *          * **** *                **
DIVKDQGACP PGTVTAMAYS GGMNTLVPGG QGYYLNANWN VEVTQAHSSY
                                 *
IPSGSTVGGL VAYKNGGFIN TNGQPTGWVA CPGGSDGRWN LVAGNASSAD
    *
ALKSCTGINL QVNYLGDSSV PASAWQYI
```

Model structure of GBP–β-1,3-glucan hexaose complex

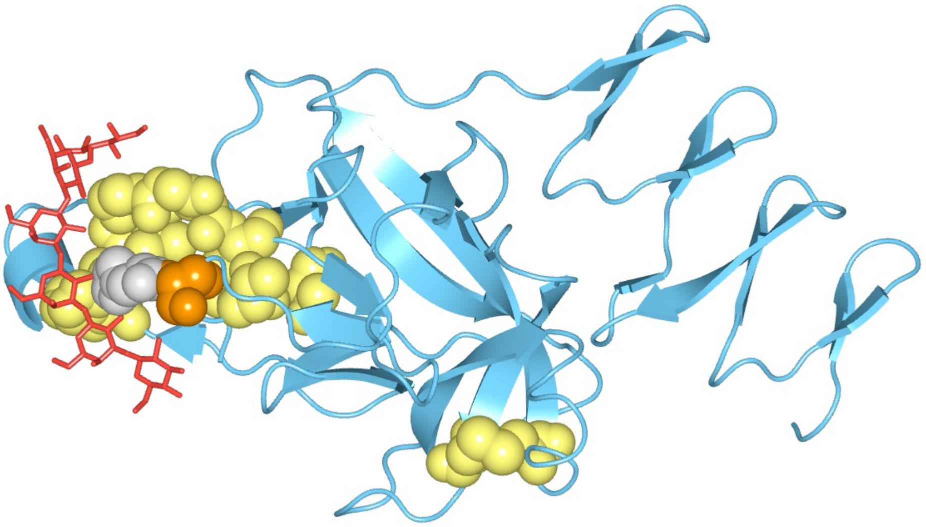

**Figure S23. Highly conserved amino acids possibly form binding surface to β-1,3-glucan.** Amino acid sequences of 113 GBP homologs were aligned using CLUSTALW. Amino acids that are conserved in more than 95% of GBP homologs are indicated with asterisks in *M. oryzae* GBP amino acid sequence and with space-filling balls in the predicted 3D structure of *M. oryzae* GBP. Molecular docking of GBP and β-1,3-glucan hexaose was performed using the Protenix server (protenix-server.com). Residues H147 and S148 of GBP are colored grey and orange, respectively, while β-1,3-glucan hexaose is highlighted in red.

Supplementary Figure S24

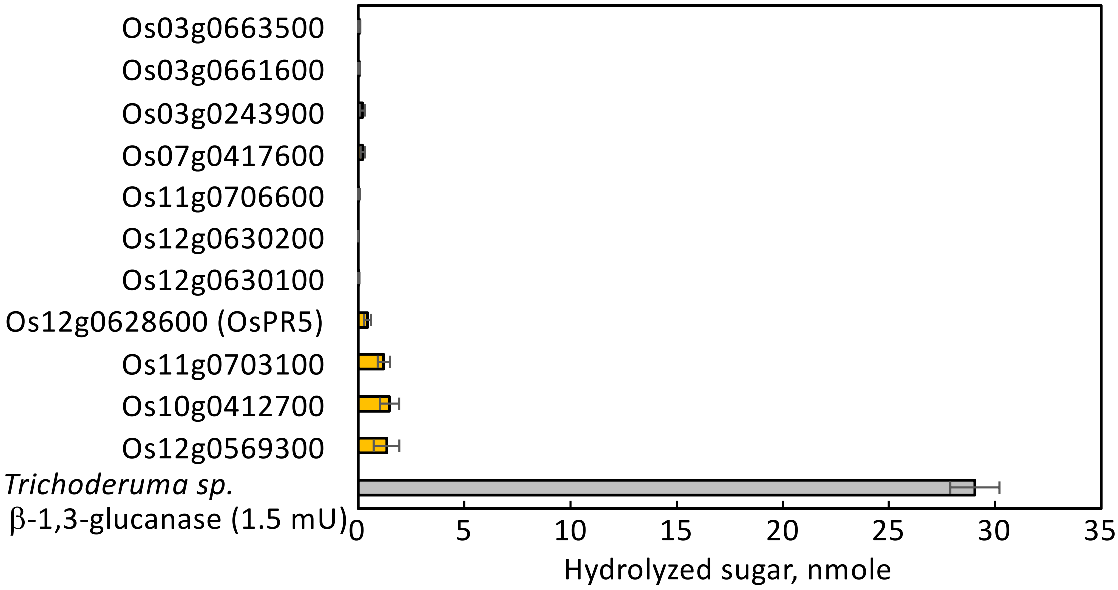


**Figure S24. Rice TLPs show low levels of β-1,3-glucanase activity**

The indicated proteins were produced in *N. benthamiana* leaves with a Flag-tag and purified by affinity chromatography using Flag-agarose. Proteins (0.5 μg) were incubated in 100 mM sodium phosphate buffer (pH 6.0) containing 0.1%, (w/v) laminarin at 30° C for 18 h. Hydrolytic activity was determined by measuring the increase in reducing power. *Trichoderma* sp. β-1,3-glucanase (1.5 mU, Sigma-Aldrich) was used as a control. Data are shown as means ± SD from three independent determinations.

Supplementary Table S1

**Table S1. List of highly abundant TLPs in rice leaves infected with *M. oryzae***

Rice proteins were extracted from rice leaves 2 days after *M. oryzae* inoculation and subjected to SDS-PAGE, followed by silver staining. SDS-PAGE gel slices containing proteins in the 10–40 kDa range were excised and analyzed by LC–MS/MS. Rice leaves treated with $ddH_2O$ were used as a control. TLPs detected by LC–MS/MS are listed.

| | Scores detected by LC-MS/MS | |
|---|---|---|
| Gene ID | *M. oryzae* treatment | Water-treatment |
| Os12g0628600 (OsPR5) | 580 | 157 |
| Os12g0568900 | 240 | 59 |
| Os03g0663500 | 165 | - |
| Os03g0661600 | 72 | - |
| Os12g0630100 | 58 | - |

**Table S2. TLPs and ThKs are widely distributed across various plant species**

Genes encoding TLPs and ThKs were searched using keyword 'thaumatin' and pfam; PF00314 in plant genome databases registered on Ensembl Plants, and Figshare.

| | TLPs | ThKs | |
|---|---|---|---|
| *Galdieria sulphuraria* | 0 | 0 | *Cyanidiales, Galdieriaceae* |
| *Chondrus crispus* | 0 | 0 | *Gigartinales, Gigartinaceae* |
| *Ostreococcuslucimarinus* | 0 | 0 | *Mamiellales, Bathycoccaceae* |
| *Chara braunii* | 0 | 0 | *Charales, Characeae* |
| *Chlamydomonas reinhardtii* | 1 | 0 | *Volvocales, Chlamydomonadaceae* |
| *Physcomitrium patens* | 7 | 0 | *Funariales, Funariaceae* |
| *Marchantia polymorpha* | 22 | 0 | *Marchantiales, Marchantiaceae* |
| *Sellaginella moellendorffii* | 19 | 0 | *Selaginellales, Selaginellaceae* |
| *Alsophila spinulosa* | 58 | 1 | *Cyatheales, Cyatheaceae* |
| Aegilops tauschii | 90 | 7 | |
| *Oryza sativa* | 30 | 2 | |
| *Hordeum vulgare* | 40 | 4 | |
| *Setaria italica* | 41 | 2 | |
| *Zea mays* | 53 | 2 | |
| *Brachypodium distachyon* | 39 | 3 | *Poales, Poaceae* |
| *Panicum hallii* | 38 | 2 | |
| *Lolium_perenne* | 51 | 7 | |
| *Eragrostis curvula* | 46 | 1 | |
| *Echinochloa crus-galli* | 55 | 0 | |
| *Degitaria exilis* | 54 | 0 | |
| *Ananas comosus* | 22 | 0 | *Poales, Bromeliaceae* |
| *Asparagus officinalis* | 8 | 0 | *Asparagales, Asparagaceae* |
| *Musa acuminata* | 51 | 0 | *Zingiberales, Musaceae* |
| *Dioscorea rotundata* | 44 | 0 | *Dioscoreales, Dioscoreaceae* |
| *Actinidia chinensis* | 38 | 0 | *Ericales, Actinidiaceae* |
| *Arabidopsis thaliana* | 21 | 4 | |
| *Brassica napus* | 93 | 10 | *Brassicales, Brassicaceae* |
| *Eutrema salsugineum* | 28 | 5 | |
| *Arabis alpina* | 22 | 3 | |
| *Gossypium raimondii* | 53 | 1 | *Malvales, Malvaceae* |
| *Medicago truncatula* | 49 | 3 | |
| *Pisum sativum* | 29 | 2 | |
| *Trifolium pratense* | 36 | 3 | |
| *Vigna angularis* | 34 | 0 | *Fabales, Fabaceae* |
| *Glycine max* | 64 | 1 | |
| *Lupinus angustifolius* | 29 | 1 | |
| *Phaseolus vulgaris* | 28 | 0 | |
| *Ipomoea triloba* | 36 | 0 | *Solanales, Convolvulaceae* |
| *Solanum lycopersicum* | 27 | 0 | |
| *Nicotiana attenuata* | 28 | 0 | *Solanales, Solanaceae* |
| *Capsicum annuum* | 31 | 0 | |
| *Helianthus annuus* | 49 | 3 | |
| *Lactuca sativa* | 22 | 4 | *Asterales, Asteraceae* |
| *Cynara cardunculus* | 24 | 2 | |
| *Rosa chinensis* | 28 | 1 | |
| *Malus domestica* | 35 | 0 | *Rosales, Rosaceae* |
| *Theobroma cacao* | 26 | 0 | |
| *Prunus avium* | 25 | 0 | |
| *Ficus carica* | 44 | 0 | *Rosales, Moraceae* |
| *Olea europaea* | 45 | 0 | *Lamiales, Oleaceae* |
| *Sesamum indicum* | 27 | 2 | *Lamiales, Pedaliaceae* |
| *Manihot esculenta* | 36 | 0 | *Malpighiales,Euphorbiaceae* |
| *Populus trichocarpa* | 42 | 7 | *Malpighiales, Salicaceae* |
| *Beta vulgaris* | 22 | 0 | *Caryophyllales, Amaranthaceae* |
| *Chenopodium quinoa* | 31 | 2 | |
| *Quercus lobata* | 53 | 0 | *Fagales, Fagaceae* |
| *Citrullus lanatus* | 29 | 0 | *Cucurbitales, Cucurbitaceae* |
| *Cucumis sativus* | 32 | 0 | |
| *Coffea canephora* | 26 | 0 | *Rubiales, Rubiaceae* |
| *Corylus avellana* | 61 | 0 | *Fagales, Betulaceae* |
| *Daucus carota* | 27 | 0 | *Apiales, Apiaceae* |
| *Amborella trichopoda* | 16 | 0 | *Amborellales, Amborellaceae* |

Supplementary Table S3

**Table S3. Verification of DNA sequences in putative off-target sites**

Genes with 70% identity to the sgRNA target DNA sequences were searched in the RAP-DB database by BLAST. The genes listed were amplified by PCR, and their sequences were verified by Sanger sequencing to ensure no off-target mutations.

| Gene ID | Annotation | Identity |
|---|---|---|
| Genes with identity to Target DNA sequence 1 (5'-tcagacgctcgctgagttcacgg-3') | | |
| (1) Os01g0113350-00 | Thaumatin-kinase | 19/23 (83 %) |
| (2) Os02g0755600-01 | UDP-glucuronosyl/UDP-glucosyltransferase | 17/23 (74 %) |
| Genes with identity to Target DNA sequence 2 (5'-ccggccaacatcacgtcgcagtg-3') | | |
| (1) Os01g0113350-00 | Thaumatin-kinase | 20/23 (87 %) |
| (2) Os03g0243700-01 | GH10 Xylanase | 17/23 (74 %) |
| (3) Os03g0158700-01 | subtilisin-like protease SBT3.18 | 17/23 (74 %) |
| (4) Os09g0484900-01 | Tonoplast dicarboxylate transporter | 16/23 (70 %) |

#### Supplementary Table S4

**Table S4. List of primers used in this study**

1) Primers for full-length DNA amplification

| Gene ID | | Forward DNA primer | Reverse DNA primer |
|---|---|---|---|
| *Os12g0628600* | OsPR5 | ATGGCGTCTCCGGCCACCTCTTCCG | TTATGGGCAGAAGACGACTTGGTAG |
| *Os12g0630100* | OsTLP | ATGGCGTCAGCTCCGGCCGCCTCTTC | TCATGGGCAGAAGACGACTCGGTAG |
| *Os03g0661600* | OsTLP | ATGGCGCCTTCCCTCGCCACCTCTTC | TCATGGGCAGAAGACGACCTGGTAG |
| *Os03g0663500* | OsTLP | ATGGCAGCTCCCGCCATCCTCCGCC | TCATGGGCAGAAGACGACCTGGTAG |
| *Os12g0630200* | OsTLP | ATGGCCAATACTCGCGTCTTCGTCC | TTAAGGGCACATGACGATCTGGTAA |
| *Os07g0417600* | OsTLP | ATGGTTGCTATGAGAGGAGCCTCGC | TCAAGGGCAGAAGGTGATTTGGTAG |
| *Os03g0244200* | OsTLP | ATGTCACAGTGCCCAACTCATGTCG | CTAGATCATGGCGGCAACGAGGGC |
| *Os03g0243900* | OsTLP | ATGATGGGAATTCAGAGAATTTGCAT | CTATAGCCGCGGCAGGTGGAGCGTG |
| *Os12g0568900* | OsTLP | ATGGCCAGCGCCCTCGCCTTCGTCG | CTAGCTGGAGGCGACGACGGTACGG |
| *Os01g0113650* | OsThK1 | ATGGCAGCGGGGAGTACCATCAGTAC | TCACCTGGGGCCTGATAAATTAGATG |
| *At1g75040* | AtPR5 | ATGGCAAATATCTCCAGTATTCACAT | AGGGCAGAAAGTGATTTCGTAGTTAG |
| *At5g38280* | AtThK | ATGGTTGAAGGGTTCTCATTGTCATTG | TTATATTAACGTTTCTCTCTTTCCAAG |
| *Solyc08g080670* | SlPR5 | ATGAGTCATTTGACAACTTGTTTAG | TTAAGCCACTTCAAGAGTATTTGTAG |
| *TraesCS7D02G551400* | TaPR5 | ATGGCGACCTCCCCGGTGCTCTTCC | TCATGGACAGAAGGTGATCTGGTAG |
| *Zm00001eb032600* | ZmPR5 | ATGGCCGCCGCCTCCTCGTCCTCGG | TCACGGGCAGAAGGTGACTTGGTAG |
| *Zm00001eb202030* | ZmThK | ATGGGAGCGAGGCGTAGTAGCACTTG | TCACCTTGGGCCTGACACTTCTGAGG |
| *MGG_05232* | GBP | ATGAAGTCCCTCTTCCTCACACTCG | TTAAATGTATTGCCAAGCCGAAGCC |
| *MGG_01944* | ThBP24k | ATGCTCACCAAATCAATCCTCCTCGC | CTACAAACTCCAGTCAGCTCCAGCAAC |
| *MGG_10456* | ThBP15k | ATGCAGTTCATCTCTACCTTCCTCGC | TTACTTCCTGTCGCACTCAAAGATGG |

2) DNA fragment for RNAi plasmid construct

| Gene ID | | Forward DNA primer | Reverse DNA primer |
|---|---|---|---|
| *Os12g0628600* | OsPR5 | GACCCTCGCCGCCGGTGGCGCCAACG | GACAGGGCGCCGGCGCAGTCGCCGGTG |

3) DNA primers for generating mutant proteins

| Gene ID | | Forward DNA primer | Reverse DNA primer |
|---|---|---|---|
| *Os12g0628600* | OsPR5$^{E84x}$ | CCGCTGACGCTGGCGnnnTTCACC ATCGGCGGCAGCCAGGAC | CGCCAGCGTCAGCGGCTTCTGGCCG |
| *Os12g0628600* | OsPR5$^{D100x}$ | GACCTGTCGGTGATCnnnGGCTAC AACGTCGCCATGAGCTTC | GATCACCGACAGGTCGTAGAAGTCC |
| *MGG_05232* | GBP$^{H147x}$ | GAGGTCACCCAGGCCnnnTCGAGC TACATCCCCTCGGGTAGC | GGCCTGGGTGACCTCGACGTTCCAG |
| *MGG_05232* | GBP$^{S148x}$ | GTCACCCAGGCCCACnnnAGCTAC ATCCCCTCGGGTAGCACC | GTGGGCCTGGGTGACCTCGACGTTC |
| *MGG_01944* | ThBP24k$^{H44x}$ | CAGGTGACGGGGATCnnnATCGG GCCGCCCTACAACCGCGCCG | GATCCCCGTCACCTGGAAGTGGTTG |
| *MGG_10456* | ThBP15k$^{H22x}$ | GCCGCCTGCATCCCTnnnAGCGTG CGCTGCACCTACTCGTTC | AGGGATGCAGGCGGCCTGGAAGGAC |
| *MGG_10456* | ThBP15k$^{S23x}$ | GCCTGCATCCCTCACnnnGTGCGC TGCACCTACTCGTTC | GTGAGGGATGCAGGCGGCCTGGAAG |

4) DNA primers for qRT-PCR

| Gene ID | | Forward DNA primer | Reverse DNA primer |
|---|---|---|---|
| *Os03g0718100* | OsACTIN | AACTGGGACGACATGGAGAAA | AGCAACACGCAGCTCATTGT |
| *MGG_03982* | MoACTIN | AACGCCCCCGCTTTCTAC | GGTACGACCCGAAGCGTAAA |
| *Os12g0628600* | | TGCATCATCTCATGCATGTAAC | GATTATCGATCAAGGTGTCG |
| *Os12g0630100* | | CTCCGACGATTCCAAGCTGCG | CTGGATCACGTTCGTTTTATTG |
| *Os12g0663500* | | TGATCGATCGATCTCTGATG | TATTCTTAGGTACGTATATC |
| *Os03g0300400* | OsPR10 | GACGTCGCACATCAAGGTGG | ACTCCTTAGCCTTGGTGATC |
| *Os03g0277700* | DUF26-protein | GCGGTTCGAGAACGAGAATT | CCATGATCACCCCGTTGTC |
| *Os03g0312300* | DUF26-protein | ATCGGACTCCAGGATGTGGTAT | CCAATGAAGTTGGCGTTCTTG |
| *Os03g0700100* | OsThionin38 | GGCGACCCCTCCAACAA | ACAGTAGTTGACGCATTTGGAGTAGT |
| *Os05g0248200* | GH18 Chitinase | CAACGGCATCGACTTCTTCAT | AGCTTGGCTAGCTTGTTGTAGTTCT |
| *Os03g0300400* | Calmodulin-like | TGACACAGCAATTAGGCGAAA | AGCGACGAGGACGGTCGCATC |
| *Os03g0300400* | Function unknown protein | AAGGCATCATCCCGTTCATC | TCAAGGCGGTAGCGCAGTAC |
| *MGG_05232* | GBP | TGGCCTCGTGGCGTACA | GTCGGCTGGCCGTTTGT |
| *MGG_01944* | ThBP24k | GCTCGCTCAGTGCGATGAG | CGTGCGAGAACCAGTTCCA |
| *MGG_10456* | ThBP15k | CCTTCTTCGTCAAGTCCTTCCA | CCGAGGTGATGTTGAACGAGTA |
