## Supplementary material for "Multipartite coevolution shapes plant apoplastic immunity against rice blast fungus": Data S2

(1)Amino acid sequences of 113 GBP homologous proteins

Amino acid sequences used to construct WebLogo are shown in red.

>*Aaosphaeria arxii*

MKSVVALSLFAAGALAQAPAYFSVLSTRSASPVHLLPLTARGGKFFLGGPGPSSYCPPQVGDGCPSGNTTVFAGGEETLSLGVVVPGGQQVYVGPDGNLSYTSPHSAYKPEGSIVDGWSKGSTGTFGTLVWENGLVACPAGEGEGYQVFGQIEGVERDPACLGFNAITANATDAGAWQY

>*Acephala macrosclerotiorum*

MFSLRNALALTLTFAISTAIADTFTLTASLPASPLDGLPVNAAGEAFSLGGSPSFYCPTSVSSACPNTTETIFAAELTALDVEVPGGQQVYVQANGALGFTQAHSASVPPGATIGGFVNVTVASDCHAPWEVITWKSPDGSTAGGIEACSEVPADTNGTAVYKVYAVTPGFNQTGCVKLAGLLASYLGNGAPFGAWQYI

>*Alternaria alternata*

MKYTVALSALAASAMATTVTVGQPSATPTPPANDNGYFGVISSRSASPIHLLPLTARGNKFYLGGGPPSSYCPVESVGAANCPPGNTTVLAGGDQTLSLGVVVPGGQQVYVAPDGALSYTTAHSAAIPAGSEQTGFSRTAPANGNAFGYLNFDTGFVACPAANVTADGYQVFGQVASGATFGDDCLGFSALTVAVDQPGAWQY

>*Amorphotheca resinae*

MHTKHFLATALAGLAAASPIRRADINTFELVAIKSGSPIQNSNIHASGSKFWIGKPTATYCPEQVGSNCPAGDQTAVSQGSSSNTLSMSVVVPGGQLVYVQTDGQVGYTIAHSASIPQGALTSPFEYTPPTDGEPLGSLTFNEKSFSACPTSDEGVYQIYANVDGFTRTDCIGILMGTSEFDETAAWEYT

>*Ampelomyces quisqualis*

MKYAVALNALVSTVLAASPCSAPAPSASSAPSGPSYFTVISARSASPVHLQPLTARGGKFYLGGGPPSSYCPAAQVGAACPSGNTTTLAGGEGTLSLGVIVPGGQQVYVAPDGALSYTVAHSASVPEGSIRDQFSVEEPSGTNSFGYLNFETGFVACPGTGADTGYQVFGQVDGATFGPECLGFNALTSQVDGPDAWQY

>*Amylocarpus encephaloides*

MKSVIAFTTILGAAVAQEIAPPAGSFVLGAWSPGSGLPGSKINANGGSFWLGKDPSSSCPAVEGLDCTAFPGTRTVLVGGNDTLSLDVAVPGGQQVYVGPNGALGYTPPHSSYMPTGSVTTGFHRFRSQSGGAPVVMNFQGNSFLACPDATNSGVYQIFAHLVVTDGGNCTFFQMRTYTAEGINAWQY

>*Aplosporella prunicola*

MKTAAAATLLAAVPAVLGQNKPFEGLTVRSGSDFQYATITQAGLSFWVGKDTESYCPEASVPIDCPPGNVTAFVGGEGTLSMDVEVPGGQQVYIAPTGELKVTQAHSASMPEGSIQDGFHKSESNSLGFLHWTNGFLACPAANSTDGYPYQIYASVDGLKRTDCLGFDFLTSNYTGTGAAAWQYA

>*Arthrobotrys entomopaga*

MKFTLATIIAAASAVSAAPVAETPLALTMMSLRSASPIHFGSVTASGQKFYIGLEHPSSYCPSPPVSPEDCPSGNYTSIGWGTGGPYMNVMVPGGQQLYIAPDHSLSYTQAHSAYMPEGSTTEGFILGPKQENTLRSISHTSGGFLACPVKANQGPWKVFVGTVADADTPSGSSADCLGFSNNGVEFTAERFGAWQY

>*Ascochyta lentis*

MKYAVAFASLVAGALAGECYAPAPPAYSAPPAPTPTPAPGPGYFAVISSRSASPVHLLPLTARGGKFYLGGGPPSSYCPENIGSGCPPGNDTIFAGGDGTLSLGVVVPGGQQVYVAKDGALSYTVPHSAYIPEGSVIDQFSKTAPGSNNLGSLSFETGFVACPTGGAGSTYQVYGQVDGFEASSECLGFSALTSASEQPGAWEY

>*Aspergillus aculeatinus*

MKLLALLSAAAGVSALPQAQQAQANYNSFSVMSARSGSPVHLLPLNAASGGFYLGLQNASSYCPDQVAELGACPPGTETVFDAGANSLDVSVPGGQQVYIAPSGALAFTQPHSAYLPAGSVVGAFSYSQTAGATFGYWTLPNSGLIACPVPAAASSSSAAPSATPAAGSAAATASKWQVFAAWSNATVPSGNVGDCLGFDALAVGRNGSAAAWEYI

>*Beauveria bassiana*

MHFLLFVLNAILAANLSSAVFVLSDEDIDDWLPGPLVQPSKECGQSYNPHDYVCHNGRLCPVINGEPLGLCGRTCFSKYMYICHNGTRLLTRPPLRTGFSTLRAENPREREEVAGQKNYTNGVDGKPMTACGHAWNIGGNTCSYCHTKTVPNCPSGNLTALSMARNSMLVMVPGGQRFFLTADWTIGYTSAHSAFEPPGSTKAGFRTYAGGGFFNNLNGAWGWAACGQGEQKRLHVRNSTNRQELKGCTGLNLRIEEFEGLRPAWQYE

>*Bipolaris maydis*

MKYTTALASLFAAAMAAPTAVQPVARQADASPYFTVISSRSGSPIHFQNMNARGGSFYLGGEGPSSYCPSESIGAENCPPGNTTTFAGGNDTLSLGVVVPGGQQVYVGPDGSLSYTTAHSAYIPPGSTVTGFSRTPAPNSDLFGYLNWDCGFLACPGSEEDGWKVYGNLDNVTFSDDCLSFSAIASPVDAPGAWQY

>*Boeremia exigua*

MKYTLAFAGLVAGALAGECYAPPPKVTSTGYATPPSPTPTPEPGPGYFGVISARSASPIHLQPLTARGGKFYLGGGPPSSYCPEVVGDACPPGNSTVFAGGDKTLSLGVVVPGGQQVYVAADGALSYTQAHSAAIPAGAIVDQFSKDAPSNGNSFGYLNFETGFVACPVGAADQGYQVYGQTQDFEASSECLGFSALTSSVDEPGAWQY

>*Botryosphaeria dothidea*

MKTFAAAATLLAAAPAVLGQYVGLAARSASPIHFGSINANSGKFWIGRETSSYCPSSVVPDCDTAYPGNVTVFAGGDNTLGLDVSVPGGQQVYIAADGSLSFTQAHSAYTGEGSVQTGFSKSESGSFGYLSWTNGFVACPSANGTSPYQIFGGGYSNNTACLGFDFLTSNYTGTSAWQYA

>*Botryotinia narcissicola*

MFSKAFIATLLASSAAATPIVSARAASDAFGLISIRSGTNLQNQAITASGGRLWIGKDTSSYCPDTVVSNCPEGTSTTFVSGGNALGLNTEVPGGQQVYIATDYSVSFTQPHSADTHGGVVSGWNYIAGENGSLGSLSFPDYGFIACPGEGAGVYGVFLTPGGSGTGNCTGIAVATVPYTGQGPSAWEYA

>*Botrytis aclada*

MFSKAFIATLLAGSAAATPIVSARAASNAFGLISIRSGTDLQNQAITASGGRLWIGKETSSYCPLETGCPAGTSTTFVAGESTLGLNTEVPGGQQVYVGTDYSVSYTQPHSADTHGGKASGFVYTAGENGGIGSLSFTDYGFIACRDTDGSYGVFLTPGGSGTGNCTGIAVGTVPYTGEGPSAWEYA

>*Paecilomyces variotii*

MKFLAVTALLPLLASALPAKRQIDIPPFTVTAARSGSPIHLLPLQAAGQSFWLGGSPATYCPAGEGIVCPPGTDTVIVGASALDVEVPGGQLIYVDPSGALSFTQAHSINIPTGSAVGPFQYEAGQPFGHWTFTGWNSTGFMACPQPQGNTTNWRVYAAISNATVPTGNIGDCLGFDALTYPWQSNSTAAAWQYI

>*Cadophora malorum*

MLSLLLVIISLFTLTTAQVFTLKTFNPGAPLNGQKIHASGQAFYLGLDGPNTYCPSKVDPNCPSPTDTIVAGALSAMWVEVPGGQQTFVRSEGSIGYTQAHSTSVPAGAHRGGFVNVIVDSDCSEDVNIFTWKAPDGSAEGILACPDVPSYMENIATFIIYVRTPEFNLTDCVELQGLEATIWLNPAPYGAWQYV

>*Capronia coronata*

MQIPSLIAVLLFVTAACASPRPQEVGSSGSPRSYSHGPTYAGNGTAAASSTAASTAISTSSSTSSGALTSPTAPLGVVAARSGSPIHFLQMNAAGLRFYLGGKTLSYCPDIVDDIGTCPPGNETAFSLCSMAVLVPGGQQLYVTPRGEIGYTQAHSTLMPNGSLPCPFTYYKAPGAPLGLLSTEAFGADGLMACPTDGDAWQVFANLKNATVPQGNISQCLGFDGLAVDYADLYAAWEYT

>*Chaetomium globosum*

MPRFTLAVAALVGLASGVLGDLKTCGPAQYDPAQYVCWNNQFLCPITAGEPLSNCAGACYSKFMYSCTNNVLALLPATETPFTLTVDNPTLPIHGKPVTAAGRHWTLNGETSSYCPEQVGDACPPGDKTVIVSSGGRVSMSVMVPGGQQAYLDPFWNMGYTQAHSAAIPAGSLLEGFGAYQGGGFVNLNGNGWGWAACPPRASGGGGPGWNLVARNETNAANYSECHPVNLKINSLPSATFGAWQYT

>*Cladophialophora carrioni*

MLHSLLLLAVSLFLAVACASPRHQADYDDPLAIRDIVTVTSTLTATVIASTLTSTTTSSTTTYATPLTQPTSWLTVTAARSGSEIHLLPMNAAGLRFWLGGNTISYCPEPAEAQGACPPGNTTVLSLCSMGVLVPGGQQIYVTPRGELGYTQAHSVSMPPGAVSCPFTYTKAPGAYIGRLLMSIGAPFGITGFMACPTRSRGIYQVFGNLKNATVPLGNVSQCIGFDPLAADAPGLGAWQYS

>*Clathrospora elynae*

MKYTLTFTSLLAAAQAASYCAVQPPPASSSTVPVPSASATPVGPAYFGVISARSASPIHLLELTARGQKFYLGGPGPSSYCPVAQVGSACPPGNSTVLAGGDKTLALGVIVPGGQQVYVAPDGSLSYTGAHSAYIPPGSIVDKFSIEAPTDGEAFGHLSFDTGFIACPAGVGQGYQVFGQITTTKFPDDCLGFDALTIASNEPGAWQY

>*Claviceps aff. humidiphila*

MKSSLSFTLTTLVPLLAAGEPIASSAVSLAVNLPGSPLNHQPVVANGGKFWIGKPTSQYCPDNVEKLGGCPKGKDTSIWVNNEQGTGSLNVIVPGGQQIYVGVDTSLSFTRPHSAYIPPGSTSAQFSLIPMNQLAMVGRRMLACLPPNTDINGSISAWQVFAPRTDQAAPPDRQCVAFVAQTAAAKDAVWEYI

>*Clohesyomyces aquaticus*

MKNAILAFSALVASAAAQDYFTAVSARSASPVHLLPLNANGGKFILGGEPSSYCPEQVKDACPAGTSTVLAGGYGTLSLGVVVPGGQQVYVAPDGSLSYTQPHSAYIPEGSVRVGWSREAQTDNVIGFLNFVSGLIACPVAGKPWQVYGKLPGLTFDSACLGFNALTINGTEAGAWEY

>*Coccidioides immitis*

MKLTILAVILPLLAAAAPGLEARQSEPPYPPKIKVMASRSGSPIHSLPMNAAGRAFWLGGTTQSYCPIQVGDNCPPGNETVIAGLSSMYVLVPGGQQMYVEPSGAVGFTQAHSTYIPPGSYIGGFTYEPRGEHGIYSFTGWGADGFMGCPDPEVGFHQVYANIQNASVPTGDIKDCLPFVALAITYPGNNPAAWQYV

>*Coleophoma cylindrospora*

MFCKTLLASALAGLAVAAPAPIAQRQEASSFSAIAVHSTSAVHLQSINANGGKFWVGAEKPTSTYCPLSTGCPAGNITTFAAGASTGTLSLSVEVPGGQQVYVDPSGALSFTVPHSAFMPAGSQTGPFVYTAAASNQTVGTLTFGNGTQGFFACPDDAVSAWQVYVFTAGSNSTSSCIGIGLSTVNEPAIAAWEYA

>*Colletotrichum aenigma*

MGLKPAILSLILSLSCIITLAAAESYTLTAYAPAIRAIHGRPINASGRRFIIGLGAPMTYCPDVVPRGSCPPGNITTVDGLLTGMMVMVPGGQNIYVDSEGFVGFTQAHSASWPANSTLMDGFYRTTAKSSSRCEAEPPTDLLDFCDPKNAASEGVYACPYPDLVGTWVLVAGSARSRARRGDCERLEGLELHESEIELGAWQYT

>*Coniochaeta pulveracea*

MSRTTFLATALLSLSSIALAELKGCGASQYDPAQYVCYDNSFLC PIIAGEPLSFCAGACYSKFMYQCVSNTLAELPQVAQGTPIGLTASNPSLAIDGKPITACGQHWSIGGQTCSYCPEAVVGSSCPAGKDTVIAASDGGASMDVMVPGGQLVYLGPDWNVHYTQAHSAYIPSGSIVTGLKAYHGGGFVNLNGNGYGWVACPPTAAGGGGENGWNLVAKNSSNAGNLNNCYAVNLKVNEL

>*Coniosporium apollinis*

MARLSFFLLSALSLGFAAAEDLQDCNGARYLPSRYTCYDGKYLCPVYNGVPTLRCGDACYSAGQYSCNNGVLSGLSFTDTSYTLRADNPTAPFDNAPIEACSRAFWIGRGTCAYCPSVVGEYCATFPNDTRLIGSSGMSTSVPGGQMFYIDPTGALGYTGAHSAYYPPGSVFGVTAYKNGDQGEFKYTPGEGWYACPTTADANVLQIFQKLPGITLASNCTGLNILVTEVGGTGAWQYT

>*Corynespora cassiicola*

MKTVIAFSALVASTVAQAGYFGVMSARSASPVHLLPVQARGGAFFLGGEASSYCPPSLPEGSCPSGNSTVFAGGQGTLSLGVVVPGGQQVYVGPDGALGYTPPHSTSKPEGSIVEGWNKTEAGTFGHLSIGDGLVACPAGEGKGYQVYGQIEGVELGSDCLGFSALTVNATGPGAWQY

>*Cucurbitaria berberidis*

MKSTFAFTALVTTALAGPYCAAPPATPPAASSTPTPTPDPSGPSYFGVISARSASPIHLQALTARGGKFYLGGPGPSSYCPVAQVGDACPAGNTTVLAGGDKTLALGVIVPGGQQVYVAPDGSLSYTGAHSAYIPTGSIVDQFSRQAPAAGQQFGYLNFETGFVACPAGEGEGYKVFGQVQGGTYGPDCLGFNALAVATDKPGAWQY

>*Cudoniella acicularis*

MYTKTLIASILASLAAAAPAPVARDNSVFAMIVIHSGSPIQGAAISENGLKFWISKPTETYCPESVEGKETALSVNNDSGLAGMHTTVPGGQLAYVTEDGQFSVTQAHSAAIPAGASQTGFKYNPQPEAGMVGELLFNGQGLSACPTGEPTDNSIAIYQIYANGASGFTQTNCTGITIGTAEVDAPPV

>*Dactylella cylindrospora*

MVFIKVSALLSLLAAVSAAPADPPNVIRNNSYGFLALRSASPIHFGNLNASNGKIWIGKDTSSYCPLPPRSCPPGKVTAFWINDTVSMAVSVPGGQQLYIDPKGALSYTQAHSGYMPKGSITNGFKVEDNHETHLSYFSHARGGFLACPTGKKATGPYQIFVDVDGKLKDKDVPSKKKRDCLGFTAAGGSAQEQFAAWQYA

>*Dactylellina haptotyla*

MKFTLATVITAASVVSAAPTYSSEPKPLALTMMAIRSASPIHFGRVDASGQKFYIGLDRPSSYCPSPPVPVGSCPSGNYTSISWGQYGPYMNVEVPGGQQIYIAPDNSLSYTQAHSGSMPPGSTTDGFTLGPKQENTLRHIDHKSGGFLACPAKKDVGPWKVFVGSVTDKNAPSGSAADCLGFLNTGIEFIGEQFGAWQY

>*Delitschia confertaspora*

MKTVIALSAFVAGILAQAPSSFGVMSTHSASPVHLLPLTARGGKFFLGGPGPTSYCPEQVQPNCPNGTETLLVGGDQTLSLSVMVPGGQQVYVAPDGSLSYTAPHSAYVPEGSIRDGWYKTEGDMFGNLKFEKGLIACPAAGQGNGYQVFGQLAGLTFAPECLGFNALTVNQTGPGAWEY

>*Delphinella strobiligena*

MKTFAVASVLTLCSTAFAAPLQARHDTTKPFTLEVYFSNLPPVSGPLQASGGSFWIGKNTSSYCPDNVGDCPSGNSTSFLSGSNGALSLNTEVPGGQQVFIAPDGLLSYTSPHSSYIPTGSITTGLVVDADNNLLNKYYNYWICSTDSAATMWQVWLEYRDDQGSIKNNGLTDDNSCARINLVANDTVSGAAAWQYS

>*Dendryphion nanum*

MKSIIAISAIVATTLAQAPPYFGVMSARSASPIHLLPLTARGGKFFLGGPGPSAYCPPQVGDACPPGNSTVLAGGYGGLSLGVIVPGGQQVYVAPDGTLGYTIAHSAYKPTGSVVDGWSRTEGDNFGNLAFTGGLYACPASEGNGYQVFGGIGGVVFDPACLGFNALTYNTTQAGAWQY

>*Didymella keratinophila*

MKYALAFAGLVAGALAGGECYAPPPATSTYGAPAPTGTPAPDSGYFGVIAARSASPIHLTTLTARNGKFYLGAGAPTTYCPDNLPAGSCPPGNSTVFAGGEGTLSLGVVVPGGQQVYVAPDGALSYTIAHSAYIPTGSIVDEFSKTAPSNGNSFGYLNFETGFVACPVGAADAGYQVYGQTDSFEASSDCLGFSALTSSVDEPGAWQYSS

>*Diplodia corticola*

MKSFAVAATLLAAAPAALGQFIGLAARSASPIHFGSINAAGGKFWIGKETSAYCPSSVVPDCETDYPGNATVFAGGDNTLGLYTAVPGGQQVYVAPDGSLSFTQAHSAYIPTGSVTTGFTLSPGTIGYLGWTDGFIACPSLNGTFPYQIYGGSTNQTTCLGFSFLASNTTGGVNAWQYA

>*Drechslerella brochopaga*

MIVAKAAALVFLATAVSAAPATTPQPVVKPWNFLALRSASPIHFGSLEASGLGIWIGKPTASYCPVDPKYCPPVSNTTSILVTENAYGAASMDVIVPGGQQVYIAPSGALSYTQAHSAAIPAGSFRTGFIVSLPGSNGISIFSHCEGGFVACPVVANKGPWKVFVAFKGVLPKNTDVPSGKAADCLGFSAATAGPAQYHPGAWQYT

>*Drepanopeziza brunnea*

MVSLSNLVFGLSILATSAVAQFTLTVVHPDSPLDGLKVHASAQAFYLGLSGPSTYCPSVVEPYCPDKTDTVFAGMGGMWVNVPGGQQTYVRADGVFGYTQAHSAYVPEGAYRGGFVNVILSGDCIEDVAVYTWQDPDGSEDDAGILACPDVPYYMNGTATYQIYAKTPEFNLTNCVELGGLKPTAWPQNAGFGAWQYT

>*Elaphomyces granulatus*

MKFIALFAGLPLLAAAAPTQRALGMPLQGDLCMPFQHDLGTPLQGGDFTAFAWFEHEPIHPPIQARGLRFWVGGEPATYCPSPPVVHCPPGKVTAFSGGGYGLSVMVPGGQRVYVSPTGELGYTQAHSAYIPPGSFLGPFEYTPGIPFGHYTFNGKGFIACPNSKEGPTTWKVFAHIPNAIVPTSDIQDCLPFQAVTEIFGSPPAAWQYT

>*Elsinoe ampelina*

MHLSTITSILILAALGTATPLDMEKRQNQQPPFTLIASRPNSPIDKLKIQAAGQRFFLGPQSPSSYCPVQVGSSCPNTTDTVLLNGAMYAMVPGGQATFVQKSGALAYTQAHSSFRPDYYSSDLAAYSDGSYRGPSGAGFAACPVQGQPGQWQVFAETPGFTGQNCLCFQAKWQKWTGGIGAWQYT

>*Endocarpon pusillum*

MRLLTYILLAAPILASCVPHDPRGHDNSRRSDKHDDKQDDKSKDRPRPTSCTTTTPPASTTSCVPTTSPTKAFTVIAAHSGSPIHLLPMVAGGLGFQVGGGTSSYCPTVVGASCPPGNETVFVGGSISVMVPGGQQQYVEVGGRIGYTQAHSASMPVGAIPGGFSYYKCPDEQFGHITTSIFNATGLMACPDNSFGTQRYVVYANIPNAVVPSGNVQDCIGFDGLTSDYNGPLPAAWQYT

>*Epichloe festucae*

MKLTLAAVIPLLPLSFVEASPTYHAANTSAVAITAALPGSPVHNRPLEASGNKFWIGKPTSTYCPLNAEECPKGKDTTIWVGNELTTCGMNVVVPGGQQVYVAPDGSLSYTQAHSAFVPPGSTRDGFCLLPERPDGLRELAISSRQFFACPEAKDSGVWQIFARADDQDVRGQSAAQGQSTAHCLKFTALASPAKDPVWQYA

>*Epicoccum nigrum*

MKYALAFAGLVAGALAGGECYAPPPPAPSSYATPTPTPTPGSNYFGVISARSASPIHLLPLTARNGKFYLGAGAPTSYCPPGLPAGACPSGDNTVFSGGDGTLSLGVVVPGGQQIYVAANGELSYTVPHSAAIPAGATVDQFSKTAPGANGLGTLSFETGFVACPVGGDGQGYQVYGQNQGFEASSDCLGFSALTCEF

>*Exophiala aquamarina*

MYPINLLFGFLTQILLVSAIIHVRQDVNTTSEANYGHHTTYTTVYNSTITVSVTHTTLTTEPRSSATSTSSTSSAPLSRPSLPFTVLAIRPGTPIHQLRVNADGQRFYLGGQPSTYCPKRVEELGGCPPGETTAFGLCSMAVLVPGGQEIFALPNGELGFTQAHSALLPPGSSTCPFEYVMEHRGDQYAHLFPNATNAFRAKGFRACPTNDGRWQVLLALANVTVPSVTNQTECLPFAAFAVDYENGYAAWQYN

>*Fonsecaea multimorphosa*

MLHVLSLVAISLFSALSFASPHPQTSTTPCISETDGNAGASDTVITVASTLTATVVASGSATSSSSTTTAGPLTTPTSWLTVTAYRLGSPIHLLPMNAAGYHFYLGGDTVSYCPTEVEEEGGICPPGNQTVLSLCSMGTLVPGSQYLYVTPSGELGYTQAHSVSMPEGSVQCPFTYSKAPGATIGRLDLRVFGALGLVACPTYGGTWQVFANLKNITAPRGNVSQCLGFDPLAFDTPNIGAWQYT

>*Glarea lozoyensis*

MYTKNILVAAFATLAAAAPTARSTADVLGLIATHSGDINVHLRSINENGLKFWIGKPTTTYCPSEVVTNCPNGTFTQLIASTDSEGISMNVEVPGGQQAYVANDGSLSVTQAHSGSTGTNGLRGPFSYTPAASEGTVGQLSFNGNGFLACPSAEASVYQIFAAGYPGFVAPEGCIGVGLATSVIDSAPVWQYI

>*Glonium stellatum*

MAVLPSLLLFALLLASARADTCGSQQYDPSQYVCYYNQFLCPFVNGEGLSYCNGACYSQYMYQCTDNNTLALLPAVNCPFTMTISNPQAPSINGLVVNACGGGFSAGELSETCTYCPTQVAPYCPSGNETVLYSSGSMASEVPGGQQWYVDPSGALAFTAPHSAFIPSGSQIGGFAAYQNGGLVNLGSPFGFYACPHVSQSVVVYWDIFQQIAGVTVSSSCVGVNILVHPVEDAEYAWEYT

>*Helicocarpus griseus*

MKSAFLASLLPLLAAAAPAPAEVPRFTVMAARSASPIHYSPMQATGRHFMLGGEPSTYCPQPPVEECPPGKITVVAGAGPTSSLNVMVPGGQQIYVEPTGALGFTQAHSAQIPEGSLTTGFKFTPGDQFGHWTFEGFGADGFMGCPPQNPDESPLYQVFANIKNATVPTGNVDDCLPFSALAPPYTGPNPAAWQYI

>*Heterodermia speciosa*

MIPLLCLGAALLYLAASSTFIHHREAPYISFGLIAIHAGSPIHLSSVHAAGNLFWIGKDTVSYCPAIDGLECPGGNHTTFITVSGSQYLRLNTMVPGGQSVYVLPDGTLGSTPAHCDFSPEGSRFEGFNAIAPDSPYNPGSLTFDGADFLACPTNGTSGPYQVFVNTDGISDSDVPCSLWECISFVGATFRNTNDSASAWQYQ

>*Hyaloscypha bicolor*

MPPFAPFLSLLIAFIGTTSADIFMANASLPDSWVDGQPINAAGQAFWVGGSPSTYCPSIVGANCPDATAQTIIIAGFSALDVEVPGGQQIYVNVNGALGFTQAHSAVVPTGAYIGGFFNLTYMSDCSTPRTVINWKSPDGSSEGILACPTKPSGPNVTVSYQLYAKTPAFNLTNCDGGVPVDGLTATYLSSPAPYGAWQYV

>*Hymenoscyphus varicosporioides*

MKSVIAVSSLIAVASALTAPTGPFTSGAWNPTTGWVGSAIQANGGAFFIGKDPSSYCPEVEGLDCTLFPGSSTVFIGGNNTLSLDVAVPGGQQVFVTPEGAIGYTPAHSTFMPTGSVLDGWDRFQSQAGGAPVPMNFQSNSFLACPVNATESGVYQVFARSLKTDATDCTLFQWRTYTASGVNAWQY

>*Hyphodiscus hymeniophilus*

MALLKSLFAIVFLWFTFAAAQSVAGQFTLTIYQPSSALHGQIVNAAGEAFHLGGSPASYCPLTNLTLCPAGNQTIFAGMGAMFVEVPGGQEVYTTSAGAISYTQAHSASIPPNAYIGDFTNVTVMDNYSAPVYLVNWQAPQTVNSTKGILACPQVAPGNSTTVVYQVYANTPAFNQTNCTLVDGLLPHYTPPGVIGAWE

>*Lasiodiplodia theobromae*

MNHAICRDDIICPVVDGFPQDLCGGQCYSSLNHTCDNGTICPRAPQEGPYLIRVLTWYGDMDELAIDACGGAFFAGRSSSSCTWCPYQQFPETCSAAVNITVLEGTENMHVIVPGGQRFYINALGALAYTRAHSGIIPENSIVGEIQAFQEGGFIYAAGGGWYACPASEPGVYQIFNRLEGVVLDAACIGINIAVNSLEGGLFAWQYQ

>*Lentithecium fluviatile*

MKTAITFTSLIASTLAEGYFGVMATRSASPIHFLPLNANGGKFYLGGTPSSFCPPQVGDVCKAYPGNSTVLAGGDGTLGLAVIVPGGQRAYIAPDGSMSYTVPHSAYIPEGSIVDGWTKEEGEAYGWLRYKDGLIACPPAEESKPWPVFGQRSNVTLSPDCLGFSALAFNSTKAGAWEY

>*Lindgomyces ingoldianus*

MKNAFAFSALVASAFAQDYFGVMSARSGSNIHLLPLNAIGGKFFLGGSPSSYCPPEVGTACPPGTSTVLAGGVGTLSLGVIVPGGQQVYIAPDGTMSYTAPHSAYKPEGSVVDGWSRSPVNDQFGYLTFSGGLVACPVDANNKPWQVYGQIASVTLSPQCLGFSALTINGTEAGAWEY

>*Lineolata rhizophorae*

MKPSTILVSLLSALPAAILAAPTSCGSDDSDSGSGVTRFGVMSARSASPIHLLPLNARNGHFYLGGETQSYCPEEQVQNCPAGNETVLAGGDNTLALAVDVPGGQLVYIGPDGAMSFTPAHSASMPEGSVLTGWNYTAGDPFGHLQWFPGGLVACMDRDNQVFGQVEGFNSTGCLGFDALAIATEEVGAWQYT

>*Lophiotrema nucula*

MKITLVGFLLVASAVADQCGSHEYNPETHVCNDNQYVCPILSGEALASCGESCYSYFEYSCDNGDLVPLPAVDKPFELVTYNLYAPDVHDFPVYACGHRLYFGNGSATCTYCPNMPQLHGLCNDPGLITNRTILLGGSGGYGLATQVPGGQRGFIGPEGGLNYTQAHSASVPNGSQYDGILAYQQGAFIDTKSEFGWAACGQFNADDEVPVWDILAFRKAVVLWDYYDVCAEVKLLVVEADLENGTYVPWQYE

>*Lophium mytilinum*

MAPIFSFPAVILALLITSSTASVCGTTDYDPSKYVCWYSSFLCPYTNGSPLSYCNGACYSPHMYHCTDGVLALNTIITSPVSLTISNPTLPDLDGYPVRACSQKFSTGYYSQTCTYCPKDVVRSCPNSTTTVIFPSNSLDVLVPGGQRWYVDPSGALGFTQAHSAYIPPGSQVDGFQAYEGGMLVNLRSPWGWVACLAGREIPGVGSGYEVFQQLGPEVKEGCFGVDILVGPPEAGFAVWEYT

>*Macrophomina phaseolina*

MKTFAAAATILAAAPAVLGQFAGLAARSASPIHFGSINANSGKFWIGKPTSSYCPSSVVEDCETAYPGNVTVFAGGDNTLGLDVSVPGGQQVYIAADGSLSFTQAHSAYTGEGSVQTGFSKSESGSFGVLSWTNGFIACPSANGTAPYQIFGGGYTNNTACLGFDFLTTNYTGAGAWQYA

>*Madurella mycetomatis*

MARLTLAAALLSLGHGALAALETCGDAQFDPAQYVCWDNQFLCPVTAGEPLSHCAGACYSKFMYKCENNVLTLLPPVETPFTLTVSNPSVPIIHDKPITAGGLHWNIDGETSSYCPDQVGDACPPGDVTAFVARGGSVSMNVMVPGGQQAYLDPYWNMAYTQAHSAFIPPGSTVGGFGAYQGGGFVNLNGNGWGWVACPPRASGGGGTQWNLVAKNETNAANLDSCRAVNLKINPLPSGSISAWQYT

>*Magnaporthe oryzae*

MKSLFLTLGLLGLSASALAEKCGDANYDPKSYICHNGNFLCPVVNGEGLSYCNGACYSKFMYQCNSGTLSQLPLLEQGSKFTLTAWNPASPSVHGKAIANSGRHWWVGGETSSYCPDIVKDQGACPPGTVTAMAYSGGMNTLVPGGQGYYLNANWNVEVTQAHSSYIPSGSTVGGLVAYKNGGFINTNGQPTGWVACPGGSDGRWNLVAGNASSADALKSCTGINLQVNYLGDSSVPASAWQYI

>*Magnaporthe poae*

MARLTKRLTALALTLLLPAATQAAQCGSAIYDPANYVCHDNQFLCPVVNGEGLSYCNGACYSRFMYQCVNSGTVLAQLPRLDDGARFALRAWNPTSTAVHNKPVAACGRTWTVGGPTCAYCPVEVVGAACPAGNATAMAYPGAMNAMVPGGQAYYLDANWGVGYTQAHSASVPLGATLGGLVAYQGGGFVNLNGPGTNWVACPPTAGGDGTGWRLRSENATTAGDRGNCVGINLAVEIVANQASAWQYT

*>Melanomma pulvis-pyrius*

MNSTIIFVMMFAALVSSETCGLTEYNSDDHVCYFGYYLCPILNGEPYQWCGGARGGCYSKFKYSCKANKLYALEKTGSAFELTLSNPKIPGIDGLPVHASGLKLVSGQDAKTSTYCDPSAPEGLCASAIKNRTIFGISGGIWFLQVQVPGGQRGYLDNNTWGDGLQYTAAHSASIPEGTEIDDFVAYENGGFFSLLHPYGWEVCQAKPHSGEGLVYSLRSRPDDGPVPDRCAGVNLLVKDSGLVDADHLWAWAYI

>*Metarhizium acridum*

MIPTPFIVISLLCLFAEAGLISRGESTVDSAVVVFAILRDTPIHHGLFAANGSAIWIGKDTASYCPPVVEERGECPKGKDTSFWVNDRCGMNAIVPGGQQAYVAPEGTISYTAPHSAYIPPGSITTGFHLVPTQFEGFWEFAIDSHELMACPETPGQGPWQVIVVRENSSVPSRDPTDCLKFKAYAKAVKDPVWEYLSTLSSQPIVVQEYL

>*Microsporum canis*

MKISFITAFLLPLLTLAAPAPADPPANIQVTARHPGRKHVDKLPMQAAGRAFWLGGSPATYCPAVVGDACPPGNATVILGLNSMSVLVPGGQQMYVEPSGKLGFTQAHSISIPPGSYIGGFAYKPLNRKSGSFYFGGWGATALMACPVPDSKFYQVFANIKNAMVPGGDVKECVEFVGVATKYEGATPAAWQYT

>*Microthyrium microscopicum*

MLRLYAFIASASGVLAAPSPAGFELHAYYTNYPGLGAPAINASGGSFWIHKPTQSYCPDSVANITTCPLGNGTSFVAGGQAGTLFMNTEVPGGQQVYVAPNGLLKYTVPHSGNIPEGSLTTGLIVNATTAQLNRYYTFWMCNLPEDSNVWQVWLQYEDEKGSIRIGGQGDKNACTRFVLISEPVQGIGAWEYL

>*Monascus purpureus*

MKLFSLLTLGALLSAANALPTASRRFIPPSLRHVGSNSTNATLSSYNATSFTFIADSNFTEIQFQPLVASGQKFYVGGNSSTYCPDVVRPCPKGVDTVVYPKGSLNVAVPGGQQIYVAPNGALSFTVPHSANIPTNSSTGPFVFKNTTTAAPFSVGEWSYARSGFVACPVNATGVERYQVFVANSTGSSADCLPFKALAQPWVATANWTVGAWEYF

>*Mytilinidion resinicola*

MARIFSFPIIILALLITSSTASVCGTTDYDPSKYVCWYSSFLCPYTNGSPLSYCNGACYSPFMYHCTNGTLYLNTITTLPVSLTISNPTIPELNGYPVRACSQKFNTGYYGQTCTYCPKDVVPKCPNSTTTVVYPEGGLDVVVPGGQRWYVDPSGALGFTQAHSAYIPPGSQVDGFQAYESGALVNLKSPWGWVAC

>*Nannizzia gypsea*

MKASLITAFFLPLLALGAPPPPADPPANIQVTARHPGCKEVDKLPMQAAGQSFWLGGSPATYCPEIVGDACPPGNATVILGLNSMSVLVPGGQQMYVEPSGKLGFTQAHSAYIPPGSYVGGFAYKPLNKKFGSFYFGGWGATALMACPVPNNKYYQVFANIKDAKVPGGDVKQCVEFVGVAKAYEGATPAAWQYT

>*Neofusicoccum parvum*

MKLAILATLLAAASTALASPTTPNEPRQATSPDGFKLNIRWSNGPLRGAISANGGSFWINKATKSYCPGRDVIHAACPAGTDTSFVGGGPDGRLSLNTVVPGGQQVYVGPRGLLRYTPAHSGDMPKGSITSGLVVQETSGFLNPFYQLWMCNTDADAGVWQIWVEQRDAEGTVKNWGKTEKNACTRVNLEMERVDGVGAWQYD

>*Neurospora crassa*

MVVEPKSSPESHSSILAMMKLTLLSAALLAFGDSAVNALSSCGQSQYDPAQYVCWSDQFLCPITAGEPLSFCNGACYSKFMYKCENNVLSLLPAHTGTPFTLSVSNPTLPIDGKPVTASGQHLSLAGNTSTYCPVEVVGAACPPGNITAFFAGNGGLSMDTMVPGGQLAYLGPDWNMHYTQAHSAYIPAGSLTQGFGAYEGGGFVNLNGNGWGWVACPPQASGGGGTAWNLVGRNETNAASLTACTPVNLKINPLPSGTVGAWQYT

>*Ophiobolus disseminans*

MKYTITALALVGAVFAGACPKPSPSPTPSPTPEGPSYFGVISARSASPIHLRALTARGGKFYLGGPGPSSYCPVAQVGAACPPGNETVLAGGYKTLSLGVIVPGGQQVYVAPDGALSYTGAHSAYIPPGSIVDGFSREAPAAGAQFGVLNFDTGFVACPAAAADQGYQVFGQVDGATFGDGCLGFNALAVAGDKPGAWQY

>*Orbilia oligospora*

MKFTLATLLATAVAAAPAPSNLSFTMMSLRSASPVHFGSVNASGQKFYIGLEKPSSYCPTPPVPASSCPSGKYTSVTFGSGYSSLNVAVPGGQRIYIAPDGSLSYTQAHSAYIPPGSTQDGFILGPKNKDNGLRPITHKKGGFVACPAKKNVGPWKVYVGQPASKDVPSGCASDCLGFSNQAVEFTSEQFGAWQY

>*Paraphaeosphaeria minitans*

MKSVAAFSTLLACAAAQTAYFGVISSRSASPIHLLPLQANGGKFFLGGTASGYCPSAAIGNETCDHYPGNSTTFAGGNGSLGLGVVVPGGQQVYVAPNGALSYTQAHSAYIPEGSERTGWNKTDRNDGSNLGSLSFDGGLVACPRNETQPWQVYGQIANFTAPEGCLGFGAVTFNVSDAAAWQY

>*Paraphoma chrysanthemicola*

MRYTIALTTLVASVFAMESCAAPAPAPTPSPAPGPGYFGVMSARSASPVHLRPLVARGGKFYLGGETASYCPVAQVGSACPPGNTTVLAGGETTLSLGVVVPGGQQVYVAPDGALSYTQAHSAYIPTGSVVDGFSREYPAAGQAFGYLQFDTGFVACPAEGDGVYQVFGQVDGATFGPECLGFNALAVSVDGPGAWQYS

>*Parastagonospora nodorum*

MKYATVIAALTSTALAQSSCSSAPPASTPTPPAGPNYFTVISARSASPVHFQTLTARGLKFYLAGGPPSSFCPTELPAGSCPPGNTTVLAGGYGTLGMGTVVPGGQEVYVAPDGALSYTQAHSAAVPAGSYRAMFSREPSGDNSFGYLNFETGFVACPAKAPETGYQVFGQVDGATFGPECLGFSALTSETDQPGAWQY

>*Patellaria atrata*

MKSTIIASSLFSLLPAIMALPAPALAQEAEVSNNFVGISARSASPIHLLPVVTHNSAFWLGGETYSYCPTQVEPNCPPGNTTVFAGGDEYLGLGVLVPGGQSVYIAQDGSMTHTTAHSAYYPPGAVRDGWTRTEGENFGHLSHEAGLVACPVEGGYKIFAQIPTFVAPNEDCLGFSFLTGNVTEQVGAWQY

>*Peltaster fructicola*

MKTAAAILALAGAVTAQNYMRVMSIRSGSPIMYAPMVARGEKIYLGGETASYCPETVGENCPAGNTTTFVYGQGHLSMGAIVPGGQAVYVDSQTGAVGYTIAHSAAIPQGAYQAGWNVTQNVAAGGILGRLDFEGGLIACPVDGTYPYQVFGQVEGVSFAADCLALMRCSATALRHQHGSIPKSSDFDEKHVQLA

>*Penicillium antarcticum*

MKFTLTALALPLLASAAPAPAANPPFTVMAARSASPIHYLQLNAAGQKFYLGGNTASYCPSQVKDCPAGNQTVFAPGGTSMDTMVPGGQQVYIDPNGALSFTQAHSANIPAGSSYGPFAYEAGEPWSHYVYQGWGASGFMACPTEDSRWQVFGALANATVPKGDVDECLGFSAMAITYEGDVPAWQYI

>*Periconia macrospinosa*

MKTFVALGAFVANAVAQTTGYFGVVSARSASPIHLRSLEANGGHFFLGGANSAYCPPELGEEVCAAYPGNQTVLAGGEVTLSLGVVVPGGQQVYIAPDGALSYTQPHSVFKPNGSVVDGWSRTEGETFGNLNFVDGLVACPTAGDKPWQVYGQLPGVELSPDCLGFSALTVNSTGAIAWEY

>*Phaeomoniella chlamydospora*

MYIASAVFAVLPLLAAAIPTPSSTKSSNTAYTTATASATPFTVMALRSASPIHFLQMNAANTSFWLGGATRSYCPIDASQCPPGNVTVFDGPNALSIEVPGGQQIYVAPSGALGFTQAHTANMPAGSVISPITYTPQTGTAYNGVISTSAFGATGFMACPTTTNNWQVYAAISNATVPTGNLADCLGFDAVTVDWTGANPEAWQYV

>*Phaeosphaeria sp.*

MKYTIALALASTAMAASECSSAPPASGTNAPLPGAPSYFGVISARSASPIHLQSLTARGTKFYLGGGPPSSYCPVEQVGDACPPGNYTVLAGGDKTLSMGVIVPGGQQVYVAPDGALSYTVAHSVAIPDGSIRDEFIKEEPSNGNSFGYLNFPTGWVACPAAEGSGYQVFAQVETATFGPDCLGFSMLTSAVDEAGAWQY

>*Phialocephala subalpina*

MFSLRNTLTVALTFGTAVADTFTLTASLPGSPLDGLAVNAGGEAFHLGGSPGFYCPTSVGSACPNTTETIFAAELTALDVEVPGGQQVYVQANGALGFTQAHSASVPVGATIGGFVNVTVASDCHAPWEVITWKSPDGSTGGIEACSEVPVGTNGTAVYNVYAVTPAYNQTGCVKLAGLLASYLGTGAGFGAWQYI

>*Piedraia hortae*

MQFNTFTLLILTALSAAAPVKQCEKYSYNPDNYRCYPGSKPVLCPVIAGVATKPCGSACYSPEQYSCSNNQLVQLPPLNDAFTLVAHHPINSPSNLDGKTIEASGQHFYINRPAGVYCPSVAGGICAASSNRTILFPGALDVVVPGGQEIYVQKNGALAFTQAHSASTTDLAVLGLGGPVYKGGAALGPNGVAWKACPVDGGAWQVFVPLPGVSFSAGCVDFYAHAATADGLGVAWQYD

>*Plenodomus tracheiphilus*

MKYTVALTSFIIGAMAGACPAPVPTSSSTPAAPAPTGYFGVISARSASPIHLQSLTARGGKFYLGGPGPTSYCPVAQVGSACPPGNETVLAGGDKTLGMGVVVPGGQQVYVAPGGALSYTIAHSAAIPSGSIVDQFSREAPTGDSQFGHLNFETGFVACPAGEGQGYQVFGQVTDGPAFGADCLGFSALTVATDQAGAWQY

>*Pleomassaria siparia*

MFCVALLILSSFALVRGESCESTTYDPEKYICYEPNGNLCPIINGSPYSYCDQGAYACFDTHQYSCSSTQLEPLEPQTESFELIVSNPNIKELHGLPVHAANYQLRTGPNARTSTYCPEGTIFACDKRVTNRTVLRAGGGRFGPVSVLPGGQHGFLYPSSLYPSYTRPHSGYIPKGAQVDGIYAYKGGAFVNSYQPEGWWACPPGSNVDEADGVWNIYGAEQNVTGCVGVDLLVKQVEPELEWFAWEYI

>*Podospora anserina*

MTRFSLTAAVLIGLGHRVLGALEQCGPAQYDPTNYVCWENQFLCPVTAGEGLSYCNGACYSKFMYTCNNNILSLLPPAESAFTLTVSNPALPQLDGKPVTAQGLRLWLGGETKSYCPSVVDPNCPPGNVTSIVAGGFGGAGMNTMVPGGQQVYLTPDWNVGYTQAHSAYMPFGSTSTGFAAYQGGGFINLNGNGWGWVACPPRASGPAGPEWTLYGRNSTNAESLNYCTPINLKVTPYPGQGAAAWQYT

>*Polychaeton citri*

MAPITTYMMALWATSIAAAAPNSGLKKCGSAYYDPSQYTCYGSKVGQLCPIFNGIPDKLCGNACYSEYMYTCDNGVLRTLPPANTTFTLTAISPKYPEIHNMPIQAAGLHFYVSKTPGTYCPTVAGDICTTFPNNETYIYPGGSALAVMVPGGQQAYTQTDGAWAFTQAHSISLYNVSSYFLGPAYQGGGMFGPNNTNLIACPQSNGAGKTSWQVFADNPAVYRTATCLGFYARVNKKESGSFGAWQYT

>*Polyplosphaeria fusca*

MKTAAAFSALFASAMAQGYFGVMSTRSGSPVHLLPLTARGGKFFLGGGPPSSYCPPEVASACPNGTQTVLAGGDVSVSLGVVVPGGQQIYIAPDATMSYTTPHSAYVPEGSVRDGWTKTEGENFGNLKFEGGLSACPATKQGDGYQVFGQVEGVTLAPECLGFNALTVNATGPGAWEY

>*Polytolypa hystricis*

MKPAIISSLAALLAATVTAQDRPSSFTVMALRSASPIHFLPMQAAGRHFWLGGKPSTYCPVQVGDNCPPGNHTVISGLTSLSVLVPGGQRMYVEPSGALGFTQAHSIYIPPGSIQYGFEYTPGTEFGIYGFTGFGATGWMACPVPDRPSYQVFAAMRNATVPSGDVDDCLGFSAAAIEYKGPSPAAWQYA

>*Pseudogymnoascus destructans*

MRFSTSAAVVGAVLSSQVSAQDPISAFTLKAWNPSSTLQGEDVNAAGFGFYLGLEGPATYCPTIPELSCPAGNETVVYKGGMSLSVVVPGGQQTFVEASGAISYTQAHSASVPAGAYIGGFTSYTTLDGNGVNQTIVSWETPEHPTFAGLVACPKVSEYVDATATHRIYGRTPGFNQTDCVELKGLIAVAQPNNDPGAWQYT

>*Punctularia strigosozonata*

MKLLAFVAFAVSPLFSAAQAPDPFTITALHSGEVNIHLQPINANSQAFWIGRPTSTFCPNPASECPPGTQTAFIGGSSTLELDDEVPGGQQIYVQSNGALGYTAPHSAVIPAGATLTGFFWTGGAAAGDLGFIGLGATSFIACPTTSGGPPYQIFENVAGGDFSSCIGFDARTTEYSGGDTSAAWEYA

>*Pyrenochaeta sp.*

MKSTFAITALIATALAGSCPAPPAPSSSPTPDPSGPSYFGVISARSASPVHLQPLTARGGKFYLGGPGPSSYCPVEVVGDACPPGNTTVLAGGYGSLSLGVVVPGGQQVYVAPDGSLSYTVAHSAYIPPGSTVDKFSREAPAEGQTFGYLNFETGFVACPAGEGQGYQVFGQVQGGTYGPDCLGFNALTVAGDKPGAWQY

>*Cryoendolithus antarcticus*

MKLNTLTTLLASTALTQAQYFTLMSIRSASPIHYGQITASGQRLYINRNTASYCPEQVGDACPAGNTTTFAGGSDSLSMGVIVPGGQQVYIDPVCGAVGYTQAHSAAMPQGAIVGGWNVSLGDPYGYLSRDRGLVACPAADGEGGYQVFGQVEGLVLSEECLRFTAFGINATGPGAWQYT

>*Rasamsonia emersonii*

MKIAVFLATLFTTLLPLLASAMPVMKKPHARDEGNGAFTVMALRSGSPIHYLPVQAGGQRFFLGGKPSTYCPTFVAKCPPGNETVFIEGVYLDTEVPGGQQVYVNPQGALAFTQAHSTFTPPGSSFGPFSYTPGQPYGQYSYTGQGANGFMACPDKAENPSSWQVFAAIKNATVPTGNVNDCLDFVAVTVPYNWSIPAWQYV

>*Rhexocercosporidium sp.*

MIAFLLLISALLAATTAQIFTLTTYAPDSPLHNQKIQAAGQAFYLGLDHPTTYCPTIVDPNCPNATDTRLASMSALWVEVPGGQQTFIRSDGSLGYTQAHSASVPSNAYRGGFVNVTISSDCSPDVNVFTWKAPDGSTEGVLACPDVPSYMENLATHQIFARTPQFNLTDCIELEGLSPTIFPQLAPFGAWQYV

>*Rhizodiscina lignyota*

MKTTLAISAILGLFSFSPVQANYFSVISVRSASLIHLQSINANGGSFWIGKTTTSYCPQAEVSDCPPGNDTVFEGGNDTLGLGTVVPAGQQVYIAPSGILSFSEAHTASFPKGSILTGFSLCPGASFGDLSWEYGLVACPTNGTNGPWQIYGQRADITLPETCLGFDALTYNLTEPGAWQYI

>*Rutstroemia sp.*

MFSKTLIATLLASTAVATPIAARTSNNAFKLISFAFGPNSPLQGLDIVAYNGGLWIGKNTTSYCPEGIDCPAGTETSFVYSDNKNVALNVVVPGGQQLYIGTDYAVSYTVAHSADVHGGKAQGWEYLAPGVDANGPVGLLTYPNYGFLACASPEGPGIYSIAATPGGEGTGNCTAIQLGALEYKGDATAWQYQN

>*Setomelanomma holmii*

MKSTLALALAPFLTTALAQSTPSPPPGPNYFGVIAARSASPIHLSTLTARGLKFYLGGGPPATYCAVTACPPGNTTVLAGGDNTLSMAVDVPGGQQVYIAPDGALSYTQAHSAYIPPGSVVDGFSREYPAAGQQFGYLNFETGFVACPTANQTANGWQVFGQVDGATFGPECLGFDALTIATDTISAWQYA

>*Sordaria macrospora*

MRLAMEKKRLTKASERLPEIAATDVSLSVKRCHIIRYEMFHQEILPIGHVAAQMDPQMRCDIKCRVASLCFVSDGGRTEIIIGISLIDLNMTRLTLLGAALLALGDSAVNALNSCGQSQYDPAQYVCWSNQFLCPITAGEPLSFCNGACYSKFMYKCENNVLSLLPAHTGTPFTLTVSNPTLPIDGKPVTASGLHLSLAGNTSTYCPVEVVGAACPPGNITAFFAGNGGLSMDTMVPGGQQAYLGPDWNMHYTQAHSAYIPAGSLTQGFGAYEGGGFVNLNGNGWGWVACPPQASGGGGSAWNLVGRNETNAASLTACTPVNLKINPLPSGTVGAWQYA

>*Sporothrix brasiliensis*

MTKTLGLALAGLLAAAGSARADLQTCGDAQYDPSQYVCDDGNFLCPIVAGEGLSYCSGACYSKFMYTCTNNALTQLTPLAAGTPFTLTVSNPTAAAVDGLAVDACNLRWNLGIETCSYCPTTGVECPVGNTTVVSAPDTAAGSSTPAMDVVVPGGQAVYLDPSGAVGYTQAHSAQMPGGSITGGLVPYSGGGFVNLNSGGYGWAACPPTAAGGGGTGYTLFAVNADNKATLAGCTPVNLKVNPQPSGTIGAWQYT

>*Phaeosphaeriaceae sp.*

MKYTAALALASTAFAATECTSAPASSSTAAPLPGAPSYFGVISARSASPIHLQSLTARGTKFYLGGGAPSSYCPVQQVGDACPPGNTTVLAGGDKTLSLGVIVPGGQQVYVAPDGALSYTIAHSAAIPSGSIQDEFSKEAPSNGNSFGYLNFPTGWVACP

>*Stemphylium lycopersici*

MKYTTALTTLLAGAMASSCTPSATDSYPSATPSATPESPYFGVISARSASPIHLQPLTARGGYFYLGGEGGPSSYCPVETVGAAACPPGNTTVLAGGDKTLSMGVVVPGGQQVYVGPNGALSYTVAHSAYIPAGSIVDAFSKEAPSNGNSFGYLNFETGFIACPAGEGKGYQVFGAITNGTFADDCLGFSALTSSVDQAGAWQY

>*Talaromyces amestolkiae*

MKFISIITPAVLAASAMALPTSQKRDGSTGPFSVIALRSASPIHLQPVNAAGERFWIGGQAATYCPSEVVDPCPPGTETVWANTNALDVEVPGGQEVYVDPSGALAFTQAHSANIPTGASLGPFTFTPGTQFGTYSTSAFGATGFMACPDDAANPTKWQVFAAFQGASVPTGNVGDCLGFDAGAPAYTGDIPAWQYI

>*Thermothelomyces thermophilus*

MHRFSLVATALLGLTKGVLGDTETCGQVPYDPSKYVCWDNEFLCPITAGEPLSYCAGACYSKFMYTCTNNVLTLLPPVETPFTLTADNPNLPIHGKPVTAANQHWVVNGETASYCPDQVGDACPPGNETVIVSGGGTVSMSVMVPGGQQAYLDPYWNMGYTQAHSAYIPPGSITTGFGAYQGGGFVNLNGNGWGWAACPPRASGGGGPAWNLVARNDTNAERYKDCSPINLKIHSLPSGTVGAWQYT

>*Thozetella sp.*

MSRSSLMAMVTAVVGLGRLVLGAQCGDAFYDPTQYVCWNNNFLCPVIAGEGLSPCSGACYSKFMYTCTNNAVTLLPAVASNTPFTLTAWNPTLPIHGAPVTAAGLRLVINGTTASYCPSQVGSACPPGTITALVASGGAAAMDVEVPGGQAVYLDPYWDVGYTQAHSASIPSGSLTSGFGAYQGGGFVNLNGNGWGWVACPPTAAGGGGPAWNLVGKNDTNAARLNGCTAVNLQINTLPTGTIGAWQYT

>*Trichodelitschia bisporula*

MLRFLFLCFPFANAATLFTAVAASGNTDIHMAPLAASAQHLWLHRDTSAYCPNEPDQCPPGNLTAFGMADSLEGTLSMDVAVPGGQRVFVNQGGEIKYTLPHSAAVDTNSSISGWQLTPSVMSGFGKLSWAPETGGASGGFVACPGESGSGEWTVLAMMGDDGTSFVTAGCSLMEVLVPMTADGGYGIDGPAAWEYE

>*Trichophyton benhamiae*

MKASLITAFVLPLLALASPYLPTDPPANIEVTARHPGRENVDKLPMQAAGRAFWLGGSPATYCPEIVGDNCPPGNATVILGLNSMSVLVPGGQQMYVEPSGKLGFTQAHSAAIPPGSYVGGFAYKPLNKKSGSFYFGGWGATALMACPVPDSKYYQVFANIKNAMVPGGDVKKCVEFVGVAKEYKGTTPAAWQYT

>*Venustampulla echinocandica*

MKSIIAVSALVAMVAAQLEPPAGPFTAGAWNPSNRWTGVAINASGRGFWIGKDSSSYCPVVKGLDCTAFPGTSTVFAGGSGTLSLDVAVPGGQQVFVGPDGALGYTGAHSASIPVGAVTTGFLRFQSVAGGAPVPMTFEGSGFRACPVDETQENVYQVFASAVGTDPGNCIQFQLRTYSAPGISAWQY

>*Zymoseptoria brevis*

MFFVIVLGLTLLPSTFSLLIPPAAANCTTSKCTSKCGDIPFNPAWYSCYTTNNGSALCPIVANIPLDLCNHDCYTPLQYSCNNLTLNPLPRYTGPVSLTALNPAQPFHNQPVQAAGQRFVIGGTAATYCPSSLPEGVCPNLTTTVLYGSGTGLATLVPGGQSLYIRPDGSIGFSQAHSIFVPAGSYQGAIGTAYSNAGFWGVNGSSWYACEADGKGEGWYIYSGLFTKSWEGTQCQEVMLKVENLPMGTLGAWQYI

(2)Amino acid sequences of 53 ThBP24k homologous proteins

Amino acid sequences used to construct WebLogo are shown in red.

>*Amorphotheca resinae*

MLLSPINFIGAAAILSSLASAAPTAAPVYPPKTTSQNFQLIANVTANDLTPSIQNYVVASYHTGAGEAYAVLMPDLSSGRIFYVNGTASDVRFGNSNILSDEGLNQATPPFPAGLAIDSANAVSINAGAGQPGIVLTQFPDPITELAVGGGDGFYACESQLVYGNAVQLFSKGSGETPDGCADVTLLPQCSEDSGAAHPYGQTVNCYADVAGIDWSVYSSD

>*Amylocarpus encephaloides*

MISYFTAIFALTISLADTAPTSSPVFPPRSTSESFRLVANVTSGDLSPSIKDYVLTSYHIGAGQAYAVLTPNISSTTGRIFYQNGTAEDIRYRRSNVLSDGGEPLFPFGIVLNEDQSVSVNAGTGTTGVNLALFPDPISGLSAVGKEGFYACEKTLLYGPAVQVFAKAYEEVTPEGCADVKLLAQCSDSSGAEHPYGATSGCYADVAGIDWSVYSA

>*Astrocystis sublimbata*

MFRTALATAAGLTSLASATPLAARQIVPNYPPNELSKGFRLVANVTDPSTDLSPSVHGQELLGIHIGAGLNEAVLTAQEGDGYVFYHNGTAAEVRAVQGSIISDSGNPPFPLGLTVQGEDEFDSAYPKEHFVHINGGAGTTGVSLLMQPDPYFYVTNFAQGTGTFVACPRKVPYYDSDFIVVRFAYDVWDSENAVSVQDVPENCTAITLIPECDLLSELPEGSLSSHAEARDSKCYVDVSAINWPEYGP

>*Bisporella sp.*

MVSSIFKAASALALLSTASAAPTTRQVPNYPPTSLSSNFRLVANVTGADLSTPINDYVLTSYHTGAGTGYAVLVPATTPTEGRIFYVNGTAEEVRYNNANLLSDSGVQLFPSGVSINADSTISINAGAGQAGVGLKRFPVPVPELAYLGNGEFYACVKDLVFGPAVQLFYKSGTDVATPAGCADVSLLPQCSEGSGAEDEFANTVNCYADVAGIDWSVYSAA

>*Botryotinia convoluta*

MLFSTTNILSTLALLTSVTSAAPTRRQTYSTPPKSTSNHFTLVANVTSGDLAINHFVLESYHTGAGTAYAVLAEDTTEANPRIFYINGTATDVRFGNSSTITDAGSGSDSFPEFLTITPSDPASSSSGRSTVEINAGAGQPYVGLTASPFQTTQLHGYQGETFYACNTTLLYGPAIQLFYRGYSQETPAGCSDVALLPQCSEGSGAEDASAATVSCYDDVAAIDWSVYSA

>*Botrytis aclada*

MLFSTTNILSTLALLTSVTSAAPTRRQTYSTPPKSTSNHFTLVANVTSGDLAINHFVLESYHTGAGTAYAVLAENNTEANPRIFYINGTATNVRFGNSSTITDSGSGSNSFPESLTITPSDPASSSSGRSTVEINAGAGQPYVGLTASPRQTTQLHGYQGETFYACNTTLQYGPAIQLFYRGYSQETPAGCSDVALLPQCSEGSRAEDASAATVSCYDDVAAIDWSVYSA

>*Chaetomium sp.*

MRFTIPALTLAAAASALPSQQPRATSSTSKGFNLVAHVTSGTGNLAQSVEGLFLSTAHTGAGQNAAVFYDTSGRLFYQNGTAASGQSSIVTDGGTPLSIPFSLAVQPAGEPRRILDINVGGAGTPNFISDRGTLVNAGGDGTYLACDAVVPYYNRVFTTLQFVYGKADPAALEGCVAIELVAKCAELNDLPDEGSLASHEFAVDVKCEA

>*Cordyceps militaris*

MIAPIAALGLAGLAAAAPTTGPLYPATSSSNGFNLVLNVTDPSKDLSPPVHGTFVTSIHVGPAYNYVGQSATAAAARLFYVNGTATEVRYGSSDTVSDGGTPPFPEVLSLVQDKASETLSTLWLNAGNGGDANYRTGLAGFPNPIPALYPSTYVACREALAYYGGKEFVIIKQAQTTVGEDGTIDYNIPEGCAPLSLLPQCAKLNDLPAGSLSSHEYAANTRCYENVNSIDWPKYHAY

>*Daldinia childiae*

MLASIITPLTFLALVGATPVPQAAQQLSRSKGFHLKAELRDQTKDLDPSVAGTYLSGIHVGAGQNIAVVSASNPSSSTPYYSNGTEAQGWVDVLNDLGTDFPWGFIVQDSESTDATYPGEHDASINVGGGSDGLTIEKSGDSVALGGLAPGYYAVCDRFIGYVRANIQVVRYVYEGETLPENCAAVNFVPQCATLNDLPATSEWTHDFVQEVDCVVPSA

>*Diaporthaceae sp.*

MKSAILALSMTTLAVAAPAVRRQAPHYPPTSSSQAFKLIANVTDPTLDFSPSIKNWLVQGYHTGAGTNAAVLAPFDAATNPGRIFYENGTAQDQAYHSTSLLTDAGSLPIPFSWQVQSPTEFDATYPTQHHVSIGVGESTPAYIASFPVVYPIVENGLGLGTFVACKNELAGNPSLGEVVTVNYAYATFNGPDENTYEYHANIPEGCVPINLIPECAALDESSAASHEFALVERCYEDVSAINWPEYGP

>*Eutypa lata*

MLNINVLTFGLLGLASAVPLSTRQIPYYPPTSKSLGFQLIVNVTDPSKDLTPPVNGWAFSAIHTGAGFNEAVISENVTASRIFYQNGTAEEIRYREGSVLSDAGTPAYPWGIQVESKDDTAGVEASVSVNVGGGTPAVLLTQFPEPLSYLTGTNPGTYAICPRNVPYYNTVYNVVRWTYDAYNNATGLYEHTVPEDCAEVKLLPQCAELPDLLEWDSSSHDFAATSNCYDDVSAINWAQ

>*Fusarium acutatu*

MLTSTFFLSLLGLAAASPVQPHINVNPKYPSRSTSKGFKLVVNVTDHSKDFDPPIHNTELTSIHTGAGLALVGVIEKDGRIFYQNGTKEERQNGEATVISDGGTPLTPNGIALVADKKNKNLFGATLNFEPGTPGVQLNQPEDPYTYLLPETFVACNKSLEYYQGKHFIVIEQAHLTIDKNNKIQRNIPKGCAPIRLVPQCAELETLPKGSYSSHKYALESKCYKDVSKIQWEKYSP

>*Gaeumannomyces tritici*

MHATFFSLGLAALATAAPTLTLRQAPHYPPSSLSKGFRLIANVTDLSRDLTPSVHGFALGGIHIGPPNSRSVLSPQADNTSRVFYLNGTAADLSLNAARVVSDAGTPPFPWGVRVQSRDEFDVPANPGSHATFINGGSSADVGVTRFPDPYSVLANRKDGPGGTFVACYHEVPYYRRPFVVVDYAYSTVDPATQLPVVEVPEGCAPVTLIPQCAELNDLPADAVSSHEFALDQKCYEDVAAIDWSQYGP

>*Glarea lozoyensis*

MFFTLATAIAILGATSQAAPTSSPIFPPRSSAADFRLVANVTANDLTPSINDYVLTSYHTGAGQAFAVLVPNTSATSGRTFYTNGTAEDIRYRRSNILTDGGSPPFPFGVVINGDESVSINAGTGTAGVNLALFPDPISKLSAVGKAGFYACEKTLPAGPAVQIFTKAFEETTPVGCADVSLLAQCAESTGVEHPFAAVSGCYADVAGIDWSIYSA

>*Hirsutella minnesotensis*

MITAAIICLLGLAAAKPLAADATAKPRYPQASKSRGFHLVVNVTDRSRDFSPPIQNTYVASIHTGAGLALVGNVRGTDNARIFYQNGTAEERRSSTTTVITDSGTPSFPSGVQLVKDKDSDFTSTAHLDAGPGTRGVRISRSPESYAYLLPETWLACNESLPYYQGKYFIIFKQSNSTVDGDGNITQNIPDRCAPVRLIPQCTKLNDLAKGSYSSHLYAIEDRCYKDVKSIKWSDFGP

>*Hymenoscyphus albidus*

MFLSAATAFALIGAFASAAPTSSPIFPPRSTSESFRLIANVTANDLTPSINGWELSSYHTGAGQGYAVLTNPTPSVGRIFYTNGTAEDIRYRQSDILSDGGTPLFPFGIVLNQDKSVSVNAGLGTPGVNLALFPDPISALSTVGKDGFHACQTALPFSPVQIYTRAYEEETPAGCADVRLLAQCSESSGAEHPFGAISGCYADVKGIDWSIYSA

>*Hypomontagnella monticulosa*

MLASITTLSLLALVGATPVPQTSTQLSRAKAYQLQVELRDSSKDLDPPVAGQFLGGIHVGAGLNIAVVEAPPTPMSNPPYYTNGTAADGWVAVLNDLGTNYPWGLNVQDPESFDYTYAGEHDADINVGGGSNGLTVVRDGDSAILGGLAPGNYAVCNRFIGYTRSNMTVVRYVYDGEDLPTDCVAVNFVPLCAQLQDLPEGSEWTHDFVQEVDCVVPPATS

>*Hypoxylon sp.*

MLGSITTGLTLLALAGATPVPQAASSSQLSRSKGYDLRIQLLDPAKDLDPPVDGQYLSTYHVGAGLNVAVAGAPRDPSYSFPPYYSNGTAEQGWVSVLNDLGTSFPWGVSAQNATSFDVTYPGEHDVDINVGGGSNGIHVAKTGDSVALAGLAPGNFAVCNRFIGYARSNLTVVKYVYEGETLPEDCVGVQFTAYCSELEDLPEGSEWTHDFVQEVDCVVPA

>*Lachnellula occidentalis*

MVFIKAATAAALLSTLASAAPTATTPQYPQQTSSESFTLIANVTSGDLSPSVQNWSVTSYHTGAGTAYAVLQADTPRIFYANGTAQDLAFNGGTVLSDEGTPATIPAAIVLGGGDSVTDSISINDGKGTAGVGITRGPDPIAIFYAAGGAGFYACEETFAPGAAKSVMLFQKHDVTPAGCADVVLLPQCSAGSGAVHANPALSGCYADVAGIDWSMYYSS

>*Magnaporthe oryzae*

MLTKSILLATLAALAAANPIARRSGPETSRSRGWKLIANVTESPITGFGAEINHFQVTGIHIGPPYNRAVLYEQYEGGGSRIFYTNGSTEQGGTSQVLTDSNGFSPHFHMQRPDEFDLPANDKSHATFLQPGPDAGDDVVLEGKKLVSRFGAGKFVACKHWVQYYQSLWNVVDYVYEGEETPEGCVDINLLAQCDELKPIEENTEWNWFSHAHVVEEECYVDVAGADWSL

>*Magnaporthe poae*

MQTTLLTLGLAALATAAPAVTPRQTVPHYPPSSVSKGFRLISNVTDPTRDLTPSVHGFALGGIHIGPPNSRSVLSPQADNTSRLFYLNGTASDLTLGTTRIVSDGGTPPFPWGVHVQGPDEFDLPANPGSHATFINGGSTLDVGITKFPDPYSVLVNRKAEGGSAGGTFVACYHEVPYYRRPFVVVDYAYATVDPDTALPVVKVPEGCAPITLIPQCAVLNDLPPDAISSHEFALDQKCYEDVASIDWKQYGP

>*Metarhizium acridum*

MMYTAALVLASLAAASPMASQPGSDYPEKSSSQGFHLVVNNTDLSRELPDPVHHMFLSSIHIGPPSNLLGTSSTAETPRVFYQNATEDEFRSGHGHILSDGGSPRISYGLSSRLEENVADAFVFRLDAGPGQHGVAISPGHHPYAFLKVPGLALCNEPVKYHNGKRMNILKQFPNPTGTLWKMPEECVPVRLLPQCTQLEDAQQGAMSSHEFVQTTHCYKNVSAINWSTYTPFDQIHFNSVV

>*Moelleriella libera*

MHFTRLAAIAFASLGAAKPLAIRDDGPPTALPPSSRSQAFNLIVNVTRLSRDFSPSIHTKYVNSIHVGAGLALLGIGNVTNNPRVFYVNGTTSEFLYAASTVISDSGTPPAPYGITLQKDADSQIASTAHLDAGSGGKGIGLTRFSVPYVYLYPETYAVCNESIPYYGGRSVPIIKQFSLGVDMSSDVIPENCVPVRLIAQCAKLNDLPPNAIANHEFAYETQCYDDVASIDWPKYRPW

>*Monilinia fructicola*

MLFSTSSILSTLALLSTLTSAAPTSRRQTPNYPPQSKSKHFKLVANVTSGDLAINNFILESYHTGAGTAYAVLAENTTEANPRTFYVNGTATDVRFGNSSTITDAGSGNNTFPESIVISPSDPASSPSNRSTVEINAGAGQPYVGLTTFPDPIIQLHGYQGETFYACNSSLPYGDAIQLFYRGYREVTPTGCSDVALLPQCSEGTGATDKYAAEVSCYADVAGIDWSVYSA

>*Monosporascus cannonballus*

MLSKTLSLALAGAVSAVPLSSRQVPNYPIPSTSLGFRLVANVTDPAKDLKPSVNGQYFNAIHTGAGKNDAVLEADGGRVFYLNGTASEIRYNQGHVLADGGLYSWGIYVQSEDEIGREGKHEVTVNVGSGTQGVGLTRHPEPYSYLLWPAAGTFAACDQFEPYYQKNYTTIRFAYDKWNDETALPERDIPDGCVAIRLIPECATLEALPEGSEASHEFAQPARCYDDVSAIDWTQYGP

>*Nemania sp.*

MSMLKALFAIGAASLASAVPLESRQVVPNYPPNSISTGFRLVANVTDPATDLTPSVDGWELQGIHTGAALNDVVLTEGTGPIFYHNGTASEIRYGLGNVLSDLGSQYPYGIGVDNQTEFDYYYPTEHDVTLNVGQGTIGVSLLAFPNPYFYLTAKAPGTYAACSRNVAYYQRDFIVVRYAYDIFDNETAQFVQDIPEGCTAITLIPECDTLPDLPEGSLASHEFASPSKCYEDVSAINWPEYGP

>*Neopestalotiopsis clavispora*

MLDLLALTVLLHSVALAAPQFARQDAVASKYPPTQSSTGFRLIANVSDPSQDFDPPVHGWILDAIHTGAGLNDAVLFADGDDIGRTYYVNGTADEVAAGRADTLTDGGTPPFPYAVSVADRNTTDADGLHDVSVNAGSGDAGVQLEVASSSYPILRGPGAAMGTYVACDQYVEYYDENYTTIRWVYDVQVDGGSSHAGAAAAVTNSTSNTTLAAPPTAPVWLRTIPDGCVAVTLLPECEVLEDLPEESISSHEFAANSTCYGSVSDIDWSLYGP

>*Oidiodendron maius*

MVSSTLSFASVLALLGVASAAPTATSPIYPPKTTSANFRLVANVTANDLSPSINNFVLSSYHVGAGEAFAVLVENVTATTGRIFYVNGTAIDVRYSTSNILSDEGSLLFPAGVSINPADASGEQTVSINSGEGESGIGMTQFPDPISELRVVGGDGFYACNNALPFGSAVQVLSKAQGQATPAGCADITLLPQCSEGSGLAHPFGQITNCYADVAGIDWTYYSTD

>*Ophiocordyceps australis*

MIAAAPVTLGLVGLSAASAIVRQVTPNYAQTSQSDAFNLVINVTDPARDFTPSVQNTYVASIHVGAGLALVGNVDSTQNARIFYQNGTAEQRRDGHSTVITDSGTPPFPSGLRLTKDDASAKVSTAHLDAGQGTPGIRLSNFPEPYVFLNPETWLACNESLPYYQGQYFIILKQAQTSVGQDGYANRNIPDNCAPVRLIPQCTQLNSLPDGSYSSHDFALQDRCYHDVSSIIWSDHGP

>*Pestalotiopsis fici*

MLFKSLSLASAAGLAVASPVSRRDAYPPLSTSTFFTLVANVTDTTKNIFDPPVNGWSLSGVHSGAGLNAAILSANGGATFFVNGTGQEVSSASTSVALPPIASTDGNGQPYYTPQGLQFSDSTTSDEVYIGLNFGLGTKGAGITPGLRSPYADLFGPFGSTLVVCNETAPAYTRPQYPVRGETVVPDNCVAIKLLAQCASGPTLEGVEELNIITEDVKCYEDVSAIDWSQY

>*Phaeoacremonium minimum*

MYTPLIALSLAVLTTAAPSGKRQVPHYPPTSQSQGFKLIANVTDPSADFTPSIKNWVVEGAHTGAGTNVAVLKPFDATTSPGRTFYLNGTAEDVRYGNTNVLTDGGTPPAPYGIYVQNQTEFEYLYPDEHGVGISVGEGSGTSLTRFPNPYPYLTSATGTGTFVACNNTVAYYGTKFITLEYAYATFTAPDGSYEYNANIPEGCAAVNLIPQCTELNELPEDAYSSHEFAAQTACYPDVAAIDWTQYGP

>*Plectosphaerella cucumerina*

MHSAMNIILAGSLAGLAIAGTTTTINQNLDKSAGFELVAVVTNPATDLTPSINFWTLSTIHDGAGRAVAGLSNDAASVWYLNGTENQIKDYESSLLTDAGTPPFPNGLDFVYTPSGTADLRVDAGPGTPGVGLNQNRDLFYLSAPGNDGTYMACRESIPYYSGKEFIVLKYAYHGVPQGCAEIRLVPQCAELAPLPPNALASHEYAMPVPCHVSVEAWETSA

>*Pochonia chlamydosporia*

MRSILLTLIMSLASASPITPAYPETTTSKGFRLVVNVTDPSRDFNPPIHNTYITGSHVGTLNDAVIPNEDKKHARIFFVNGTQDEANADKATTIWADWDTYRSWKLNLESPSQTLSGAFMSGGEGTVGAGIRHYKSPYAFLTPETYVACRQVLPIGGEQVVIKQAKTTYPTNGSPNYNIPKDCAPVRLLSECAELLPLSPDARFNYRFAVESQCYKDVRGIDWTKYM

>*Podospora anserina*

MQTIQLTIITDQTDTILPSNLSDKPPRIATMHSYLLLASLLGLASAAPTSSTDIPRQWTPQEPGSIPRHSGARGFRLRVNVTDPSLDLTPPVHNLFLSTAHIGPAQNRAIVSYSGPIFYQNGSYSDIANYRAGILTDAGPPESPFPEAIQYQQGSDDTKGSELFINAGTPGAGTAISKLTMPYSTLQILEGAPIVKSFIVCGNTTIPYYGESRKFEVVNWLGATRDSTGTHLRVPEGCVAVNLVPECAYEFVDLPEGAAYDRTFVNDVRCYESVAAIDWSKYAY

>*Pseudogymnoascus sp.*

MHYTAATAILAFASAAVAAPQLDKPISPPWTQSTNFRLVANVTGADLVPSIQDYVLTSYHVGAGQGAAVLVPNDATNPGRQFFVNGTAEDIRYNRGTVQSSGGAAPNAYPYGIQIAPAPATSVSINAGLGTTSVGLERFPSPVTYLTAPEAATYVACNQQLPFSEAIALNVLRTGEAVPAGCAQLRLLPQCSAGDGSVHETENIVQCYVDVAGIDWSLYID

>*Pseudomassariella vexata*

MFNSIISLGLFGLASAVPLTSRQLPNYPPSSSSTGFKLVVNVTDPAHDLSPSVNGLFLQAIHTGAGLNDAVLQVDGRIFYQNGTADEIRYNQGNILTDGGNPPFPSGIDVQNKDTTDADGKHEVAINGGAGTKGVSLALFPVPYTILTFIARGTFVACDQFVPYYKQNFITIRYAYDIINPLTSQFEHDIPQGCVAINLIPQCDVLPQLPVPAIASHQFAANVKCYPDVRAIDWTKYGP

>*Rosellinia necatrix*

MAVMKTLLAVGAASLAAAVPLESRQVVPHYPPNKVSTGFRLVANVTDPATDLTPSVNGWELHGIHTGAGFNDAVLTADAGRVFYQNGTAAEVRFAQGSILSDGGEPPFPWGIYVQKEDEHDSTYPAEHDVNINVGSGTIGVSLLAFPNPYFYLTAMAQGTFVACPRTVPYYNQEFIVVRFAYDTSDPVTALPVHHVPEGCTAITLIPECDTLADLPEGSISSHEFARDAKCYEDVSAINWPEYGP

>*Rutstroemia sp.*

MLFSTPTILSTLTLLAPLTSALPTIRQASSSPPTTTSKHFQLVANVTSGSLPIQNFVLESYHTGAGTAYAVLAANTSTANPRIFYINGTASETRFGLSSTLSDAGSGNYSFPEAASNASSVYINAGAGQPYVGLTTFPAPIIQLHGYQGETFYACNNTKLPYGAAYQLYYRGYSQKTPKGCSDVALLPQCSQGAVGVVDKFANEVSCYENVAGIDWSKYSY

>*Sclerotinia borealis*

MLFSTTNFLSTLALLTSFASAAPTIKRQTYSYPPQSTSKNFNLVANVTSGNLAINNFVVESYHTGAGTAFAVLAENTTIANPRIFYVNGTATDVRFGNSSTISDTGSGIDTYPQAIIISPSDPALSSSPNGSVVRMDIAAGQPYVGLTTFPDPIIQLHGYQGETFYACNNTLLYGNAFQLFYRGYSQVTPAGCSDVALLPQCSEARGAVSEFAATVGCYADVAGIDWSLYSA

>*Stachybotrys elegans*

MISSLLPISLLAGLAAASPLAARQVVPNYPPTSTSKGFHLVVNVTDPSLDFEPPINNFYVSSIHVGAGQNLVGVSAEPGRIFYQNGTIEEVQFAQSTVISDGGTPPAPYGFSLQPDAGSEVVSTARLDGGAGTPGVVLSRFPEPYRFLLPGTYLACNESIEYYQGQNFVIIKQADVTVDEDGTINYNVPDNCIAVRLLPECTELNELPSDAFSSHEFAANSQCYEDVSALVWSDYGP

>*Stromatinia cepivora*

MPSSTTNIILSTIALLSSFTSAAPTIKRQTYTYPPQSTSKNFKLVANVTSGDLAINNFVLESYHTGAGRAYAVLAENTTEANPRIFYINGTATEVRFGTSSTISDSGSDDFSFPEAVIITPSDPATSSSGRSVVEINAGAGQPYVGLTTFPAPIIEFHGYQGETFYACNTTLQYGNAIQLFYRGYSQETPAGCSDVTLLPQCSEGSGADDEFAATVNCYPDVAGIDWSIYSA

>*Thermothelomyces thermophilus*

MVRIAIPTLALAATTVTALSKSKGFNLVAHVADPGSDLSPSVEGLLLTTAHTGAGFNAAVFADADAGSERVFYQNGTAARPGGMRSKSTLLTDGGTPPFPFSMQVQSGGEPRRIVDISVGQGTSHYVDARGRLVNAEDAGRRGSWLACNAVVPYYNETFTTLQYAYDYHRGDSPAGLEGCAAIELVARCAELNDLPDDAYSTHEFAVEVRCEK

>*Thyridium curvatum*

MHSAKSILSLALLHAAALTSASPLSLLAARDTSKGFTLIAKVTDPACELDPPVAGWQLDTAHTGAGLNAAVLSDPGQDGGPRIWYLNGTAPPAQQVLTDGGTPLYPYGLSLQAADSPGEHGAVVNVGQSSPTTVKGGRLVNLEGPDGTFLACKRELEYYHSEFVVLQYAYAGEAIPDKCAAITLAPRCAELEVLPPDAGSSHEFAQEVECSAK

>*Tolypocladium paradoxum*

MIASALTLGLIGLAAASPVADRPVVPHYPDTSESKAFHLVINVTDRCKDFNPPIQNTYIASIHTGAGLALVGNLADEGYARIFYQNGTAQEQRSGGITTISDSGTPPFPSGFKLVKDPGSETVSTAHLDGGLGSKGVGITTFPEPYAFLMPETYAACNESLPYYHGKHFIIIKQAQLTVDENGGIHHNIPEGCAPVRLLPECTTLNDLPKGSYSSHEFVIEARCYKDVKSIKWSEYGP

>*Torrubiella hemipterigena*

MFVSSALALAALAAASPLGQRSDGPSYPPTSSSKGFRLVANVTDLSRDLSPSVQNQYVTSIHVGAALNLVGVGSKDVSRLFYINGTAGEFMRGEATTVTDGATPPVPFGFTLTPDSADSALSTANLNAGPGNKGDVTTRIPVPYVFLAPEQMAICDEYVAYYRKNMQIVKQLNLNKDKTLPANCAPVRLMAECAELNELPAGSITNHDNVYTGDCYDDVASIEWSKYTAWGN

>*Truncatella angustata*

MLNALISLAFAGLSCAAPLNARQVVPHYPPTQLSTGFRLIANVTDPSKDFNPPVNNWVFNGIHTGAGSNDAVLLPDGTDSGRIFYVNGTAEEVFYGQGSTLTDGGSPPFPFGIYVASEDQTDADGFHDVSVNVGSGNKGVQLTRFPVVYPTLKGVGQGTYVACDQKVPYYNANYITVRWTYDTVDPETFLNVHQIPEGCVAINLVPECATLNDLPEGSLSSHEFASSSVCYGDVSAIDWTQYGP

>*Ustilaginoidea virens*

MHQLLALLLASLTAASPLFSRADLPPSSKSKAFNLVVNVTEPDRDFFPSIHGEYINSIHVGAGQSLLGVGGKSTNPLTFYINGTTLEFHFARSTVVSDIGTPLSPWGVFFTNDEGSDMAHTAHLDGGPGAKGIGLTRPPVPYTFMYPETYAICNETVPYYKKKFLIVKQFDLGLKLTNAIPPNCVPVRLVAQCTKLDPLPEGSFSNHDFAYETSCYKDVAAIDWAKYQPW

>*Verticillium alfalfae*

MRNIAFLMGLASLTSASPLAVRQTSYPQKSTSSGFTLVVNVTNPAADFCESINHYTLSSIHVGAGQALAAISSESTRVWYVNGTQQEISESKGTVVTDGGTPPFPNGIDIAEAEDSVSAVRVDAGDGTKGVQLTSGSEPYAYLTAPIIGSYIVCNESVPYYQGRKFLLLKHAETEVNEEGESESNIPEDCVAIRLVPQCAKLADLPAGAIASHEFVNEVGCYDDVASIDWSK

>*Xylaria acuta*

MAMFKTLFAAVGAASIASAAPLESRQVIPNYPPNELSAGFRLVANVTDPATDLTPSVDGWEFQGIHTGAGLNDAVLASGTGRIFYHNGTAAEIRFGQGSIVSDSGSPPFPEGIYVQGEDQFDATYPAEHNVAINVGSGTIGVSLLSFPDPYFYLTARAPGTFVACPRTVPYYNQEFIVVRFAYDTFDPDTALSVRNVPEGCTAITLIPECDTLAELPEGSLSSHEFVENSKCYKDVSAINWPEYGP

>*Xylariaceae sp.*

MFKPILALGAACLASAVPLESRQVVPHYPPNSLSTGFRLVANVTDPATDLKPSVNGWELTGIHTGAALNEAILANAPGRVFYHNGTASEIRYGQGSILSDGGSPPFPYGIYVQSESEHDATYPAEHDVNINVGSGTIGVSLLAFPNPYFYLTGPNSAQGTFVACPRVVPYYGVEYTVVRFAYDTIDPSTSLPVRNVPKGCTAITLIPECDTLPKLSKCNKNNGGQASHEFAVESKCYKDVSAINWPEYGP

>*Xylariales sp.*

MLNTVLSLGLAGLASAAPLTSRQVPHYPPTALSTAFRLIANVTDPATDFNPPVNNWVFNAIHTGAGFNDAVLLADGADSGRIFYQNGTVEEIRYNDGSVLTDGGTPPFPFGIYVSSADESDSDGLNDVSVNVGSGTKGVELEHFPTVYPTLRGPAPGTYIACNQTVPYYHANYITIRYAYGILNPDTALYDYDIPEGCVAINLVAECATLNDLPEGSQSSHEYASTVNCYADVSAIDWTQYGP

>*Xylariomycetidae sp.*

MVNFLLTTAALTGLASATPLVSRQTPNYPPTSQSQGFQLIANVTDPSTDLSPSVNGFVLSGIHVGAGLSEAVVSADSGLVFYHNGTAQAIKSGSGSILMDGGTPPFPWGISIAAPSTPSNLSGPEHYVGVNAGSGTSGVTFASFPDPYAYLTAPGQQGTFAACARPVEYYGGQVWTVVRWVEDVWDEEAKLYLPLVPQGCTRVNFLPQCAPLPGLPADAIASHEFALSQRCYEDVSAIQWGMYGP

>*Madurella mycetomatis*

MHFTTATALLSIAAFAAAAPVDIDSYPPTVNSKAFRLVVNVTDPSRDLSPPVHGTEIFPYFINSSLIRAGVAQAGAGSVFLQFPDPNPSSPKFAPFGGQLLTEQGGPDNVLGLNQISHPDDTPFPDNDPRYAMSFTPGKGTMGIQLAVGGHYAFVRPSEFSGGLGNFAICRIDLGQRDHHGHERWMQALDSISLSTDENGEFGLHIPEGCTHVRLVAQCDVLKEVPEGSAAWPFHQAAQEVRCYEDVMAIDWSQYEFSL

(3)Amino acid sequences of 48 ThBP15k homologous proteins

Amino acid sequences used to construct WebLogo are shown in red.

>*Astrocystis sublimbata*

MQINAIVSALFAVAASGAAVKSRAVTYEISGFSADCAAHSTQCGYGFRLVPSTDAPDSTGLICGNLVTGPDSLPPLPTTGCFENMDYAYSIAIADGGLTLTVTSALDASTNITGSHTAAADQFTYENYGIATRQIYVGPKNFTIEAKEVAI

>*fungal sp.*

MQLNALVGALFAVAASGTSVKSRDAVLYNISGFTASCTPHSVQCGYGFRLVPSTDAPGSNGTICGNLVNGPDSLPPLPLTGCFENYAYAYSIALADGGIILTVTSALDENTNITGSHTATADQFIFTPNGISNSESYVGPTNFTIETTEVAV

>*Nemania sp.*

MQINTLVGAFFAVAASGAAVKPRDVTYNIFGFSAFCVPHSVQCGYGFRVIPSTAAPGNNGTICGNLIIGPDNLPPLDLTGCFDPPFAYSIAIADGGLTLTVTSPLDEKTNITGTHTATADQFTIYNGGTTMSQTYVGPQNFTMETVEVAI

>*Rosellinia necatrix*

MQVAAILGAVFAVAASGAAVQPRDAILYNISGFSAFCMPHSVQCGYGFAVISSAAPAGANKTICGNLIIGPDNLPPLPLTSCFEPEYSYAVAIANGGLTLSVTSALDANTNITGTHTATADQFIIQNGGTTMSQIYVGPQNFTIETVEVAV

>*Xylariomycetidae sp.*

MMLNVITTSVFAAVVSGAAVSKRDGCAYSVTNFYASCIPHSVECSYEFDVISAIGSAPTGCSSMLMGPDYLPAVGLTGYQNAAYSWSAQVQKGWLALEITTSLNDRVNLTGTHEIPNSQLAIQDNGASRSQRYTGLQNFAVAATGTEA

>*Biscogniauxia sp.*

MMFNAIIGAIFTAATVSGAALRPRQATFFNVTDFHASCIPHSLLCSYEFDVVAQSSMFPTSCSLMLQGPDILPTVDQTACENSAYAWSVATSNGTLALSVTTVFNGRMNLTGTHIIPTGELIMEQNGASISQKYTGPTGFLMNTVGTPRR

>Xylariaceae sp.

MQFNLILATVAAAATAVSGAAVQPRQATFYTVSNFTASCIPHSVMCSYDFDVVAESSMFPTNCSAFVQGPDLLPAIPPTECDNPAYTWSATPEGEGEQDGSGSGSGSGITLRVSTPFNARINITGIYTIPADQLVVEQNGASSSQRYTGPSEFTLPVTGEA

>*Monosporascus cannonballus*

MKASAICSTLVAVLALGAAQIGRQATEYKVSAFAGSCIPHSLYYNYEFDVATTSALESTHCSLMLLGPDLLPPVRPIGCEDAAYSWSVALGNESLALTVMTPLGEETNLTGVHTITKDQLAMLDHGSVVIQYYKGPRNFAIATERTLA

>*Hypomontagnella monticulosa*

MQLSLLATVFAATASGSVLKARQANSFSVSNFSASCIPHSVMCSYAFAVITNPESTAPSGHPDPVQCGLMLQGPDYLPPVNLTGCSEGAPYSWAVAVNQDRSLQLSVTTPHDSHSNYTGVYDLPADELVVEQHGAVMTQRYTGPAAFEVPLGQPTRH

>*Neopestalotiopsis clavispora*

MQLTFATIFAAAVSARTLTQRDADSYAITDFYASCIPHSTLCSYQFEVDAATNCTILLQGPDILPAVALTGCEDPAYSWSVATTDDAGLALSVTTVSTDDPATNVTGVYDITSDELVITNNGASSSQSYVGPLNFTVATA

>*Valsa mali*

MKFAAAALIAATAVSGAALTKKQADSYSVTEFSANCIPHSVSCNYHFEVMASSGASSPVTCDVTLQGPDSLPAVPLSACSSPSYSFSVVKAASGLDLTITTPLGASSNVTGTHHIDAADIASTQSGAVTTQSYTGSPSFTVPASVAQF

>*Daldinia childiae*

MKVSAVLAIATTASAASVQRRQAQTYDIANFAADCIPHSTSCAYNFNVVTEPYFPLDTCSAFVQGPDNLPDIKEGKCNNTAYTWSTTKATDGSIDFAIWYPFNARSNITYCHTIPASQITTQNNGAVSTQHYTGPKNFTASIFDCASS

>*Truncatella angustata*

MKFSAVIALAASASAATLQKKQAQEYPIGNFYADCIPHSTFCSYNFTVTSDPVLPASHCGAFLQGPDQLPEIKDGSCPDNVAYTWSFTKTADGGADFAIWYAFNSRSNITYCHSIPADEIVTENNGAVQTQHYRGPQNFTASIFDCPSTSA

>*Eutypa lata*

MKFAAVAALASLASGATLQKKQATEYPIANFVASCIPHSVMCSYSFEVISDPSLPASHCEAFVQGPDYLPEIKDGSCSDNVAYTWSVDKVADGGLDFKIWYAFNSRSNITLCHGIGADQLTVDDNGSVQDQRYSGPANFSASIFEC

>*Penicillium flavigenum*

MKVSNLIAFAASASAATIQSRQALNLGIGKFHADCIPHSTYCSYDFTVTSDPVLPESHCNAFIQGPDQLPSISEGSCKDNAAYTWSVTDKADGGLDFAIWYPFNSRSNITYCHSIPASQINTKNNGAVQTEHYRGPVNFTAAVFDCPSA

>*Pseudogymnoascus sp.*

MRFPALVVLAASAAAGTIQSRQRQSLSIGEFHADCIPHSSMCSYDFNVTSDPVLPPSHCNAFLQGTPNLPDAVEASCPDNVAYTWSITNKDDGGLDFAIWYPFNSRSNITYCHSIPAAELVVEQNGAAQSEHYRGPAGFEASFLNCPTA

>*Xylariales sp.*

MKFTPVVLLAASASAAHLKRQSTSLAIGTFDADCIPHSTLCSYNFTVVDDPSLPASHCGAFLASNSNLLPNVTDGSCPDNVAYTWTITSTSSGGLDFSIGYPFNSRSNITYCHSIAASELTVQNDGASVSQHYTGAANFTATVENCVVSAL

>*Magnaporthiopsis poae*

MQFLSALLLAGSASAASLQARQYPLLYVSNFTAACIPHSVRCTYSFGLDSQPSLGWKPTACNITLNGPDRLPNVTQAACERTFHFDVNTTAADAGLDLTVWQWPNSRTYQNYCHHIPGDQLIIEDHGAVQTQRYIGPTDFQVTVADCHNS

>*Magnaporthe oryzae*

MQFISTFLALAATASAATLVPRQQAFFVKSFQAACIPHSVRCTYSFNITSDPAVFPEGSHCEITLNGPDKLPAVTQAPCDNSPFKFDVKPQDGGLYLAASQPINSRTNQDYCHKIPADQLVTDNNGAVQTQRYTGPADFQVSIFECDRK

>*Coniochaeta pulveracea*

MKAATALLAVLSAAVSASPVATRQTTTEFDVSSLYASCTPHSSLCYYSLNVTFPPSTTAVECSGTALGYQTLPELKPTACKDPSVSFAYTLTEGGADLTIDWEYQDGFNLTGKHFIPSSEIEWTNTEIPTGQVQAYSGPRDFAITDLTATASV

>*Hypoxylon sp.*

MHISTIFTGLIPTVALGAAIRSTDACGAQFAIYNFTASCIPHSVFCAIDFNVKTSPGAPSVECSFWGPGPDRLPQVQLSGCRDPNVSFSFGGNGHHELTVVSALAPEQNLTGTYAISDSDIVLEDHGSVQTENYEGPSTFLISDLTTISL

>*Coniella lustricola*

MQFSTAVVALASAMTVSADVIFQVSNFTAACIPHSSECSYDFVVIQPGTMETTGVECTDMRVSDGTLPAVADGTCAESSRTWTVTKPLTGGLVLTVSQPVTPSSNTSGTYTIPVTDLVLTQTGASIQQSYDGPTSFALSS

>*Cryphonectria parasitica*

MQFSTAIVVLASAMSASAETLFSVSDFTAACIPHSSQCSYNFKVLQPGTGETTGVTCSAQKTSDGTLPAVTDGTCTDSARTWTISKPAAGGLTLTVSQQVSAASSQSGSYTIPASDLTTSQTGASVQQSYTGPTAFDLSD

>*Cytospora leucostoma*

MQFSTTIIALAAAMGASAGTSTVFKVSNFAASGIPHSSEVSYSFTVIQPGTMETTGVNCSKLLPSNGALPAVTDGTCKDSSRTWTVEKSGDDLILTVSQPASPSSTQDASYTIPASDLEYKTADASSYQAYTGPTSFDLTESSYSA

>*Diaporthe ampelina*

MQFSTGALLALAASVASADLVFHVSDFQAACIAHSSQCSYSFGVVKIGSGETTPVQCSAMLTGSNGELPEVTDGTCTDSSRTWTITKFSTNHASGLELAVTEPVTPSSSLVGRYDLLSGTFEMQQTGATTQQVYVGEETEFDLAN

>*Massarina eburnea*

MQFFIAASLLAATASAASPVFNVSKFAASCAPHSSLCDYSFGVVQASAGESTPVDCSAQVVSDGSLPAVTDGTCKDSSRTWTITKNDDGGFTLTVSQAVTPSTNLKGTFVAPASDISKEDTGASSQEKYIGSTDFGLASA

>*Diaporthaceae sp.*

MQFSAILALAAASVASADVVFKVSDFSASCVPHSAQCIYEFGLLQPGTMQTEPQKCRAQVTGSDGTLPAITEGTCEQTSRTWTVTKADGGLVLEVSQQVTPSSNQTGKHTIPASELEMEQTGASIQQKYTGAKDFDLV

>*Botryotinia calthae*

MQFSSAIISAITVALASAAAIEKRQNVFDVSDFSAGCIPHSTQCSYSFTVIQPGTMATVGVKCTALVSGNVDGTLPDIPQWGGSCLESSRRFWFTREDAGLKFFVYQQITPSSNQTAFHLLANDDFVMVPNTIGSTQSYTGPTSFGLDYGL

>*Botrytis aclada*

MQFSSAVLSAITVALASAAAVKRDSSVFNVSEFSAGCIPHSTHCLYSFTVIQPGTMETTGVKCTALVSGNTDGTLPDIPKWGGSCIDSSRRFWFTREDAGLNFFVSQQVTPASNQTASHLIPNSELVMVPSSIGSTQSYNGPTSFGLDYATSA

>*Sclerotinia borealis*

MQFTTAIISAITMAVASASVIGTQRYVTFKVSDFSAGCVAHSSQCLYSFTVIQPGTMETTGVKCTALVGSIDGTLPNIAQWQGSCGDESSRTFWITREDAGLEFYVSQPVSPASNTTASYLIPNSELVLTPNTIGTTQSYTGPSDLDLYQY

>*Monilinia laxa*

MQFSTAVLSAITVAVASAATIGQRDTVFKVSAFSAGCIPHSTQCLYSFNVIQPGTMETTGVNCSALAPATTSGNLPDLAKWQGKCGESSRTFWLTHKDNGVEFTVSQPVTPSSNQTGSYLLPNSDFTTSQTTIGPVQSYTGPSSFNLS

>*Colletotrichum aenigma*

MQFLTILASAAVASAAAIQQRDVTFKVSEFSAGCIPHSSQCSYSFTVIQPGTMETTGVKCSGLFQAQGTSTLPDVKEAPCTDSSRTFDVVRSAEGLTLTVSQPVTPSSNQTGSHLIPNDQLVVATQPNAQVQSYNGPTAFDLE

>*Ophiostoma piceae*

MMFSAAAILSAVVAVSANPLSARSDTVFDVTEFEAGCIPHSSQCRYSFGVVQANNGETIPVQCVLLATANGGDLPNVTDAPCTDSSRTFSVVRGTEGLTLTVSQPVSPASNETGSHLLAASDLVTATQPNAEVESYEGPTSFPLVR

>*Dactylonectria estremocensis*

MKFFTAAILSAVAVSASPVAVVADPVFTVSDFSAGCIPHSTQCLYSFGVLQPGTMETTPVKCEALVSANTDGTLPNVKHGKCKESSRVFSVTRGAKGLYLKVWQPITPSSNQSGKHLLAKKDLVISNKPNAVVQSYKGPRSSAFTRLGVEKRRDNIHRRIAHSITT

>*Ilyonectria destructans*

MKFFTAAILSAVAVSASPVAVAADPVFTVSDFSAGCIPHSTQCLYSFGVLQPGTMETTPVKCEALVTANTDGTLPNVKHGKCKESSRLFSVTRGAKGLYLKVWQPITPSSNQSGKHLLAKKDLVTSNKPNAVVQSYKGPKKFTLY

>*Fusarium albosuccineum*

MKFFTAAILSAAAVSATPVFTVSDFSAACIPHSTQCSYAFGVLQPGTMEKTPVQCVALVSANTDGTLPNVKEGKCKESSRTFNVVRSKKGLTLTVSQPITPASNQTGKHLIPNKQLVISNKFKAVVQSYKGPKKFNLN

>*Geosmithia morbida*

MKFSIVAVLSAAAAVSALPVFEVSDFSASCIPHSSQCSYAFNVIRPGNGEVTPIKCSSLTTSDGTLPAVKEGTCKDSSRTFSVTRILAGLHLTVSQPVTPSSNLTGSYTIPQSQLVTSTEPNAEVQSYNGPTSFRLD

>*Chaetomium globosum*

MKLSLLSMATAASAAAVTKRDVVFEARNFTASCVPHSAMCHYSLNAFMPGTMDTQGYDCSASAPGAGVAILPEIQGGACPPSSRTFDIVRSEEGLTVIVSVQVSRLSYTRGSHFIPNEQIKMIEGVTPTGNYQAYVGPKDFSLERI

>*Microdochium bolleyi*

MKFFTAITLAASASAATLERKQEPSKAISEFKAACIPHSVRCQYSFTISSGGLPGSHCSAEALGGGNLPAIGDGKCEDNSAYTWGIVSTMQDGGMRFGVTYPLNSRSNITYCHQLPASDFVVDDNGSVQTQRYAGPTEFEIKWADC

>*Podospora anserina*

MKLSLLALASAASAAVITTRDVVFEARNFTASCVPHSVMCFYSLNAFMPGTMDTLGYNCTASGVGGLGILPEIKGGTCPPSSRTFDVLRNATGMTVIVSVQVSRLSYTRGAHFIPSEQIQIIKGVTPTGDYTAYVGPKDFPLERIW

>*Thozetella sp.*

MKLTFILAATGASALAVKRYTVFEVSNFTASCIPHSVQCYYSFGVFQPGTMQTEPQHCSAFLTSAGVGQLPDVKEGKCEETSRTFDIVRGDDGLTLTVSQPVSPISNQTASHLIPADELVYAGEPNGVVESYVGPKDFDLE

>*Periconia macrospinosa*

MKFLTVAATSILAATASAAAILPRDNVFEVSNFYANCVQHSVMCNYQFSVLYPATGETIPVNCSATLSSIDGTLPPVTDGDCLLSSRTFDVARNDDGLLLTVKNPVSPSTNITGVYEIPAEQLVVEQSGASQAEKYVGPKEFGLAYAS

>*Decorospora gaudefroyi*

MHFFTIASLLAATVSGAACGAKKSDEPAAPTFDVENFKAFCLPHGGACYYVFEVYKHRSGEDIPTECNTSVRGDGSLPSINDGQCNSSRTFDVIRNEDGSLNFSVTQQVTPISFTTSNHTIPADQIEVQDGQEVYVGPTSFGLYGY

>*Plenodomus tracheiphilus*

MQFFTTASLFAAAVASAAVMPRQETYTVFDVTNFSAECLSHGGACIYSFNVVQHNNGEFIPTECDNSVRGDGVLPALSNGQCKDSSRTFNVTKNADGTYTLSVTQPVTPSSDEEGSHTITADQIELNEDDQQVYVGPKNFTLQSAPSTISK

>*Grosmannia clavigera*

MKFIAAALFSAAVVSADAIYSVTDFNAGCIPHSVECSYSFSVVHVNNGEFVPVNCTASAVSDNTLPAVTGTCEESSRTFVIARNSTGLTLTVTQPVSVVSYESGSHFIPKDEITTKSDQVATLEIYDGPTAFTLFD

>*Xylaria acuta*

MKYTSAAAFLSLASVANIDARLATDLKYEISDFSAACKPESIYCLYEISIVTSNNPEFKVSCDAIGTSDNGELPAVGKTHCGTYTISVAKSDDGGLILRVKSNIEPLAGTFTIPTDDLTTTTSGESTVQSYTGDSAFTIDIEDASSASASASASALGTTASTATASSASSASSSASSVSPTAASEPSATGTTTSSSTSPSETNGASRESAFAGVPFAVGLMAFIF

>*Clonostachys byssicola*

MKFSTIAFAAGATAAVYKRQDTVFEVTNFSASCIPHSTQCRLLDLVYSVGTVADDCLVSSYDFDVFEPGTMQTTPQHCAAMVSADGIGELPALDNGECEQTSKTFKIIKSFGGLALYVYEPVTPSSNKSGVHQIQTAELEMVVPAENPNGAHQVYKGPTSFPLTPQGM

>*Pestalotiopsis fici*

MKFSAVVALAASASAATLQKKQALDYPIGNFVASCIPHSTYCSYNFTVTSDPSLPASHCGAFIQGPDVLPNVTAGSCPDNVAYTWSATRTEEGGLDFAIWYAFNSRSNITYCHSVAADEILTENHGAVSDERYIGPANFTASVLNCNVA

(4)Amino acid sequences of thaumatin-domains from *O. sativa* and *C. reinhardtii* used for phylogenetic tree reconstruction

>Os03g0663400

NKCGYTVWPAALPSGDGNQLDPGQSWAVYVPAGTKGARVWGRTGCGFISGGSLGQCQTGDCGGTLRCAAVGAPPVTVAEFSLGQASKDDYFDISLVDGFNAPMAIVPAAAGGRRCPRGGPRCAAEITLQCPGELRAKAGCSNPCRGNSTCGPTKDTEFFKKLCPETVTYARDGQGTTFTCPAGTDYQIVFCP

>Os03g0244200

NNCGYTVWPGLLSGAGTAPLSTTGFALAHGASATVDAPASWSGRMWARTLCAEDATGKFTCATGDCGSGGIQCNGGGAAPPATLMEFTLDGSGGMDFFDVSLVDGYNLPMIIVPQGGGAAAPAGSGGGSGGKCMATGCLVDLNGACPADLRVMAASTGTGAAAPGGGPVACRSACEAFGSPQYCCSGAYGNPNTCRPSTYSQFFKNACPRAYSYAYDDSTSTFTCTAGTNYAITFCP

>Os03g0243900

NRCTGTVWPGILSNAGSARMDPTGFELPPGAARAVPAPTGWSGRLWARTGCTQDGTGKVVCATGDCGSGTLECAGRGAAPPATLAEFTLDGGGRNDFYDVSLVDGYNLPLLVEPSGALGATATTCAAAGCAADLNARCPAELRAVGGAACRSACDAFGKPEFCCSGAYANPNTCRPTAYSQVFKSACPRSYSYAYDDPTSTFTCAGGRDYTITFCP

>Os03g0233200

NNCTYTVWPATLSGNTAVAVGGGGFELSPGANVSFPAPAGWSGRLWARTDCAPSGTASLACVTGDCGGAVSCSLGGAPPVTLAEFTLGGTDGKDFYDVSLVDGYNVGIGVAATGARVNRSTCGYAGCVGDVNALCPAELQVAGKENDQQSGAAATTTVACRSACEAFGTAEYCCTGAHGGPDSCGPTRYSRLFKAACPAAYSYAYDDPTSTFTCGTGAQYVITFCP

>Os01g0839900

NNCPYPVWPGIQANSGHDVLEGGGFFLPALSHRSFAAPAHPWSGRIWARTGCTGAGAQLHCATGDCGGRLQCAGLGGAAPATLAQVSLHHGNDQTSYGVSVVDGFNVGLSVTPHEGRGNCPVLACRKNLTETCPSELQLRTPAGSVVACKSGCEAFRTDELCCRNMYNSPRTCRSSKYSEFFKRECPQAFTYAHDSPSLTHECAAPRELKVIFCH

>Os01g0113650

NRCSFTVWPAAVPVGGGMRLDPGESWALDVPANSGAGRVWARTGCSFDANGNGSCQTGDCGGVLKCKNSGKPPQTLAEFTVDQTSVQDFFDISLTDGFNVPMDFLPVPAPEQRHGAPPCSKGPRCPANITSQCPSELKAPGGCNSACNVFKQDKYCCTGTTGTKTCEPTTFSLPFVRMCPDAYSYSLDDSSSTTFTCPSGTNYQIIFCP

>Os01g0113350

NLCPFTVWPAAVPVGGGRRLDPGTSWALDVPAGAAPGRVWARTGCTFDATGANGTCLTGDCGGALSCTGYGEPPQTLAEFSLGQADGQDLFDISLVDGFNVPMDFLPAPPPDQSPPCSKGPRCPANVTAQCPGELRAHGGCNSACRVFKQDKYCCTGNGTNTCEPTTYSLPFVRMCPDAYSYSRNDASSPGFTCPSGTNYQIIFCP

>Os07g0419300

NNCPETVWPAALSSAGRPPFPTTGFALPPGASLSVAGVTATWSGRVWGRHRCATGGGRFSCESGDCGTGQVACNGAGGAPPATLAEFTLGGGGGGAGALSDFYDVSNVDGFNLPVEVRPELLERRQEGAAAALCRTTACPADINRVCPSELAVRAADRGGAAAAAVVACKSACLAFGTDEQCCRGRFASPDKCAPSEYSRLFKAQCPQAYSYAFDDRSSTFTCANATGYRITFCP

>Os07g0417600

NNCHEVLYPGVLTPATAQAFPTTGFELQPGASAAYDGVPDNWSGNIWARRLCSTDASGRFSCESGDCGTGRVECDGRGNGPPSTLSEFTLRGGSARDTDFYDISNVDGFNVPVQVAPSGAGCSAVACAADIDASCPAELAVKGAGGAVVGCKSGCLAFGRDDLCCRGAYGTPDKCPPSQYSKFFKDKCPQAYSYAYDDKSSTFTCTSGASYQITFCP

>Os06g0691200

NRCAETVWPGIQPSAGKELLARGGFQLAPNRATSIRLPAGWSGRVWGRQGCSFDAAGRGRCATGDCGGALYCNGAGGAPPATLAEITLASTPAAQDFYDVSLVDGYNIPIAMTPSHGSGANCVPAGCISDLNRVCPAGLAVRGGGGDNRVVGCRSACAAYGAPQYCCTGQFGSPQQCKPTAYSRLFKTACPKAYSYAYDDPTSILTCSAGASYIVTFCP

>Os10g0524900

GGVSGRERRRRRASAAADGDASALSSSTAPGECGGRLDCGGTGATPPATLFEVTLGKGGGGGGAGDLDYYDVSLVDGYSLPIVAVPQAGGGCATTGCTADLNRSCPKELQVDGGGGTVACRSACEAFGEEEYCCSGAYATPATCRPTAYSAIFNELR

>Os10g0412700

NKCPFPVWPAAAPNAGHPVLAGGGFLLPPGQSRRVSAPPTWNGRFWGRTGCNFTSTATNHHGNAAASCLTGDCGGRLACNGTAGAPPATLVEVDLHEDQSKGSSYDVSLVDGYNLPVAVWTKPPTPGAAAADRKCVIPGCAKNVNAVCPPELQVTAAAAVVACKSACVAFGTDAFCCRGAHGTAETCRGSAYSRVFRDACPAYVSYPYDTAAARCYAEVYVLTFCP

>Os09g0536400

NRCGGTVWPGVLSNSGSSALGTTGFALGAGETRSLAAPAGWSGRFWARTGCTFDDDGKGTCATGDCGSGEVECRGAGATPPATLVEFTLGSGGGGGKDYYDVSLVDGYNLPMVVEAAAAGCPATGCVVDLNQRCPAELKAGHGQACRSACEAFGTPEYCCSGDHGNPDTCHPSVYSQMFKRACPRSYSYAYDDATSTFTCTGTDYSITFCP

>Os09g0536300

NSCAYTVWPGLLSSAGSPPLATTGFALAPGESLAVDAPAAWSGRVWGRTLCGADPGGSGRFACATGDCGSGAVECGGGGAAPPATLAEFTLDGAGGNDFYDVSLVDGSNLPMVVVPQGGGAACGATGCLVDLNGPCPADLKVAGADGAGIACRSACEAFGTPEYCCNGAFGTPATCRPSAYSQFFKNACPRAYSYAYDDATSTFTCASGTASYLVVFCP

>Os09g0498300

NYCSHPIWPGTLAGAGTPQLSTTGFRLDPGQTAQLAAPAGWSGRIWARTGCVFDADGAGVCQTGDCGGRVECRGAGAAPPATLFEVTLGRGGGEDFYDVSLVDGYNLPVVAIPRAAAACNATGCMADLNRCKCTHERAPRRRHFAIAGADDDDVCSVRS

>Os08g0517800

NHCAQTIWPATLAGAGTPQLATTGFRLDPGQSVQVPAPAGWSGRIWARTGCDFSGAGGAAAAAAAGAAACQTGDCGGRLECGGTGATPPATLFEVTLGKVGGGAGAGDLDYYDVSLVDGYNLPVVAVPQAGGATGGGGGCATTGCTADLNRCTCRASMHACPGRWRAGARARRSGRRSTAAAARTRRRR

>Os12g0629300

NQCPYTVWPAATPVGGGVQLNPGDTWTIDVPAGTSSGRVWGRTGCSFNGAGTGSCATGDCAGALSCTLSGQKPLTLAEFTLAGSAGGSQQLDFYDVSVIDGFNVGMSFSCSSGETLTCRDSCCPDNTKLRHCNANSNYQVLFCP

>Os12g0628600

NRCSFTVWPAATPVGGGVQLSPGQTWTINVPAGTSSGRVWGRTGCSFDGSGRGSCATGDCAGALSCTLSGQKPLTLAEFTIGGSQDFYDLSVIDGYNVAMSFSCSSGVTVTCRDSRCPDAYLFPEDNTKTHACSGNSNYQVVFCP

>Os12g0569500

NLCPYTVWPIVSPDSGSPPIADGIRLEGRGVGLRSLNLPAGFWSGRVVPRTWCRDGGRCDTGNAPPATVVRLSFNGAGGLAEYSVNLGEGFNVPTVVSPHAIGGGMCPALGCTADLNAGCAAGQRVYGGDTGGDVVACRGPASYFKQRCPLT

>Os12g0569300

NLCPYPVWPLVTPNTGFPSISGNTARLDGGGRGLVSYDFPASFWAGRVVARTGCGGGGGLVRCETGNAPPATVVQLVVHSPEGAQDLAAYSVSLVDGFNVPAVVSPQAIAGGGQCPALGCAADLNAGCPRSQRVVGAGGAVVACRGTADYFKARCP

>Os12g0568900-ThEGL-01

NLCPHPVWPLVTPTSGQPISDNTARLDPNSLISLAFPPTPWSGRVAARTGCDAAASPPAGCETGASPPSTVAQLSVHGGGDVATYSVSLVDGFNVPVVVSPQAVGGGQCPALGCVVDLNCDCPLGQRFSDGAACRGPPE

>Os11g0706600

NNCGESVWPGLLGTAGHPTPQSGGFHLGAGEEAALEVPAGWSGRVWPRRGCSFDSRGRGSCATGDCGGVLRCNGAAGATPATVVEMTLGTSASAMHFYDVSLVDGFNAPVSMAAVGGGVGCGTAACGADVNVCCPSALEVRDREGRVAGCRSACRAMGGDRYCCTGDYASPSACRPTIFSHLFKAICPRAYSYAYDDATSLNRCHAKRYLITFCP

>Os11g0703100

NLCPYTVWPLVTPNAGQPAIITGGATIRLDPNGLASLAFPAAAGWSGRVVPRTGCTGAATCATGDAPPATVAQVSVNAAGGLAEYSVSLVDGFNVPATITPHAFDGSQTCPVLGCAADINAACPADARVGAGCRASP

>Os12g0630500

NRCSYTVWPGALPGGGARLDPGQSWSISVAAGTPAARIWPRTGCSFDGAGRGRCATGDCAGALSCAVSGEPPTTLAEYTLGRPGAGGDDFLDLSLIDGFNVPVSFQPANGGGARCSKGRGPSCAVDITARCLPELRVPGGCASACGKFGGDTYCCRGRFEHVCPPTSYSMFFKGLCPDAYSYAKDDQTSTFTCPAGTNYRVDFCP

>Os12g0630200

NRCRYTVWPGALPGGGVRLDPGKSWTLNVAAGTKAARIWPRTGCDFDGAGRGRCLTGDCRNALSCAVSGAPPTTLAEYTLGTPGAAGGDATDYFDLSLIDGFNVPMSFQPTSNAARCGARRRGPSCGVDITAQCLPELKVAGGCDSACGKFGGDAYCCRGKYEHECPPTKYSKFFKDKCPDAYSYAKDDRSSTFTCPAGTNYQIVMCP

>Os12g0630100

NRCSFTVWPAATPVGGGTQLNPGQTWTINVPAGTSSGRVWGRTGCSFDGAGRGRCATGDCGGALSCRLSGQPPLTLAEFTLGTSGGNRDFYNLSVIDGYNVAMSFSCSSGVTLTCRE

>Os12g0629700

NRCSFTVWPAATPVGGGTQLSPGQTWTINVPAGTSSGRVWGRTGCNFDGAGRGGCATGDCGGALSCSLSGRPPMTLAEFTLGGSQDFYDLSVIDGYNVAMSFSCSSGVGLTCRDS

>Os03g0661600

NRCQYTVWPAAVPSGGGTKLDPGQTWTINVPAGTTGGRVWARTGCGFDGSGNGQCQTGDCGGKLRCTAYGAAPNTLAEFALNQWNNLDFFDISLIDGFNVPMAFLPAGSGAGCPKGGPRCATAITPQCPSELRAPGGCNNACTVFRQDRYCCTGSAANSCGPTNYSEFFKRLCPDAYSYPKDDASSTYTCPAGTNYQVVFCP

>Os03g0663500

NKCQITVWAAAVPSGGGQQLDPGQQWVIDVPAGTTGGRVWARTGCSFDGSGNGRCQTGDCGGVLRCAAYGQPPNTLAEFALNQFSNLDFFDISLIDGFNVPMDFLPAGDGAGCAKGGPRCEADVAGQCPSELRAPGGCNNACTVFKQDQYCCTGSAANNCGPTNYSQFFKGLCPDAYSYPKDDQTSTFTCPAGTNYQVVFCP

>Os12g0629600

ATFTITNRCSFTVWPAATPVGGGRQLSPGDTWTINVPAGTSSGRVWGRTGCSFDGSGRGSCATGDCGGALSCTLSGQPPLTLAEFTIGGSQDFYDLSVIDGYNLPMSFSCSSGVTVTCRDSRCPDAYLFPEDNTKTHACGGNSNYQVVFCP

>Os05g0538500

EAAGTTVFTMRNNCTYTVWATTLSRNTAVAIGAGGFELSPGANVSFLAPDGWSGRLWARTDCATSGTASLACATGDFGGAVSCSLGGAPPVTLAEFTLGGGDGKDFYDVSLVGGGRRGSPGAAPPPAARALLLAAPPRRPPAASPPTAASRHPPPFLPAPSTACGG

>Os11g0703000

NLTLHNLCTHPVWPLVTANAGLPAIADAAGAATRLDGNGDGLATLAFPPGAWSGRVVARTGCRGNGSSRCDTGDAPPVTVAQVSVHGAGGLAEYSVSLVDGFNVAVVVTPHGFEQGRLCPSLGCAVDLAADCPGDGGRGGCMAAGQAEAFKARCPDTRTTPTDVEVTPQRCIQPAELKVVFCP

>*Chlamydomonas*_TLP

IDVRNNCGYAVTAFWRSGAGSTYNSRVLPGQVVRITFSGGVWIGGVIWGSRTGSINNGQATQLEFTIGADMGGGRKQDFYDVSVVNAYNTPARIRPLNPPSVSGSWCGSPTCILPSLTTFCTSPNYLAGADPACINKDGPGLVATSGTKAFKAKCPQAYAYSKDDATSMFACQWGTNYEVKFCP
